# grassp: A graph-based framework unifies subcellular proteomics across assays and studies

**DOI:** 10.64898/2026.11.23.768756

**Authors:** Max Frank, Shivanshi Vaid, Madhura Bhave, Kyle Awayan, Marika Clark, Frank McCarthy, Johnson Truong, Brian C. DeFelice, Leandro Lima, Camille Januel, Rodrigo Baltazar-Nunez, Zachary Yu, Madhuri Vangipuram, Leila Njoya, Laura Savy, Daniel N. Itzhak, Joshua E. Elias, Manuel D. Leonetti, Duo Peng

**Affiliations:** Biohub, San Francisco, California, USA

**Keywords:** subcellular proteomics, protein localization, mass spectrometry, protein-neighborhood graphs, graph-based annotation, data integration, integrated pan-human map, scverse, data portal

## Abstract

Mass spectrometry-based subcellular proteomics is generating a rapidly growing collection of protein localization datasets (“maps”) across diverse biological contexts. This offers unique opportunities to understand how localization shapes cellular function at scale. However, differences in experimental assays, data formats, analytical strategies, and compartment vocabularies hinder systematic comparison and integration. Here we present *grassp*, an open-source computational framework that represents subcellular maps as protein-neighborhood graphs. Using novel graph-based annotation methods, *grassp* enables analysis, comparison, and integration across experimental assays. We used *grassp* to uniformly process 103 human, mouse, and yeast datasets, and to combine 51 unperturbed human datasets into an integrated pan-human map. The integrated map surpassed every individual map in subcellular resolution, separating proteins down to the level of protein complexes. This provides a new data-driven reference for the localization of 9530 human proteins. The full harmonized data compendium can be explored interactively or agentically at https://grassp.apps.biohub.org/.

## 1 Introduction

Eukaryotic cells are partitioned into membrane-bound organelles and membraneless compartments, organizing intracellular proteins into distinct environments that separate interaction partners, chemical contexts, or substrates. Therefore, protein localization has emerged as one of the key determinants of protein function in eukaryotes [1], and as a key driver of disease [2]. Cells also remodel their subcellular organization during differentiation, stress, or infection, frequently without changing protein abundance [1, 3]. This establishes protein localization as a key modality one needs to build a comprehensive model of the cell [4]. For such a model to learn generalizable principles of protein localization, it will require robust, comparable training data spanning diverse cell types, biological conditions, and disease states.

Mass spectrometry-based subcellular proteomics (MS-SP) is uniquely suited to provide such data, because it produces untargeted, proteome-wide measurements of subcellular organization [5, 6]. In recent years, several complementary MS-SP experimental approaches have been developed. Fractionation-based methods infer localization from protein abundance profiles across biochemical fractions, such as density gradients or differential centrifugation (DC) [7–13]. In contrast, affinity-based methods such as proximity-dependent biotinylation and organelle immunoprecipitation (OrgIP) enrich cellular compartments using compartment-specific bait proteins [3, 14]. These approaches differ in throughput, coverage, and subcellular resolution, and produce distinct quantitative measurements (e.g., fractionation profiles vs. bait-specific enrichment scores). Despite these differences, they all generate quantitative profiles for individual proteins that encode information about their subcellular localization.

A central analytical task is to infer protein localization from these profiles, typically by comparing them with reference proteins of known localization. A range of computational tools has been developed to this end. The pRoloc ecosystem [15] provides a comprehensive toolkit for subcellular proteomics analysis written in R, supporting supervised and semi-supervised classification, novelty detection, visualization, and quantitative assessment [16–21]. Other tools support specific experimental designs or analysis tasks, including MetaMass, SubCellBarCode, DOM-ABC, and C-COMPASS [3, 13, 22–25].

We refer to the protein profiles measured in a single experiment under one condition as a dataset, and to the representation and annotation of protein localization computed from it as a *subcellular map*. Three barriers hinder the development of a shared, cumulative MS-SP resource from individual maps. First, experimental platforms and data-processing pipelines produce incompatible measurement data and software objects. Second, studies annotate localization labels using different reference marker sets and compartment ontologies. As a result, differences in annotations may reflect these ontology choices rather than the underlying measurements, complicating comparative analysis. Third, raw data, processed maps, metadata, and annotation results are currently not stored in a unified format or a central database that is shared and accessible throughout the field. To our knowledge, no existing framework attempts to overcome these barriers through a unified workflow for data ingestion, annotation, integration, and exploration while maintaining compatibility with specialized analysis tools.

Here, we present *grassp*, an open-source framework that makes subcellular maps from any assay comparable, combinable, and shareable. To unify quantitative results across multiple experimental assay types, *grassp* represents each subcellular proteomics dataset as a protein-neighborhood graph and introduces robust graph-based annotation methods using either curated marker proteins or hierarchical compartment ontologies. All data and analyses are stored in a unified AnnData format, a common data structure in single-cell genomics [26, 27] that is gaining adoption in MS-based proteomics [28, 29]. *grassp* is written in Python, while the *grasspio* R package also makes these maps accessible through R/Bioconductor and its repertoire of community-supported tools.

We used *grassp* to uniformly process and annotate 103 human, mouse, and yeast datasets, creating a standardized collection available through the *grassp* web portal for discovery and cross-study analysis (https://grassp.apps.biohub.org/). We further combined 51 unperturbed human maps into an integrated pan-human map that shows improved resolution of subcellular compartments and protein complexes relative to individual maps in our benchmarks. This integrated map provides a new reference for human protein localization, bringing together complementary evidence across studies and experimental systems. Just as the scverse infrastructure and community portals have enabled the training of large-scale gene expression models, we expect *grassp* to provide a foundation for models of protein localization as additional maps become available.

## 2 Results

### 2.1 The *grassp* framework enables data ingestion, analysis, interoperability, and exploration

The *grassp* framework consists of three software packages and two web applications organized around the AnnData [26] format as the common handoff between data ingestion, analysis, and exploration (Fig. 1). The search-engine-agnostic *protdata* package converts quantitative proteomics outputs into AnnData objects. Because all down-stream steps operate on standard AnnData objects, other converters, such as the recently published AlphaPeptTools [28], can be used in its place. Each AnnData object stores the protein-by-feature quantitative matrix together with protein and sample metadata. It can also store transformed measurements, protein-neighborhood graphs, embedding coordinates, compartment annotations, and analysis parameters, allowing intermediate and final results to travel with the underlying data rather than being distributed as disconnected files.

**Fig. 1.**
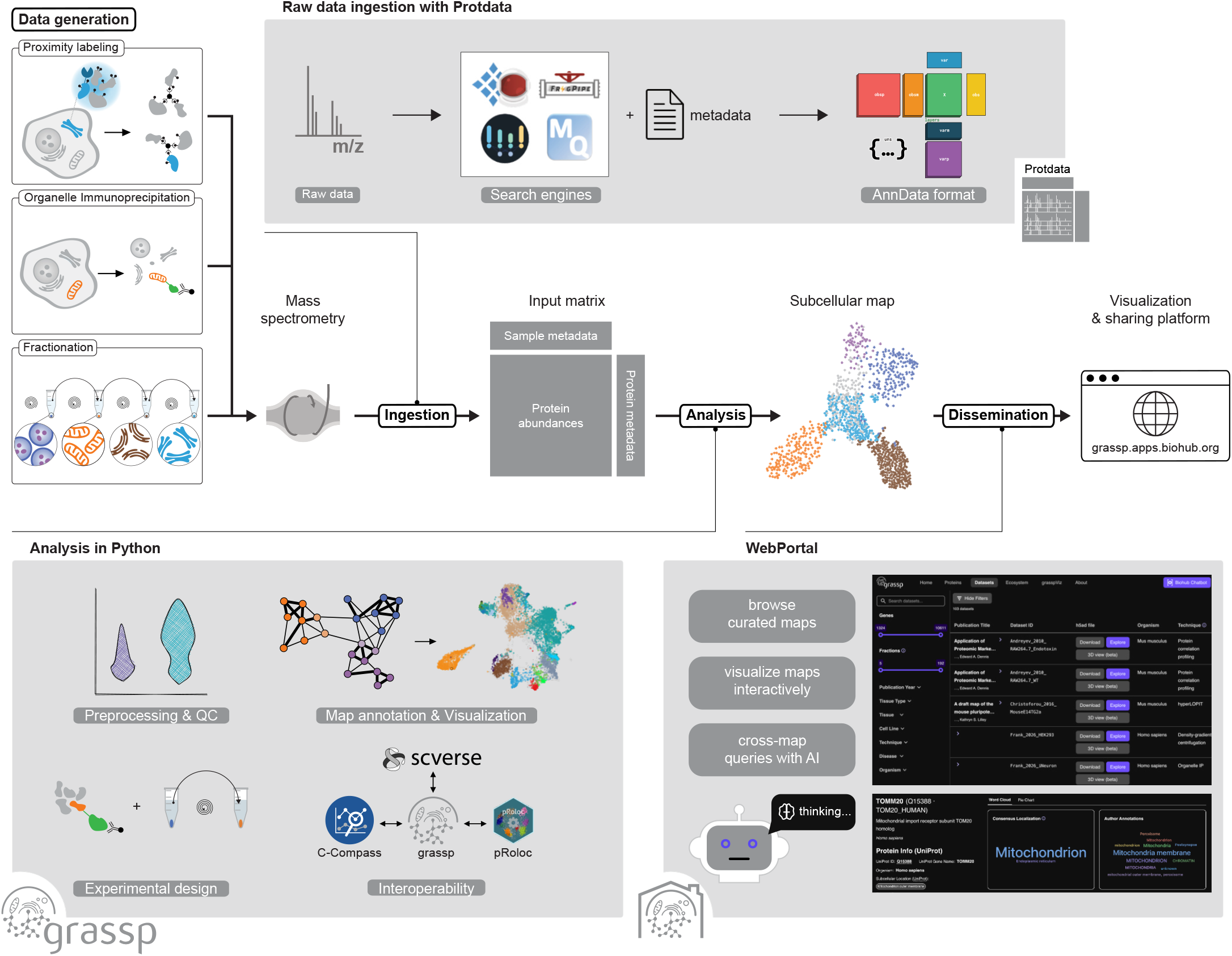
*grassp* provides an end-to-end framework for subcellular proteomics analysis, integration, and exploration. Quantitative mass spectrometry outputs from proximity labeling, organelle immunoprecipitation, and fractionation experiments are imported by *protdata* into AnnData objects that retain protein abundances and experimental metadata. The *grassp* Python package supports preprocessing, quality control, protein-neighborhood graph construction, compartment annotation, visualization, and map integration. The companion *grasspio* R package enables interoperability with the pRoloc analysis framework. Measurements, metadata, graphs, annotations, and analysis results remain together in portable objects that can be shared through the *grassp* portal and inspected interactively in the web-based *grasspViz* viewer. Programmatic access and natural-language queries enable researchers and software agents to discover, explore, and analyze localization evidence across studies.

The *grassp* Python package is the central analytical layer of the framework. It provides functions for preprocessing, quality control, profile transformation, protein-neighborhood graph construction, dimensionality reduction, clustering, and visualization. Its central functionality uses this graph to analyze and annotate subcellular maps (see below). Each analysis step updates the AnnData object to add new information.

Beyond its core workflow, *grassp* connects to established methods across the MS-SP ecosystem. It includes native Python reimplementations of methods available in pRoloc [15]: support vector machines (SVMs) and TAGM [17] for compartment annotation, and `QSep` [20] for assessing map resolution. For access to the broader pRoloc functionality, the companion *grasspio* R package converts processed AnnData objects to and from the `MSnSet` class used in R [30]. *grassp* also wraps C-COMPASS [25], a neural-network method for detecting protein multi-localization and condition-dependent relocalization, and writes its predictions to standard AnnData objects (Methods).

Processed objects can subsequently be published through the *grassp* web portal and directly explored in the *grasspViz* interactive viewer. Because embeddings, annotations, profiles, and statistical results are already stored in the AnnData object, users can inspect and share an analysis without rerunning its computational steps.

By connecting modular tools through a shared AnnData representation and interfaces to other analysis software, the *grassp* framework supports the complete computational workflow for subcellular proteomics, from data ingestion and analysis to interactive exploration and data sharing.

### 2.2 *grassp* annotates subcellular maps through a common graph representation

A fundamental task in subcellular proteomics is to assign proteins to subcellular compartments based on their relative abundance profiles across cellular fractions with differential compartment enrichment. After assay-specific preprocessing (detailed in a collection of tutorials [31]), *grassp* represents each map as a protein-neighborhood graph that links each protein to its nearest neighbors in profile space (Fig. 2a). Edges are weighted by an affinity between 0 and 1 that decreases with profile distance, scaled to the local neighborhood density [32] (Methods). Such graphs are useful in many analytical tasks beyond dimensionality reduction [33]. Because they abstract away the scale and dimensionality of the measurements, they enable consistent analyses across datasets and experimental designs. *grassp* provides two complementary graph-based annotation strategies, competitive diffusion and independent diffusion, both of which assign each protein a probability for each candidate compartment that can be thresholded to prioritize confident assignments.

**Fig. 2.**
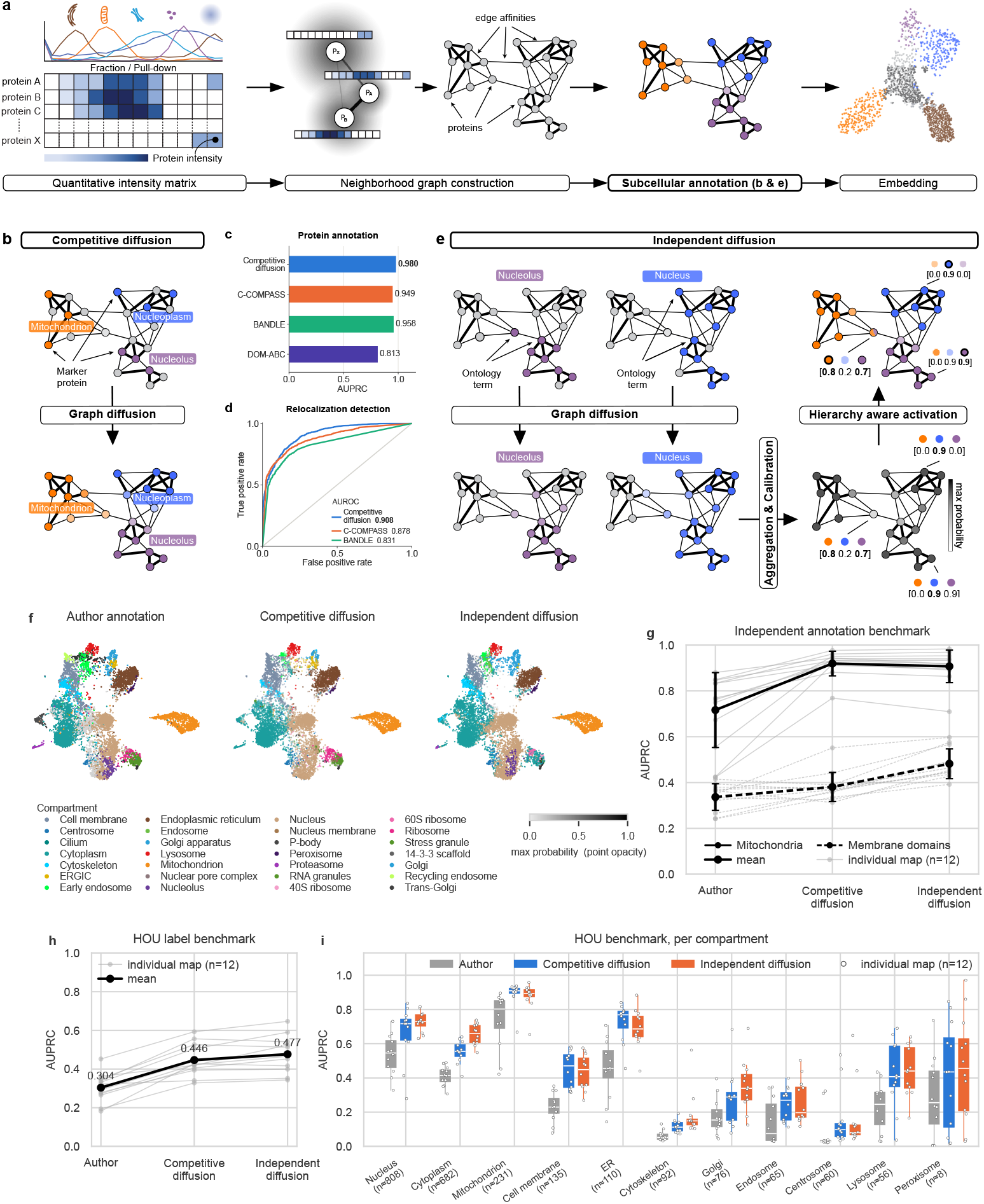
Graph-based annotation methods match or outperform established methods in predicting protein subcellular localization. **(a)** The *grassp* annotation workflow. Relative protein intensities across fractions or pulldowns form a quantitative intensity matrix, from which *grassp* builds a protein-neighborhood graph, with edges weighted by profile affinity, as the substrate for subcellular annotation (**b**, **e**) and embedding calculation. **(b)** Competitive diffusion is an annotation strategy that diffuses mutually exclusive compartment labels from marker proteins across the graph. Compartment probabilities compete in local neighborhoods. **(c)** Comparison of competitive diffusion with established methods that support marker-based compartment prediction, using a simulated dataset from Haas et al. [25]. Performance is measured as area under the precision–recall curve (AUPRC), macro-averaged over eight simulated compartments. **(d)** Comparison of competitive diffusion with existing algorithms for detecting relocalization, on a simulated dataset from Haas et al. Performance is shown with the receiver operating characteristic (ROC) curve. **(e)** Independent diffusion is a gene-set-library-based annotation strategy that tailors compartment specificity to each map’s resolution. Ontology-term memberships are diffused independently (one-vs-rest) across the graph. The resulting per-term scores are aggregated and calibrated for each protein. A hierarchy-aware activation selects each protein’s label from its compartment-probability vector, resolving nested compartments (e.g., nucleolus within nucleus). **(f)** Comparison of three annotation sets on the HEK293T OrgIP map from Hein et al. [3]. The same UMAP embedding is shown in three side-by-side views, with proteins colored according to author annotation, competitive diffusion, or independent diffusion. Competitive diffusion is seeded with the literature-curated marker set of Hein et al., whereas independent diffusion uses the consolidated UniProt subcellular-location vocabulary. Point opacity indicates the maximum label probability. **(g)** Benchmarking competitive diffusion, independent diffusion, and the original author annotations against external references across 12 published subcellular maps. Mitochondrial localization is evaluated using the intersection of MitoCarta and MitoCoP annotations, whereas assignment to membrane-bounded compartments is evaluated using proteins with transmembrane or intramembrane segments annotated in UniProt. Performance is measured by AUPRC. Each gray line represents one of 12 published subcellular maps (Supplementary Table S1). Thick black lines connect the mean values. Error bars indicate *±* s.d. Solid and dashed lines denote the mitochondrial and membrane-domain benchmarks, respectively. **(h)** Held-out evaluation of competitive and independent diffusion across 12 published maps using the HOU (Human Protein Atlas, OpenCell, and UniProt) benchmark [47]. Original author annotations are shown for comparison. Competitive diffusion is seeded with curated compartment markers from Hein et al. [3], whereas independent diffusion uses UniProt-derived compartment gene sets. Both seed sources are restricted to HOU training proteins, with test-protein gene symbols additionally excluded from the seeding gene sets used in independent diffusion approaches. Original author annotations are evaluated as published. Performance is measured by AUPRC on HOU test proteins, macro-averaged over the scored compartments from 11 shared compartment classes. Each gray line represents one of 12 published subcellular maps, and thick black lines connect the mean values. **(i)** Results from the evaluation in **h** are shown separately for each compartment. Each point represents one of 12 published subcellular maps. Boxes show medians and interquartile ranges, whiskers indicate 1.5 times the interquartile range, and *n* denotes the mean number of HOU-positive test proteins per map.

#### 2.2.1 Marker-based annotation with competitive diffusion

When marker proteins with well-established subcellular localizations are available, their labels can be used to annotate the remaining proteins in a map. *Competitive diffusion* is a graph propagation method based on label spreading [34]. Marker labels propagate through the graph according to its edge affinities, and propagation is repeated until the compartment probabilities converge (Fig. 2b and Methods). For each protein, probabilities are normalized to sum to one, so stronger support for one compartment reduces the probability assigned to others. Information flows through unlabeled as well as labeled proteins, making this a semi-supervised approach, which has been shown to improve classification performance [35].

We benchmarked competitive diffusion on a simulated fractionation dataset introduced with C-COMPASS [25], against the three methods compared in that study (C-COMPASS [25], BANDLE [19], and DOM-ABC). Competitive diffusion assigned non-marker proteins to their simulated compartments more accurately than these methods, as measured by the area under the precision–recall curve (AUPRC) averaged over the eight simulated compartments (Fig. 2c and Supplementary Fig. S2). It also better distinguished proteins whose simulated compartment changed between the two conditions (relocalization) from proteins that stayed in place, as measured by the area under the receiver operating characteristic curve (AUROC; Fig. 2d). For proteins simulated in two or three compartments, a multi-localization task for which C-COMPASS was purpose-built, competitive diffusion achieved the second-lowest localization error (Methods), close to that of C-COMPASS (Supplementary Fig. S2d). Competitive diffusion outperformed other methods in terms of runtime (Supplementary Fig. S1) and was run with default parameters for all benchmarks.

#### 2.2.2 Marker-free annotation with independent diffusion

Marker-based annotation can be limiting when curated marker sets are unavailable or incomplete, and the resolution of the final annotation is constrained by the predefined label set. To reduce dependence on bespoke marker panels, *grassp* provides *independent diffusion* using structured ontology vocabularies for protein localization such as Gene Ontology cellular component (GO CC) [36] or UniProt subcellular-localization terms [37] (Fig. 2e). These vocabularies annotate far more proteins than typical curated marker sets and describe compartments at finer resolution. However, their annotations can be noisier, can include cell-type-specific labels, and often assign the same protein to several nested compartments at different resolutions (e.g., nucleolus and nucleus). We therefore developed a novel annotation strategy, building on methods used in gene function prediction from interaction networks [38, 39]. Here, each term is independently diffused through the graph, allowing a protein to accumulate evidence for multiple subcellular compartments without forcing their probabilities to compete. The resulting scores are calibrated into a probability for each protein and each localization term, preserving evidence for overlapping localizations and annotations at different levels of resolution (Supplementary Fig. S3). Finally, a hierarchy-aware resolver uses the nested structure of the vocabulary to select a primary annotation (Fig. 2e, Methods). Alongside this primary annotation, the method reports a multi-localization label based on the independent annotation probabilities.

#### 2.2.3 Benchmarking annotation across diverse subcellular maps

We first assessed agreement with expert annotations using a published OrgIP map of HEK293T cells, which had been annotated through unsupervised high-resolution clustering followed by expert curation [3]. Competitive and independent diffusion matched the published annotations for 80% and 82% of labeled proteins, respectively (Fig. 2f). Differences in input vocabularies contributed to the remaining disagreements. For example, the nucleolus and the proteasome, which were labeled in the original study, are not among the marker classes that seed competitive diffusion.

Next, we evaluated whether the new algorithms are robust across a diverse set of subcellular maps. To this end, we curated and reprocessed 12 published datasets (Supplementary Table S1) that span a variety of enrichment techniques and MS methods. The enrichment techniques comprise OrgIP [3], density-gradient fractionation (hyperLOPIT [10, 40] and dLOPIT [41]), DC-based profiling (Dynamic Organellar Maps [11–13], LOPIT-DC [42, 43], and DIA-LOP [44]), proximity labeling (BioID) [14], and detergent-based fractionation (SubCellBarCode) [23]. The MS methods include data-dependent and data-independent acquisition, with label-free or label-based quantification. For each map, we evaluated competitive diffusion, independent diffusion, and the original author annotations using AUPRC, which summarizes performance of retrieving ground-truth labels across score thresholds without requiring a single confidence cutoff. This metric is useful for compartment prediction, where positive and negative classes are often strongly imbalanced.

We also evaluated the strategies against two external references based on strong biological priors: mitochondrial proteins listed in both MitoCarta3.0 and MitoCoP [45, 46], and proteins with UniProt-annotated transmembrane or intramembrane segments, which should reside in membrane-bounded compartments. Both diffusion strategies recovered mitochondrial proteins better than the original study annotations in all 12 maps (AUPRC, mean ± s.d. 0.92 ± 0.05 and 0.91 ± 0.07, versus 0.72 ± 0.16; Fig. 2g). For membrane-bounded compartments, competitive diffusion improved on the original annotations in 7 of the 12 maps (0.38 ± 0.06 versus 0.34 ± 0.06) and independent diffusion in all 12 (0.48 ± 0.06).

We next evaluated prediction of held-out annotations using the recently published HOU benchmark [47], which combines localization evidence from the **H**uman Protein Atlas [48], **O**penCell [49], and **U**niProt [37]. Performance was assessed both across compartments, and separately for each compartment (Fig. 2h,i). Here again, both competitive and independent diffusion strategies showed improved annotation performance relative to the original study annotations.

In all cases, methods were applied with default settings and without dataset-specific manual parameter tuning, and ran faster than comparable methods (Supplementary Fig. S1). Together, these results support the applicability of graph-based annotation across a wide repertoire of MS-SP experimental designs (Supplementary Table S1).

### 2.3 *grassp* enables comparison and integration of subcellular maps

#### 2.3.1 *grassp* recovers documented neuronal localizations despite weaker organelle enrichment

To demonstrate the applicability of *grassp* to a new biological system, we generated a subcellular map of day-21 NGN2-induced neurons (iNeurons) derived from human induced pluripotent stem cells (iPSCs). The dataset profiles 5973 proteins across 14 OrgIP baits and three centrifugation fractions (nuclear, organellar, and cytosolic) (Supplementary Table S2). Because iNeurons are costly to culture and grow at low density, each OrgIP sample used 7.5 × 10^5^ cells (one well of a 6-well plate), about 13-fold fewer than the 10^7^ HEK293T cells per sample in our previous OrgIP study [3]. Consistent with this lower input, organelle markers were less enriched in iNeurons than in HEK293T cells (median log_2_ enrichment of 3.0 versus 6.2 for mitochondrial, 0.6 versus 2.9 for plasma membrane, and 0.7 versus 2.1 for endoplasmic reticulum (ER) markers, respectively). The iNeuron dataset therefore tested *grassp* under experimentally challenging conditions. Despite this weaker enrichment, held-out organelle markers were annotated comparably well in both HEK293T and iNeuron maps (0.89 vs 0.91 AUPRC respectively for 9 compartments with shared baits) using the competitive diffusion strategy (Fig. 3a,b and Supplementary Fig. S4b,c).

**Fig. 3.**
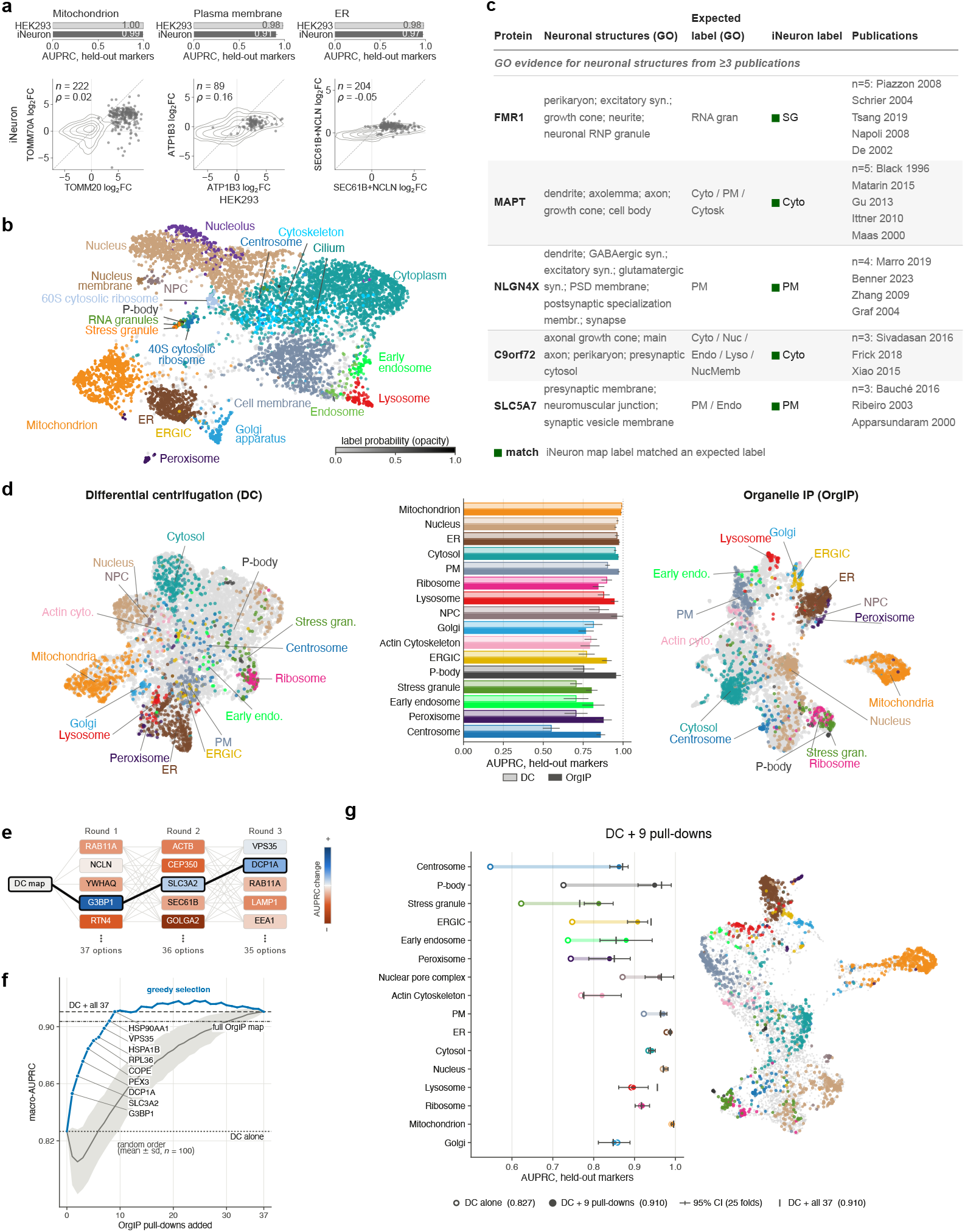
*grassp* enables comparison across cellular systems and integration of complementary subcellular maps. **(a)** The iNeuron map processed with *grassp* shows only slightly lower annotation performance than the HEK293T OrgIP map, despite weaker marker enrichment in the iNeuron pulldowns. Bars depict resolution of three example compartments, measured by the area under the precision–recall curve (AUPRC) of held-out marker protein retrieval (25 cross-validation folds: 5fold, 5 repeats; whiskers represent 95% confidence intervals over folds). Scatter plots depict log_2_ fold change of proteins in pulldown experiments of characteristic bait proteins for a cellular compartment (Mitochondrion: TOMM70A in iNeurons and TOMM20 in HEK293T cells. Plasma membrane: ATP1B3. Endoplasmic reticulum: SEC61B and NCLN). Fold changes of multiple pulldowns were averaged. Gray points depict marker proteins curated by Hein et al. [3], and gray contours are the kernel density of all other shared proteins under the same contrasts. Here, *n* indicates the number of shared marker proteins for each compartment. The remaining compartments are shown in Supplementary Fig. S4. *ρ*, Spearman correlation between the two maps across markers. **(b)** UMAP representation of the iNeuron subcellular-proteomics map (5973 proteins), colored by compartment label assigned to each protein by *grassp* independent diffusion using UniProt annotations. Labels identify compartment neighborhoods. Opacity indicates label probability. **(c)** Comparison of compartment assignments from *grassp* in the iNeuron map with expected compartment labels for the five reference proteins with three or more supporting publications (the remaining 12 reference proteins with two publications are shown in Supplementary Table S4). Abbreviations: GO, Gene Ontology; Cyto, cytoplasm; Nuc, nucleus; PM, plasma membrane; Cytosk, cytoskeleton; Endo, endosome; Lyso, lysosome; NucMemb, nuclear membrane; RNA gran, RNA granules; SG, stress granule; RNP, ribonucleoprotein; PSD, postsynaptic density; syn., synapse; membr., membrane; neurite, neuron projection; cell body, neuronal cell body. **(d)** Compartment resolution of two HEK293T maps built with different enrichment techniques: differential centrifugation (DC; this study) and organelle immunoprecipitation (OrgIP; Hein et al. [3]). Left and right, UMAPs of the DC and OrgIP maps, respectively, colored by marker compartment. Center, per-compartment AUPRC for competitive diffusion predictions on held-out compartment markers; light bars, DC; dark bars, OrgIP. Error bars, 95% confidence intervals over 25 cross-validation folds (5-fold, 5 repeats). **(e)** Schematic of the forward greedy selection of OrgIP pulldowns that best complement the DC map, showing the first three of 37 rounds (all rounds in **f**). Starting from the DC map, each round tests every remaining pulldown (boxes, named by bait) as an addition to the current map and keeps the one giving the highest macro-averaged AUPRC on held-out markers. Colors indicate the change in AUPRC. Black outlines and connecting lines mark the selected path. The total number of candidates evaluated in each round is given below. **(f)** Resolution of the combined map as OrgIP pulldowns are added to the DC map in greedy order (blue) or random order (gray line and band, mean *±* s.d. over 100 orders). Resolution is the macro-averaged AUPRC on held-out markers (mean of 25 cross-validation folds; 5-fold, 5 repeats). Labels, first nine greedily selected pulldowns (by bait). Dotted line, DC map alone; dash-dotted line, full OrgIP map alone; dashed line, DC map plus all 37 pulldowns. Nine greedily selected pulldowns reach the resolution of all 37. The early dip under random order and the late decline under greedy selection show that some pulldowns lower resolution. **(g)** Compartment resolution of the DC map before and after adding the nine pulldowns labeled in **f**. Left, per-compartment AUPRC on held-out markers for the DC map alone (open circles), with the nine pulldowns (filled circles), and with all 37 pulldowns (black ticks). Error bars (drawn for filled circles) are 95% confidence intervals over 25 cross-validation folds (5-fold, 5 repeats). Right, UMAP representation of the DC map with the nine pulldowns, colored by marker compartment as on the left. In **e**–**g**, the DC and OrgIP maps were restricted to 7667 shared genes after collapsing protein accessions by gene symbol.

Next, we examined whether individual protein-localization results agreed with published evidence for neuron-specific localization. Using stringent filters for experimental support (Methods), we compiled a GO-derived reference set of 17 human proteins with neuron-specific localizations that were detected in the iNeuron dataset (Methods; Supplementary Table S3). Of these, 13 received compartment assignments that matched the literature, either through exact label matches or coarse matches at the level of a broader cellular compartment (Fig. 3c; Supplementary Table S4), while the four nonmatches had plausible biological explanations. Of these, RPLP0 (a known ribosomal protein [50]) is annotated in the literature as localized to the postsynaptic density while *grassp* assigns it to the 60S ribosome. GPHN’s cytoplasmic assignment and AKAP5’s plasma membrane assignment are compatible with their respective roles in cytoskeletal scaffolding [51] and postsynaptic membrane anchoring [52]. SNAPIN’s lysosomal assignment is consistent with its role as a subunit of the BORC complex, which associates with the lysosomal membrane and regulates lysosome positioning [53]. Together, these results show that the *grassp*-annotated iNeuron map recovers documented neuronal localizations despite weaker organelle enrichment.

#### 2.3.2 Integration of subcellular maps across experimental systems with *grassp*

Because *grassp* is agnostic to the enrichment technique, maps from different assays can be combined. We sought to demonstrate how combining experimental strategies could reduce the experimental effort needed to build high-resolution maps. Specifically, OrgIP and DC offer complementary tradeoffs. OrgIP isolates compartments in a targeted manner and therefore can achieve higher resolution than DC [3], but every bait requires engineering a CRISPR-tagged cell line and a new immunoprecipitation experiment. DC needs no cell-line engineering and covers the proteome broadly, but separates compartments only by sedimentation rate. We therefore asked whether a DC map supplemented with a small, targeted set of OrgIP pulldowns could match the resolution of a full OrgIP map.

We generated a DC dataset of HEK293T cells covering 8595 proteins across seven fractions (Supplementary Table S5). DC resolved large compartments, such as the mitochondrion, the ER, and the nucleus, nearly as well as a recently published OrgIP map of the same cell line [3]. The OrgIP map better resolved smaller compartments and assemblies, including P-bodies, peroxisomes, centrosomes, early endosomes, the nuclear pore complex (NPC), and the ER–Golgi intermediate compartment (ERGIC) (Fig. 3d).

To find the pulldowns that best complement DC, we used a forward greedy selection strategy. Starting from the DC map, each step added the pulldown that maximized the resolution of the combined map, measured as AUPRC on held-out markers (Fig. 3e). Nine greedily selected pulldowns raised the AUPRC of the DC map from 0.83 to 0.91, matching the DC map combined with all 37 pulldowns and exceeding the full OrgIP map (0.90, Fig. 3f). Their gains were concentrated in compartments that DC alone resolved poorly (Fig. 3g). Pulldowns added in random order improved resolution far more slowly, and the first few even lowered it (Fig. 3f, gray curve). Stepwise addition to a DC map can therefore guide the choice of a minimal OrgIP panel for a given experimental budget.

#### 2.3.3 An integrated pan-human map resolves subcellular organization across scales

We next evaluated whether *grassp*’s common graph representation could support integration and harmonization at a broader scale: across studies, biological systems, and experimental platforms whose raw measurement dimensions cannot be aligned directly. We uniformly processed 101 human, mouse, and yeast MS-SP datasets published since 2010 and spanning the whole spectrum of MS-SP assays (Fig. 4a and Supplementary Table S6), together with the new iNeuron OrgIP and HEK293T DC datasets from this study. Starting from quantitative protein-abundance matrices from each dataset, *grassp* constructed a protein-neighborhood graph and assigned compartments using the robust label diffusion strategies introduced above. This resulted in harmonized annotations that can be compared across datasets and that were more consistent between maps than the authors’ published labels (Supplementary Fig. S5). We then used BIONIC, a graph convolutional network (GCN)-based integration method [54], to combine the protein-neighborhood graphs from 51 unperturbed human maps (maps of experimentally perturbed samples were excluded) into an integrated pan-human map (Fig. 4b,d). The resulting graph captures protein–protein relationships supported across the collection. Importantly, this integration is unsupervised: no protein annotations were used for training.

**Fig. 4.**
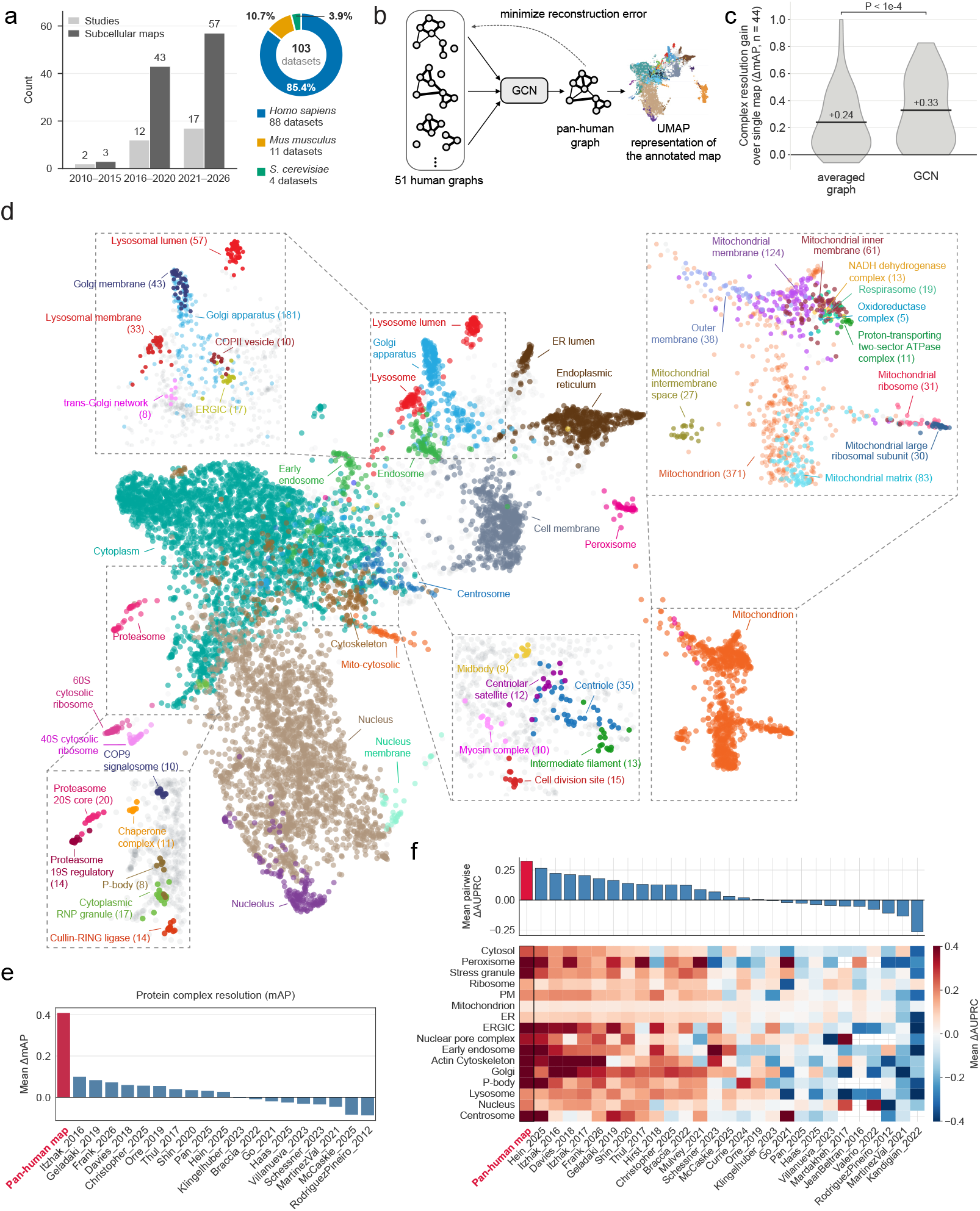
Integrating subcellular maps across studies produces a pan-human map with improved compartment and protein-complex resolution. **(a)** Left, number of studies (light gray) and individual subcellular maps (dark gray) processed in this study, grouped by publication period. Right, species composition of the complete collection, comprising 101 published maps and two generated in this study: *Homo sapiens* (*n* = 88), *Mus musculus* (*n* = 11), and *Saccharomyces cerevisiae* (*n* = 4). **(b)** Integration strategy. Rather than merging heterogeneous raw data, the protein-neighborhood graphs of the 51 individual unperturbed human subcellular maps are integrated with a graph convolutional network (GCN) trained to minimize graph reconstruction error, yielding a single pan-human graph and a UMAP representation of the annotated map. **(c)** GCN integration improves protein-complex resolution beyond graph averaging. The 44 benchmark protein complexes were derived from human EBI Complex Portal entries requiring manually curated physical-interaction evidence and containing at least six listed members. After deduplication and assignment of each protein to a single complex, groups with at least two assigned members present in the integrated pan-human map were retained. Violins show the distribution of per-complex improvement in mean average precision (mAP) over the high-resolution HEK293T map (Hein et al. [3]). The black bar is the mean over complexes (averaged graph +0.24, GCN +0.33). The *P* value was calculated using a two-sided Wilcoxon signed-rank test of GCN against the averaged graph, paired by complex (*P* = 1.2 *×* 10*^−^*^5^). See also Supplementary Fig. S7. **(d)** UMAP visualization of the integrated human map (9530 proteins), colored by *grassp* independent diffusion annotation. The global view uses the consolidated (coarse) UniProt subcellular-location vocabulary. Insets zoom into fine substructure resolved by the map, displaying selected independent diffusion annotations using the Gene Ontology Cellular Component (GO CC) and COMPARTMENTS vocabularies. Proteins are colored only when their assigned term has a probability of at least 0.6. Numbers in parentheses give the number of proteins colored per fine compartment. “Mito-cytosolic” marks a 110-protein cluster described in the text (Supplementary Fig. S8b), not a vocabulary term. **(e)** Protein-complex resolution assessed by pairwise comparisons across 59 maps with sufficient complex coverage. Resolution is measured by mean average precision (mAP), which quantifies how highly members of the same complex rank. Scores are averaged across reference complexes from the EBI Complex Portal [66] (Methods). Each comparison is restricted to proteins shared by the two maps and complexes with at least four members present. Bars show each map’s mean pairwise mAP difference, ordered from highest to lowest. Positive values indicate better resolution than its comparison maps on average. The integrated map (crimson) ranks first, with a mean difference of +0.41. The plot displays 20 of the 59 maps: the integrated map and, for 19 publications, the representative map chosen as in **f** (with one additional constraint: publications whose representative lacked sufficient complex coverage are not shown). **(f)** Compartment resolution assessed by pairwise comparisons across 89 maps, using the area under the precision–recall curve (AUPRC) of competitive diffusion predictions for held-out ground-truth markers from Hein et al. [3] (5-fold cross-validation with identical folds for both maps of a pair). Each pairwise comparison was restricted to shared proteins and compartments scored in both maps (see Methods and Supplementary Fig. S6a). Top, mean pairwise AUPRC differences for each map, averaged across compartments and sorted in descending order. The integrated map (crimson) ranks first. Positive values indicate higher AUPRC for the indicated map, on average, than for its comparison maps. Bottom, the corresponding mean differences for individual compartments (red, higher AUPRC for the indicated map; blue, lower; white cells with crosshairs, compartment not scored for that map). The display includes the integrated map and one representative map from each of the 27 publications, the map with the highest single-map marker macro-F1 (dataset names are in Supplementary Table S13).

Altogether, the integrated map included 9530 proteins, of which 8673 (91%) were assigned to a subcellular compartment by independent diffusion under the consolidated UniProt vocabulary. We benchmarked the ability of the integrated map to predict the labels of held-out compartment marker proteins against individual datasets. The integrated map outperformed every individual map based on mean pairwise performance differences, with resolution gains across nearly every subcellular compartment (Fig. 4f and Supplementary Fig. S6a,b; see Methods for details). For a more challenging task, we used the mean average precision (mAP) metric [55] to quantify how well protein complexes could be resolved in the integrated map. This benchmark showed an even greater increase in resolution compared to individual maps (Fig. 4e and Supplementary Fig. S6c), indicating that integrating across datasets has the potential to resolve subcompartment structures down to the level of protein complexes. To separate the contribution of pooling many maps from that of the integration method, we compared GCN with two simpler integration strategies: the annotation mean, which averages the per-map compartment probabilities without building a shared graph, and a graph with averaged edge weights of the 51 input graphs. At the compartment level, all three strategies reached the same held-out marker resolution and outperformed the best single map (Supplementary Fig. S7a), indicating that this gain comes from breadth of input rather than from the integration strategy itself. At the protein-complex level, however, GCN integration significantly outperformed the averaged graph baseline (Fig. 4c and Supplementary Fig. S7b; note that the annotation mean strategy is not applicable in this case).

These performance gains were accompanied by biologically coherent organization at multiple scales. The integrated map resolved fine structure in several regions, including those associated with cytoskeletal and cell-division assemblies, cytoplasmic ribonucleoprotein (RNP) and proteostasis machinery, and the Golgi–endolysosomal system (Fig. 4d, insets). The mitochondrial region provides a detailed example of suborganellar resolution: the map disambiguated matrix, intermembrane-space, inner- and outer-membrane, and mitoribosomal neighborhoods (Fig. 4d, top right inset and Supplementary Table S7). Among proteins receiving fine submitochondrial labels, these assignments agreed closely with MitoCarta/MitoCoP subcompartment reference annotations (92.1% accuracy; 0.89 macro-F1). Importantly, the independent diffusion algorithm allowed annotations to be finer or coarser depending on the resolution of the local neighborhood, falling back to the generic “Mitochondrion” term when there was insufficient enrichment for a subcompartment label (Supplementary Fig. S8a). UniProt transmembrane annotations provided orthogonal validation: annotated transmembrane segments occurred in 155 of 223 proteins (69.5%) assigned to membrane regions, compared with 1 of 171 (0.6%) assigned to the matrix, intermembrane-space, and mitoribosomal regions (two-sided Fisher’s exact test, *P* = 1.4×10^−53^; Fig. 4d, top right inset). Oxidative-phosphorylation assemblies, including ATP synthase, complex I, and the I-III_2_-IV respirasome, also occupied a compact inner-membrane neighborhood (Fig. 4d, top right inset).

The integrated map also resolved a distinct 110-protein cluster between the mitochondrial island and the cytosol (“Mito-cytosolic” in Fig. 4d, Supplementary Fig. S8b, Supplementary Table S8). Compared with the mitochondrial island (849 proteins; Supplementary Fig. S8b–e), this cluster was enriched for proteins annotated as mitochondrial-matrix in MitoCarta3.0 [45] (80% versus 48% of MitoCarta3.0 members in each group; Supplementary Fig. S8c), depleted of proteins with transmembrane segments (3.6% versus 30% of all proteins in each group), and more frequently annotated by UniProt as having dual localization to both mitochondria and the cytoplasm or cytosol (33% versus 7.9%; Supplementary Fig. S8d). HEK293T OrgIP data showed that proteins in the Mito-cytosolic cluster were enriched in a mitochondrial pulldown (median log_2_ enrichment 4.9, versus 6.6 in the mitochondrial island) but also abundant in the cytosolic centrifugation fraction (median 65% of each protein), while proteins in the mitochondrial island were not (below 1%; Supplementary Fig. S8e). Several Mito-cytosolic cluster members have compartment-specific functions that provide a biological basis for dual localization. GARS1 charges tRNAs for both mitochondrial and cytosolic translation [56], while FPGS maintains separate folate pools supporting mitochondrial one-carbon metabolism and cytosolic nucleotide synthesis, because polyglutamylated folates cannot cross the mitochondrial membrane [57]. ARL2 performs distinct regulatory functions in the two compartments, contributing to cytosolic tubulin homeostasis [58, 59] and regulating mitochondrial fusion [60]. Together, these findings support the cluster as a candidate population with both mitochondrial and cytosolic localization evidence.

We next asked whether the integrated map provides localization information that existing databases do not. We first compared it with SurfaceBench, a curated benchmark of cell-surface accessibility enriched for proteins on which databases disagree [61]. The integrated map included 21 proteins annotated in SurfaceBench as “permanently accessible” from the extracellular surface and 20 “conditionally accessible” proteins. With all benchmark proteins held out of the annotation vocabulary, the plasma membrane was the highest-scoring compartment for 18 of the 21 “permanently accessible” proteins in the integrated map (86%, Supplementary Fig. S9), while the map’s annotation for the remaining three proteins matched their known biological function: TFRC, the transferrin receptor, is a known recycling protein constantly shuttled between the cell membrane and endocytic compartments [62]; GPNMB has recently been identified as a lysosomal protein [63]; and MUC16 is a mucin, the core component of the secreted extracellular mucus [64]. Additionally, all 19 “conditionally accessible” proteins with a stated baseline location were placed in the compartment that the benchmark curators describe as that location (Supplementary Fig. S9 and Supplementary Table S9). This agreement where databases diverge prompted us to examine proteins for which databases offer no localization at all. The integrated map generated localization hypotheses for 400 human proteins lacking both a UniProt subcellular-location annotation and a high-confidence COMPARTMENTS assignment (Supplementary Table S10). Using teams of curator and verifier AI agents, we reviewed the primary literature for all 400 proteins and focused the comparison on the 131 with a maximum integrated-map annotation probability *>* 0.8 (Methods and Supplementary Table S11). Published studies in human cells or tissues reported localizations for 61 of these 131 proteins, 54 of which could be compared with the map assignment. The map assignment agreed with the published observations for 50 of these 54 proteins (92.6%), 40 at the level of a specific compartment and 10 at the broader level of the cytoplasm. The remaining 70 had no qualifying localization evidence in human cells or tissues. Although agreement with the literature is not direct experimental validation (which is out of scope for the current study), the high agreement rate we observed above lends confidence to the map assignments for these 70 proteins, which are candidates for experimental follow-up.

Finally, we tested if the integrated map encodes functional information. We used PRESAGE, a model that predicts transcriptional responses to unseen genetic perturbations using gene embeddings from different biological knowledge sources [65] (Methods). We summarized performance across four prediction tasks in a composite score, scaled so that 0 corresponds to chance and 1 to the default PRESAGE model, which combines 40 knowledge sources (see Methods). As the only knowledge source, the integrated map reached a composite score of 0.72, ranking fifth of 41 sources (Supplementary Fig. S10a and Supplementary Table S12). It was surpassed only by three variants of the STRING protein–protein interaction network and by DepMap gene-dependency profiles (0.74–0.78). Notably, it was the highest-ranked source of any other kind, ahead of curated Gene Ontology cellular-component annotations, pathway geneset collections, and protein language-model and text-derived embeddings. None of the 51 individual input maps came close to the integrated map when used in the same way: the best scored 0.55 (Supplementary Fig. S10c). Importantly, the integrated map contributes information not captured by the 40 existing knowledge sources: adding it as an additional source still raised the composite score from 1.00 to 1.02 (*P* = 0.016, two-sided Wilcoxon signed-rank test), whereas a control map with shuffled gene labels left it unchanged (1.00, *P* = 0.57; Supplementary Fig. S10b).

Together, these different analyses demonstrate two scales of integration with *grassp*: combining assays within a cellular system to improve resolution efficiently, and combining graphs across studies to provide a new reference map of human protein localization that also serves as an informative and complementary prior for models of cellular perturbation responses.

### 2.4 Interactive visualization and comparative analysis through the *grassp* portal

To make MS-SP data a shared and cumulative resource, we developed the *grassp* portal, which hosts the harmonized collection of 103 human, mouse, and yeast maps described above. Each map is available as a standardized AnnData object that contains the underlying measurements as well as annotations, metadata, and analysis results. By combining this collection with tools for interactive exploration and crossdataset analysis, the portal provides a common foundation for investigating protein localization across studies.

#### 2.4.1 Dataset discovery links directly to interactive map exploration

The dataset catalog can be searched by keyword or filtered by protein coverage, number of quantitative dimensions (e.g., fractions or baits), publication year, tissue, cell line, experimental technique, disease, and organism (Fig. 5a), and includes publication and dataset identifiers together with key experimental metadata. Each map can be explored interactively in the *grasspViz* viewer, and each complete *grassp* analysis can be distributed as an h5ad file storing the quantitative matrix, protein metadata, protein-neighborhood graph, embeddings, compartment assignments, and analysis parameters. Users can also load their own .h5ad files, including ones generated or modified with other tools, into *grasspViz*. *grasspViz* presents the stored results without requiring any processing: users can select among stored embeddings, identifiers, annotation labels, and per-protein statistics; search for individual proteins; and select protein groups with lasso or box-selection tools for download or direct Gene Ontology enrichment analysis (Fig. 5b). Each map can be viewed in PCA and UMAP embedding plots. The underlying graphs can be explicitly visualized, while additional visualizations such as heatmaps and volcano plots support deeper exploration. Analyses can be shared through stateful links: URLs that retain the current exploration context, including the selected map and embedding views, coloring, and highlighted proteins. When a recipient opens the link, *grasspViz* loads the corresponding map and restores the state, facilitating collaborative exploration and discussion of specific findings.

**Fig. 5.**
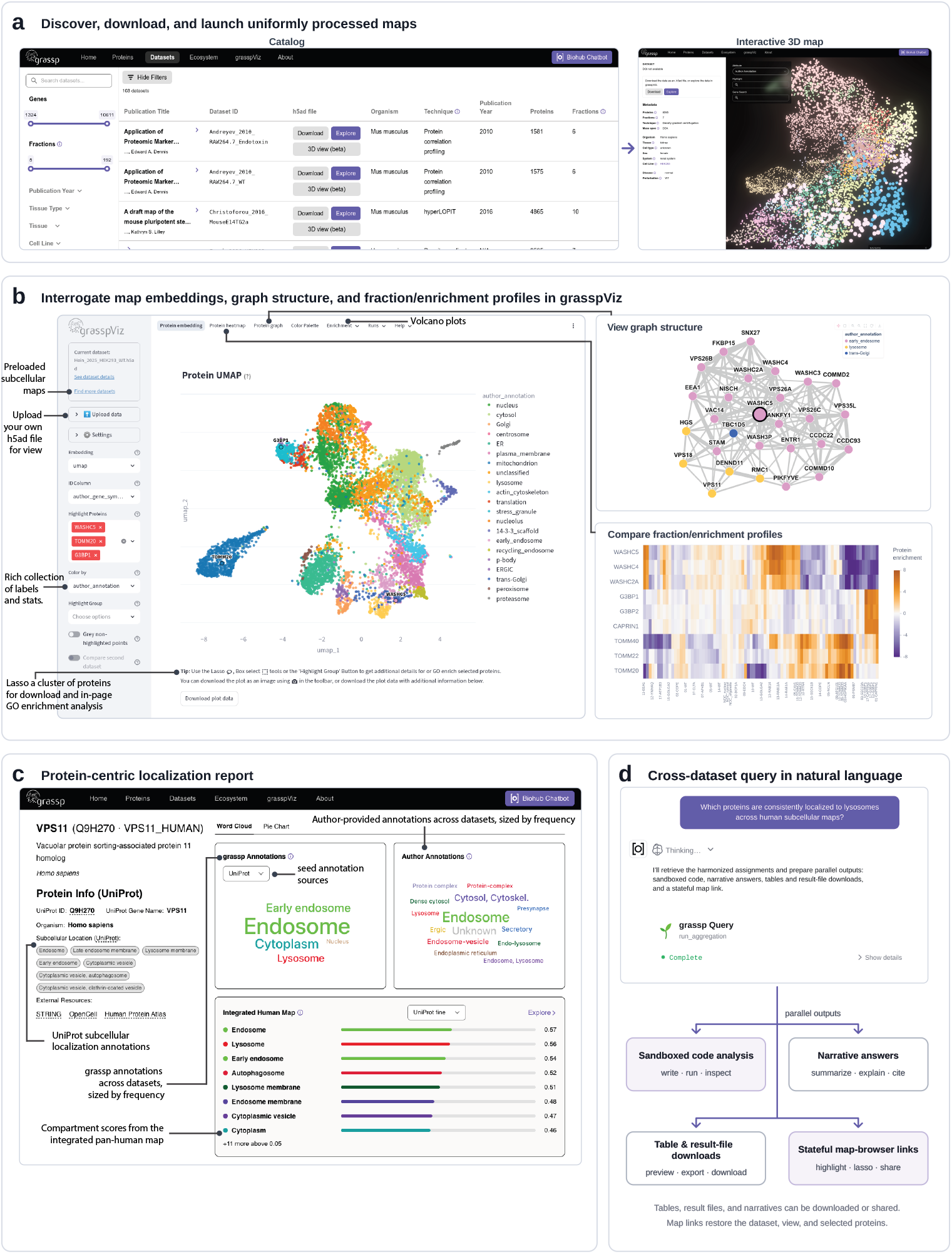
The *grassp* portal supports exploration, comparison, and cross-dataset analysis of subcellular localization maps. **(a)** The dataset catalog supports searching and filtering by biological and technical metadata. Each map can be downloaded as an .h5ad file, launched in *grasspViz*, or previewed in an interactive embedding viewer. The HEK293T differential-centrifugation map (from this study) is shown on the right. **(b)** Interactive exploration of the Hein et al. [3] HEK293T organelle-immunoprecipitation (OrgIP) map in *grasspViz*. The viewer loads maps from the portal or user-uploaded .h5ad files and displays their stored labels and statistics. Besides the protein UMAP, *grasspViz* also displays protein heatmaps, graph structure, and enrichment results such as volcano plots. Upper right, the one-hop neighborhood of WASHC5 in the protein-neighborhood graph. Lower right, fraction/enrichment profiles across the OrgIP baits compare the WASH complex (WASHC2A, WASHC4, WASHC5), the mitochondrial TOM complex (TOMM20, TOMM22, TOMM40), and a G3BP stress-granule module (G3BP1, G3BP2, CAPRIN1). Heatmap rows represent proteins and columns represent pulldown contrasts. Colors indicate protein enrichment. Proteins within each group show similar profiles, whereas the three groups have distinct enrichment patterns. **(c)** Protein-centric view for VPS11. Curated UniProt subcellular-location terms are shown (left) alongside two cross-dataset summaries (right): vocabulary-harmonized *grassp* annotations under the UniProt vocabulary at a selectable coarse or fine resolution (coarse shown) and the original author-provided annotations. Word size represents annotation frequency across datasets. The “Integrated Pan-Human Map” view displays the protein’s compartment scores in that map, allowing the combined evidence to be inspected alongside the distribution of localization assignments across individual studies. Links (left) provide access to external protein resources. **(d)** Cross-dataset natural-language query example asking which proteins are consistently localized to lysosomes across human subcellular maps. Depending on the request, the AI chatbot can produce parallel outputs: analysis with accompanying code executed in a sandbox, narrative answers, downloadable result files, and stateful map-browser links that reopen a prepared map view.

#### 2.4.2 Protein-centric reports and natural-language queries enable cross-dataset analysis

The harmonized data collection also supports protein-centric exploration: users can query a protein (gene name, UniProt accession, or protein name) and compare its localization evidence across all maps (Fig. 5c). Word clouds present the annotations obtained by *grassp*’s harmonized re-analysis of individual maps, together with the original authors’ annotations. Annotation scores from the integrated pan-human map are also presented, with interactive links to the map for exploration. Keeping curated knowledge, standardized calls, and original authors’ labels separate makes agreement and disagreement among these layers visible without collapsing them into a single label.

Finally, the portal is equipped with an AI chatbot that enables users to query and analyze the *grassp* collection in natural language (Fig. 5d). It can provide narrative answers, generate and execute analysis code in a sandbox, return tables and downloadable results, and create stateful links to relevant map views. Overall, the *grassp* portal provides a single point of access for finding datasets, exploring their underlying measurements, and comparing protein localization across the collection.

## 3 Discussion

As subcellular proteomics grows in scale and experimental diversity, building cumulative knowledge and deriving generalizable insights depend on standardizing, comparing, and integrating measurements without erasing the contexts in which they were generated. Here, we introduced the *grassp* Python framework, which converts assay-specific raw datasets into protein-neighborhood graphs and stores measurements, metadata, and analysis results together in AnnData objects. This combination of a portable data object and a common graph representation allows the same downstream methods to operate across experimental designs while retaining access to raw quantitative data and metadata.

Using this common representation, *grassp* provides two complementary strategies for annotating subcellular maps. Competitive diffusion is a computationally efficient marker-based classifier that required no dataset-specific tuning. It is particularly useful when high-confidence markers are available for specific compartments and the biological question can be addressed through protein assignment to those compartments. Independent diffusion uses ontology terms from more comprehensive hierarchical compartment vocabularies. By propagating ontology terms independently, it allows for overlapping and nested labels and adapts annotation specificity to the resolution supported by each map. This makes it useful when the resolution is not known in advance, or when compartments lack curated marker sets but are included in the more comprehensive ontology vocabularies. Across diverse datasets and assay types, independent diffusion matched or outperformed competitive diffusion when evaluated against source-study labels, external compartment markers, and an independent imaging-based reference. Neither method, however, explicitly models replicate-level measurement uncertainty. Incorporating this uncertainty in future variations could improve score calibration and annotation robustness when between-replicate variability is substantial.

Beyond analyzing individual maps, *grassp* enables integration across experimental approaches with orthogonal measurement spaces. DC datasets, for example, typically comprise ∼10 relative abundance measurements with dense coverage, whereas BioID collections may include hundreds of sparse bait enrichment measurements. Representing each dataset as a protein-neighborhood graph shifts the integration problem from incompatible assay-specific axes to a common description of local protein relationships, avoiding direct feature alignment and the need to impute a single cross-study feature matrix. Using this strategy, we combined 51 unperturbed human maps into an integrated pan-human map that improved the resolution of subcellular compartments and protein complexes beyond that of any individual previous study. The integrated map, covering a total of 9530 proteins, also recapitulated fine submitochondrial organization and generated localization hypotheses for 400 proteins lacking both UniProt and COMPARTMENTS subcellular assignments. Together with our comparison and integration of DC and OrgIP maps, these results suggest that different experimental strategies capture complementary aspects of subcellular organization, and that combining their evidence can improve resolution beyond individual maps. Because it outperformed every individual map we evaluated on compartment and proteincomplex resolution, the integrated map provides a new, data-driven reference for human protein localization. The integrated map was also among the most informative knowledge sources when used as a prior to predict genetic perturbation responses, and improved predictions beyond 40 established sources. This shows that the integrated map contained information not covered by established knowledge sources. A current limitation of the integrated map, however, is its heterogeneous source data. The map is shaped by the unequal representation of cell lines, tissues, conditions, and experimental platforms in the source literature, and it cannot distinguish multi-localization within individual cells from context-dependent variation across cell types or datasets. Annotations from the integrated map should therefore be interpreted alongside the distribution of annotations across individual studies, both of which can be explored in the *grassp* portal. We expect the coverage and accuracy of the integrated map to increase as more maps become available.

We envision *grassp* as the foundation for a cumulative, community-driven resource of subcellular proteomics. The standardized AnnData format, shared ontology, and structured metadata are designed so that researchers can deposit their own maps into the *grassp* web portal, where they become immediately discoverable, explorable through *grasspViz*, and available for integration alongside existing data. The *grassp* collection of subcellular maps is built to be read by AI agents as well as people. Similar infrastructure [27] and distribution [67] efforts in the single-cell sequencing field have recently enabled training of large-scale deep-learning models of transcriptomes across cell types, perturbations, and species spanning the evolutionary tree [68–71]. Enabling similar advances in subcellular proteomics requires interoperability across data formats, programming languages, and analysis tools. We therefore built *grassp* to integrate with scverse tools and provided compatibility with R/Bioconductor through *grasspio*.

To date, subcellular proteomics has contributed little to large-scale computational models of the cell, despite the central role of protein localization in cellular function. To fill this gap, grassp paves the way to uniform processing of subcellular maps across a diversity of cell types, conditions, and contexts. Its framework could be further extended to lipidomics and metabolomics to achieve a comprehensive description of subcellular organization. However, each individual map only represents a static view of cellular state. The next frontier will require systematic charting of how subcellular organization changes during natural cellular dynamics and in response to genetic, chemical, or environmental interventions. This would provide a foundation for cell models that connect molecular organization to cellular behavior.∎

## 4 Methods

### 4.1 protdata

*protdata* is a standalone Python package that provides functions to convert the quantitative output of popular proteomics search engines into AnnData format that *grassp* can use. *protdata* stores samples as rows and proteins as columns, following the single-cell convention, and the grassp.io readers transpose its output to the protein-by-sample orientation used by *grassp* (Section 4.2) and set missing values to zero. Supported search-engine outputs are MaxQuant [72], FragPipe [73], DIANN [74], Spectronaut [75], and mzTab [76]. It is designed to distinguish quantitative columns from metadata and store them in the appropriate AnnData slots. During the preparation of this manuscript, AlphaPeptTools [28] was released as an independent scverse-native framework for MS-based proteomics, reflecting the growing adoption of the scverse ecosystem for proteomics. Because both packages produce AnnData objects, the search-engine readers of its alphabase component can be used in place of *protdata* without any changes to the downstream *grassp* workflow. We provide a tutorial demonstrating this interchangeable use in the *grassp* documentation.

### 4.2 *grassp* architecture

The *grassp* architecture is modeled after the popular scanpy [77] package. Like scanpy, *grassp* is a collection of functions that can manipulate an AnnData [26] object. Placing *grassp* inside the scverse [27] ecosystem in this way means that a fractionation experiment is represented in the same container as single-cell data, and that *grassp* functions can be interleaved freely with scanpy. One notable caveat is the orientation of the data. While single-cell data are typically stored with cells as rows and genes as columns, MS-SP data in *grassp* are oriented differently, with proteins as rows and fractions or samples as columns. This orientation was chosen because proteins are the primary observations in MS-SP experiments and researchers are interested in protein–protein relationships rather than relationships between fractions or samples. This transpose makes scanpy functionality directly applicable to MS-SP. For example, a nearest-neighbor graph, Leiden clustering, or a UMAP embedding computed by scanpy operates on proteins in profile space.

*grassp*’s functions are organized into five submodules that mirror scanpy’s namespaces. `grassp.io (gr.io)` reads the output of common MS search engines through *protdata* (see Section 4.1), and additionally reads the `.rda` objects distributed by pRolocdata [15], so that users can access published datasets without installing R. `grassp.preprocessing (gr.pp)` covers quality-control metrics, filtering of proteins, samples, and replicates, protein and sample aggregation, profile normalization, contaminant removal, imputation, the calculation of subcellular enrichment profiles for pulldown- and fractionation-based protocols, and marker annotation from published marker sets and from UniProt [37]. `grassp.tools (gr.tl)` contains MS-SP analysis methods: graph-based and Markov clustering, ontology-aware merging of clusters, semi-supervised localization assignment (independent diffusion and competitive diffusion), and SVMs. It contains Python reimplementations of the TAGM-MAP [17] algorithm, model-based gene set analysis (MGSA) [78], and `QSep` [20] for assessing cluster separation. Additionally, it includes functions for the integration of several datasets into a common embedding [3]. `grassp.plotting (gr.pl)` provides visualization functions, including profile and marker plots, embeddings shaded by classification confidence, resolution heatmaps, ternary diagrams, and relocalization Sankey diagrams. `grassp.datasets (gr.ds)` exposes published reference datasets, programmatic access to the *grassp* data portal and to pRolocdata, and the marker and ontology metadata used for annotation.

Results are written back into the object under documented slots rather than returned as detached tables, following scanpy’s in-place, `key_added` convention: categorical assignments become .obs columns, per-protein by per-compartment score matrices become .obsm entries stored as named pandas DataFrames, graphs are kept in .obsp, and parameters of function calls are recorded in .uns. Because the annotations, embeddings, and quantitation stay in one container, an analysis can be serialized to a single h5ad file at any point and resumed, shared, or handed to another framework. Unless stated otherwise, the analyses of this study were run with *grassp* 0.8.0, *protdata* 0.1.4, scanpy 1.12.4, and AnnData 0.13.3 under Python 3.12.

### 4.3 *grassp* interoperability

To ensure the correctness of the TAGM-MAP reimplementation (`grassp.tl.tagm_map_train` and `grassp.tl.tagm_map_predict`), we compared it to outputs of the native pRoloc implementation (`tagmMapTrain` and `tagmMapPredict`, version 1.52.0) on eight datasets analyzed by Crook et al. [17], obtained from pRoloc-data, reduced to proteins without missing values, and run with identical defaults (100 expectation–maximization iterations, empirical priors). Across the 17,668 unlabeled proteins, both implementations allocated every protein to the same compartment, and the largest difference in any allocation or outlier probability was 4 × 10^−7^, consistent with floating-point accumulation over the iterations. The comparison code is available in the analysis repository (https://doi.org/10.5281/zenodo.24484247, `notebooks/figures/supplement/figure_S7` and `scripts/run_proloc_tagm.R`).

C-COMPASS [25] predicts, for every protein, a class-contribution vector over compartments that sums to one and is the basis of its multi-localization calls. Its published implementation is intended to be used through a graphical user interface that also includes limited preprocessing and visualization capabilities, but the learning core is reachable as a Python library. *grassp* provides a wrapper function around the C-COMPASS function `ccompass.MOP.multi_organelle_prediction`, writing the results back into standard AnnData slots. The compartment calls of C-COMPASS are therefore stored in the same format as those of *grassp*’s native annotators, and can be plotted and evaluated using *grassp* functions.

To ensure correct implementation, the compartment classifications through the *grassp* wrapper function were compared to labels from Haas et al. [25]. Fractionation profiles of the three replicates of the humanized-liver dataset (3 replicates × 14 fractions) were reconstructed from Supplementary Table 8 of Haas et al. [25] (sheet HumLiver DataReport), restricted to proteins identified in at least two replicates, and normalized as described in the original paper (missing values set to zero, then perreplicate sum-to-one profiles, with the min–max scaling of the wrapper switched off). They were classified with the *grassp* function grassp.tl.ccompass, trained on the marker labels of the original study (one network per replicate, 3 rounds × 10 runs, reliability-filtered class contributions, fCC). Each protein was then compared to the authors’ own published C-COMPASS output (Haas et al. [25], Supplementary Table 8, sheet O1(stats) chow: MainCompartment and CC values) over the 11 shared compartments. In total, 7212 proteins were assigned by both pipelines. The classifier is stochastic, so it was run 10 times independently. The main compartment matched the published one for a median of 95.0% of proteins (range 94.6–95.6% across the 10 runs). This is the same as the agreement between two independent runs of the wrapper (median 95.2%, range 94.5–96.0% across the 45 pairs of runs), so the remaining disagreement reflects the variability of the classifier rather than a difference between the two pipelines. The comparison code is available in the analysis repository (https://doi.org/10.5281/zenodo.24484247, `notebooks/figures/supplement/figure_S4`).

pRoloc [15] is an established R/Bioconductor framework for MS-SP data, built on the MSnbase `MSnSet` class [30]. To enable interoperability with the pRoloc ecosystem in R, we provide a standalone R package called *grasspio* that converts between the on-disk AnnData (`h5ad)` format and the `MSnSet` class in both directions (`grassp_as_MSnSet` and `grassp_write_MSnSet`) using anndataR [79].

### 4.4 Semi-supervised subcellular annotation with *grassp*

This section describes how MS-SP maps can be annotated with *grassp*, starting from either a curated set of labeled marker proteins (competitive diffusion) or a compartment vocabulary (independent diffusion).

We use the following notation throughout. An MS-SP experiment is a matrix **X** ∈ ℝ*^n^*^×^*^p^* of *n* proteins by *p* fractions/samples, whose rows **x***_i_* are the normalized abundance profiles of each measured protein. Proteins are annotated over a label space of *m* subcellular compartments: a subset *ℳ* ⊆ {1*, . . . , n*} carries prior annotation, and *U* = {1*, . . . , n*} \ *ℳ* denotes the unannotated remainder. These annotations are represented as a one-hot encoded matrix **Y**^(0)^ ∈ {0, 1}*^n^*^×^*^m^* with *Y_ic_*^(0)^ = 1 if protein *i* is annotated with compartment *c* and 0 otherwise.

The two kinds of prior annotation used in this work differ in the structure of these rows. Curated marker sets assign a single compartment per protein, so their rows sum to one:

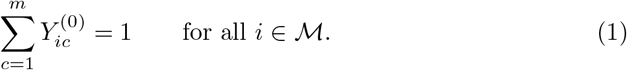

Compartment vocabularies, such as COMPARTMENTS [80], the Gene Ontology cellular component [36], or the UniProt subcellular-location terms [37], instead may carry several entries per protein:

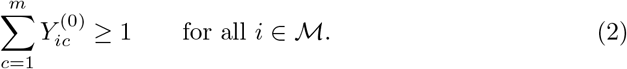

Overlaps can occur because of proteins with multiple localizations or because of the hierarchy of the ontology annotating proteins at multiple resolutions.

Annotation methods return a score matrix **Ŷ** ∈ [0, 1]*^n^*^×^*^m^*, whose entry *Ŷ_ic_* estimates the probability that protein *i* resides in compartment *c*. Whether the constraint Σ*_c_ Ŷ_ic_* = 1 is enforced in the output matrix **Ŷ** depends on whether multi-localization is allowed.

Throughout this work, AUPRC is the stepwise average precision (scikit-learn [81]). For a score vector **s** and binary labels **y**, proteins are ranked by decreasing score, with tied scores grouped, and

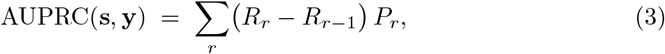

where *P_r_*and *R_r_* are the precision and recall among the *r* highest-ranked proteins and *R*_0_ = 0.

#### 4.4.1 Competitive label diffusion

Semi-supervised annotation in *grassp* (`grassp.tl.competitive_diffusion`) is an implementation of label propagation, a semi-supervised learning algorithm proposed in [82] and [34], adapted to MS-SP by the addition of class balancing and by operating on the same protein-neighborhood graph used for clustering and embedding.

First, a protein-neighborhood graph based on protein profile similarities is constructed using the UMAP [32] algorithm implemented in scanpy’s `scanpy.pp.neighbors` function. This graph is represented as a sparse, symmetric, non-negative affinity matrix **W** ∈ [0, 1]*^n^*^×^*^n^* with zero diagonal and degree matrix **D** = diag(**W1**). Importantly, while users define a neighborhood size *k* to compute this matrix, the actual number of edges from each node is determined by the local density of points in the feature space. We used *k* = 20 for all single-map analyses in this work and found it a reasonable choice for most MS-SP datasets. This graph is then used to diffuse marker labels to unannotated proteins in one of three ways described below.

In a single step of propagating labels, the seed matrix **Y**^(0)^ is diffused once along the graph,

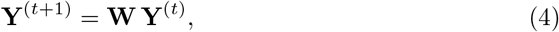

so that each protein accumulates the annotation mass of its immediate neighbors, weighted by affinity. The weights are then row-normalized so that the class probabilities sum to 1,

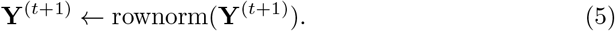

When only a single step is run, this normalization is applied once, after the class balancing described below. Proteins that receive no propagated mass are assigned equal probability for every compartment. Iterating this step propagates mass further along the graph. This iteration can be repeated for a fixed number of steps or until **Y**^(^*^t^*^)^ converges. Labeled data can be clamped to their original labels after each iteration (`fix_markers=True`, the default),

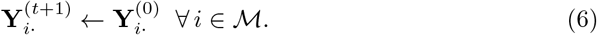

This is effectively the label propagation algorithm proposed in [82], but using the affinity matrix **W** instead of the proposed transition matrix.

Zhou et al. [34] introduce a regularization parameter *α* that replaces the hard clamping with a soft constraint, and spread labels over the symmetrically normalized affinity matrix

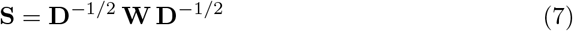

rather than over **W** itself,

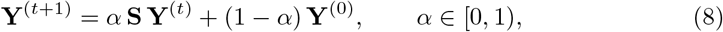

so that the label of a marker can be outweighed by its neighborhood during diffusion, which makes the result more robust to mislabeled markers. We recommend this algorithm for most users and used it for all analyses in this work. It is the default of `grassp.tl.competitive_diffusion` (`method=“spreading”`, with *α* = 0.8). Unlike propagation, spreading does not renormalize **Y**^(^*^t^*^)^ between steps, so the normalization in Eq. 7 keeps the iteration bounded. The affinities in **W** are neither row-stochastic nor spectrally normalized (the largest eigenvalue of **W** lies between 7.3 and 33 on the 12 maps of the annotation benchmark, Section 4.4.4), so iterating with **W** would diverge for most *α*. The eigenvalues of **S** instead lie in [−1, 1], so for *α <* 1 the iteration converges to the unique fixed point

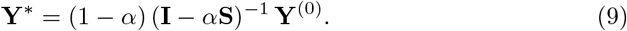

Proteins without edges have zero degree and receive a zero row and column in **S**.

Label spreading iterates until the change in the propagated matrix, relative to the injected seed, falls below a tolerance *ɛ*,

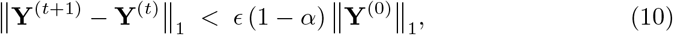

with *ɛ* = 10^−4^. Iterated propagation, which has no *α*, instead stops when ‖**Y**^(^*^t^*^+1)^ − **Y**^(^*^t^*^)^‖_1_ *< ɛ* ‖**Y**^(^*^t^*^+1)^‖_1_. In competitive diffusion both are capped at 1000 iterations by default.

##### Label assignment and class mass normalization

To obtain final labels for unannotated proteins, the propagated mass is typically converted into hard assignments by taking the argmax over compartments. These assignments can then be filtered by the largest probability. In some cases, the number of marker proteins per compartment carries useful prior information about the size of the compartment. *grassp* therefore applies a post hoc step by default (`renormalize_class_mass=True`) that renormalizes the propagated mass across compartments to match the marker counts. This step, which we refer to as class balancing, is the class mass normalization of [82], with the marker counts supplying the target class proportions. Let

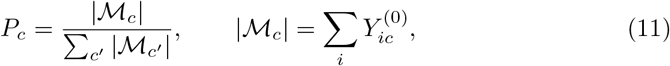

denote the target proportion of compartment *c*, where |*ℳ_c_*| is the number of markers seeding it, and let *z_c_* = Σ*_i_ Y_ic_* be its class mass, the total propagated mass currently assigned to *c*. Each column is rescaled so that its mass matches the target proportion, after which the rows are renormalized to probabilities,

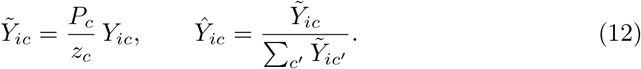

After rescaling, the class masses are proportional to the marker counts, Σ*_i_ Ỹ_ic_* = *P_c_* ∝ |M*ℳ*|. Then labels can be assigned as above.

##### Relationship to k-nearest-neighbor classification

The established alternative in the field is *k*-nearest-neighbor (*k*-NN) classification, which assigns each protein the majority vote of the *k* nearest *markers* in profile space, reporting the fraction of votes for the winning class as its score. Single-step propagation (Eq. 4) is closely related to that algorithm, especially with a binary adjacency matrix as **W**. They differ in two respects. First, the vote is taken over *all* neighbors rather than only over markers, and is weighted by affinity, so that closer neighbors have more voting power than distant ones and confidences are continuous rather than multiples of 1*/k*. Second, because unannotated proteins participate, annotation mass can travel through them when the propagation is iterated, whereas *k*-NN is strictly local: a protein whose neighborhood contains no marker is nevertheless assigned the class of the *k* nearest markers in the entire dataset. This means that arbitrarily distant evidence is indistinguishable in the score from a genuinely local vote.

#### 4.4.2 Independent diffusion

Competitive diffusion (Section 4.4.1), like most published annotation methods, treats compartments as mutually exclusive: every row of **Ŷ** is normalized to a simplex, so evidence for one compartment is evidence against every other. This encodes an implicit assumption that a reliably localized protein can never have more than one compartment with high probability. Two common scenarios violate this assumption. First, some proteins genuinely occupy several compartments, e.g., tRNA synthetases are active in both the cytoplasm and the mitochondrion. Second, many compartment vocabularies annotate proteins at several resolutions at once, e.g., a nucleolar protein is also a nuclear protein.

Expert-curated gene-set libraries (e.g., Gene Ontology cellular component (GO CC) [36], COMPARTMENTS [80], and UniProt subcellular locations [37]) encode comprehensive prior knowledge of protein–compartment associations and typically cover many more proteins than curated marker panels, but they are noisier and organized hierarchically. Independent diffusion (`grassp.tl.independent_diffusion`) annotates a map from such a vocabulary. It diffuses every term of the vocabulary separately over the protein-neighborhood graph of the map. It does not constrain the resulting memberships to a simplex and overwrites prior annotations where the graph neighborhood contradicts them. It then decides which label to report in an explicit postprocessing step.

##### Overview and notation

The seed **Y**^(0)^ is the multi-hot membership matrix of the vocabulary restricted to the proteins in the map, and the output **Ŷ** ∈ [0, 1]*^n^*^×^*^m^* holds the final membership probability per protein and term. Vocabulary members are matched to the map by gene symbol. When several entries of a map share a gene symbol (e.g., isoforms), only the first carries the annotation and the others are treated as unannotated, so that a protein cannot support its own annotation through a duplicate entry. After this matching, *ℳ_c_* = {*i* : *Y_ic_*^(0)^ = 1} denotes the members of term *c* in the map, and *ℳ* = ∪*_c_ℳ_c_* the annotated proteins.

The algorithm consists of five steps, each of which turns one quantity into the next:

1. Per-term diffusion spreads each column of the seed over the graph, which gives the diffused mass *y_ic_*^(^*^T^*).
2. Correction of the diffused mass removes the influence of graph density and of a protein’s own seed, which gives the corrected score *s_ic_*.
3. Selection of the diffusion depth chooses, for each term, the depth *α*_*c*_*^⋆^* at which the corrected score is evaluated.
4. Calibration standardizes *s_ic_* to a *z*-score *z_ic_* and maps it to the membership probability *Ŷ_ic_*.
5. Hierarchy-aware resolution selects, for each protein, the set of active terms *A_i_^⋆^* from which the reported labels are read.

##### Per-term diffusion

Independent diffusion uses the same normalized operator **S** (Eq. 7) as label spreading in competitive diffusion (see Section 4.4.1). Each column **y**_*c*_ = **Y***_c_*^(0)^ is spread separately,

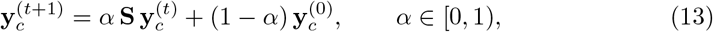

until the convergence criterion of Eq. 10, applied to the single column, is met, yielding **y***_c_*^(^*^T^* ^)^. The parameter *α* controls the diffusion depth. It is selected automatically for each term in the third step.

##### Correction of the diffused mass

Let **u**^(^*^T^* ^)^ denote the result of diffusing the indicator of the annotated proteins, *u_i_*^(0)^ = 1 for *i* ∈ *ℳ* and *u_i_*^(0)^ = 0 otherwise, with the same *α*. The corrected score is

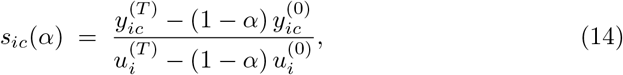

the fraction of the diffused annotated neighborhood of protein *i*, excluding its own seed, that carries term *c*.

Both parts of the correction are needed because the raw diffused mass is not directly interpretable. Without competition, the mass a protein receives depends on the density of its graph neighborhood, and as *α* → 1 the diffused mass becomes independent of seed placement. The denominator removes this confound by dividing by the mass the protein would have received had every annotated protein carried the specific label. In addition, every seed protein re-injects a fraction (1 − *α*) of its own label at each step, so a protein would partly read back its own prior annotation, which we want to avoid to keep annotations data-driven. The terms subtracted in the numerator and the denominator remove that direct self-contribution. Only the small part of a protein’s own seed that returns to it along closed walks of two or more steps remains.

##### Per-term diffusion depth

The diffusion depth *α* is selected separately for each term to accommodate differences in term size and compactness. For each term, *α* is chosen from a grid A (by default 19 values evenly spaced from 0.1 to 0.9) to maximize the AUPRC (Eq. 3) with which the corrected score ranks the term’s own members above the other annotated proteins,

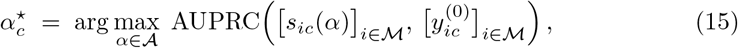

with ties broken toward the smaller *α*. Because the direct self-contribution is removed in Eq. 14, a member’s score rests on its neighbors, so this selection cannot be won by a depth that merely returns the seed. Terms with fewer than five members in the map, the number of cross-fitting folds used below, are too unstable for this estimation and are dropped, which leaves *m*^′^ terms. Hereafter *s_ic_* = *s_ic_*(*α_c_^⋆^*).

##### Calibration

Scores of terms of different sizes are not comparable: compartment terms with more members will diffuse more mass to a random protein even with uniform distribution in the graph. Scores are therefore converted to probabilities in two steps, a standardization against a null model followed by a monotone mapping to probabilities.

First, each score is standardized against the null in which the term’s members are scattered at random over the annotated proteins of the graph. Under that null, *s_ic_* is a weighted average, over the effective annotated neighborhood of protein *i*, of Bernoulli draws with rate *p_c_* = |*ℳ_c_*|*/*|*ℳ*|, the fraction of annotated proteins that carry the term, giving

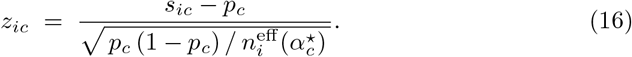

Here *n_i_*^eff^ (*α*) is the Kish effective neighborhood size [83] of the diffusion kernel at depth *α*, **K***_α_* = (1 − *α*)(**I** − *α***S**)^−1^, which maps a seed to its fixed point (Eq. 9), with the self term excluded and only annotated proteins counted,

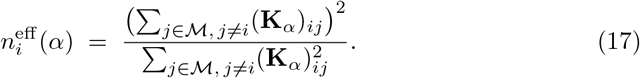

It counts how many annotated neighbors a protein effectively sees once their weights are taken into account, and therefore how much evidence its score rests on. Eq. 16 places all terms, whatever their size, and all proteins, whatever their local density, on one scale.

Second, *z* is mapped to a probability by isotonic regression [84] against the observed membership of the annotated proteins. Two monotone curves are fitted. A per-term curve *g_c_* uses only that term’s own (*z_ic_, Y_ic_*^(0)^) pairs. A size-conditional curve *h* pools terms of similar size: the *m*^′^ retained terms are binned into *Q* = min(6, ⌊*m*^′^*/*8⌋) quantile strata of log |*ℳ_c_*| (at least one stratum), one isotonic curve is fitted on the pooled pairs of each stratum, and *h* for a given term interpolates linearly in log |*ℳ_c_*| between the curves of the two strata whose median values of log |*ℳ_c_*| bracket it. Terms smaller or larger than every stratum median take the curve of the nearest stratum. The two curves are blended with a weight that reflects how much data the term itself provides,

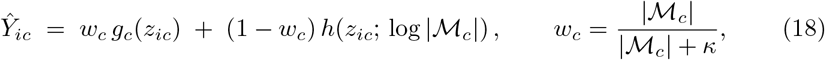

where *κ* is a shrinkage constant (default *κ* = 30). Large terms are thus calibrated more heavily on their own evidence, while small terms borrow strength from same-sized peers rather than overfitting a handful of members. All curves are cross-fitted over five folds of the proteins, so no protein’s probability is read from a curve fitted on itself, and unannotated proteins never serve as negative examples. Simpler calibrations (per term only, pooled across all terms, or none) are available as options.

##### Hierarchy-aware resolution

**Ŷ** holds independent memberships, which means that simply annotating a protein with the highest-scoring term is not sufficient: a protein in the nucleolus should score highly for nucleolus *and* nucleus. When several nested terms have high probability, the most specific one should be chosen. If several non-nested terms have high probability, all of them should be used in the annotation. The resolver therefore selects, for each protein, the smallest set of active terms that explains its whole membership profile, in three stages.

###### Nesting of terms

The nesting of terms is described by the containment matrix *C*, where

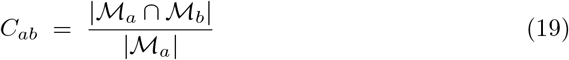

is the fraction of the members of term *a* present in the map that also belong to term *b*. Term *a* is treated as nested within *b* when *C_ab_* ≥ *ν* (`nest_threshold`, default *ν* = 0.8).

###### Selection of the active set

For protein *i*, the candidate terms *T_i_* are those with *Ŷ_ic_* ≥ *τ* (default *τ* = 0.4). If no term passes the threshold, the single highest-scoring term is the only candidate. If more than 12 terms pass it, only the 12 with the highest membership are kept as candidates (cap). A hypothesized active set *A* ⊆ *T_i_* predicts that candidate *b* is observed with probability 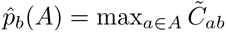, clipped to [*δ,* 1 − *δ*] with *δ* = 0.02, where 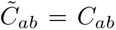 if *a* is nested within *b* and 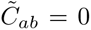 otherwise: if *a* is truly active and is nested within *b*, then *b* must be observed too. The active set is chosen by penalized Bernoulli likelihood over all subsets of at most *a*_max_ candidates (default *a*_max_ = 3), including the empty set,

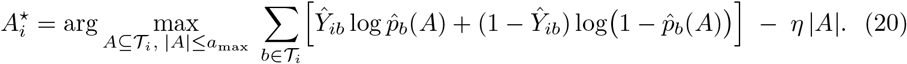

The penalty *η*|*A*| (default *η* = 1) makes the procedure prefer the smallest set of compartments that accounts for the whole observed profile: a high nucleus score is *explained away* by an active nucleolus rather than counted as separate evidence.

###### Reported labels

The reported label is chosen among the leaves of *A_i_^⋆^*, i.e., the terms of *A_i_^⋆^* within which no other term of *A_i_^⋆^* is nested (where two terms are nested within each other, the smaller one is treated as the nested term). It is the leaf whose removal from *A_i_^⋆^* lowers the likelihood term of Eq. 20 the most, i.e., the leaf that explains the largest part of the protein’s membership profile. A protein is left unlabeled if *A_i_^⋆^* is empty. Alongside this single call, *grassp* reports a multi-localization label that lists all leaves of *A_i_^⋆^*, ordered by calibrated membership.

##### Relationship to existing methods

Predicting overlapping, hierarchical labels one term at a time by network propagation is an established strategy in gene function prediction [39]. GeneMANIA [38], for example, solves a Gaussian random field label propagation problem independently for every Gene Ontology term over a gene association network. Independent diffusion follows this one-vs-rest design but adapts it to the protein-neighborhood graph of a single map and to the downstream need for a calibrated localization call. GeneMANIA handles the related problem of class imbalance before propagation, by seeding non-members with −1 and unlabeled genes with the mean label [38]. We instead seed only the members and account for the base rate of each term after propagation, in the calibration step, so that the corrected score can be compared against an explicit null. That null (Eq. 16) is the label-permutation null underlying the parametric *z*-score of Picart-Armada et al. [85], who showed that such standardization corrects the topology-driven bias of raw propagation scores.

In protein function prediction, the task of turning independent per-term predictions into a consistent set is known as reconciliation and is usually solved by projecting the probabilities onto the hierarchy so that no term exceeds its ancestors [86]. Subcellular annotation additionally requires a single most specific call per protein. The resolver therefore takes the generative view of MGSA [78], which explains a set of observed genes by a small number of hidden active categories and lets overlapping categories explain each other away. Here the roles are transposed to one protein at a time: the observations are the protein’s calibrated term memberships, and the hidden variables are which terms are truly active for that protein. The active set plays the role of the hidden term states, the containment-based 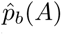 replaces the noisy- OR that links active categories to observed genes, the clipping constant *δ* plays the role of the false-positive and false-negative rates, and the penalty is the log prior odds of activation, *η* = log (1 − *p*)*/p*, so that *η* = 1 corresponds to a prior activation probability of *p* ≈ 0.27. Because each protein has at most 12 candidate terms, of which at most *a*_max_ are active, the posterior mode is found by exhaustive enumeration of the active sets rather than by the Markov chain Monte Carlo sampling that MGSA requires over thousands of categories.

#### 4.4.3 Benchmark of competitive diffusion on the C-COMPASS simulated dataset

We benchmarked the competitive diffusion algorithm against C-COMPASS [25], BANDLE [19], and DOM-ABC [13] on the simulated fractionation dataset introduced with C-COMPASS (Supplementary Fig. S2). The three existing tools’ predictions, and the 854-protein marker set they were given, were taken from the original publication [25].

For competitive diffusion, replicate profiles were averaged across replicates, scaled to sum to one, and filtered to proteins quantified in at least two replicates (4029 proteins), resulting in the same input the published tools received. The published markers were then propagated over the protein-neighborhood graph (*k* = 20) by label spreading with default parameters. The two simulated conditions were propagated independently.

Classification was evaluated on the 1139 single-localization non-marker proteins defined by Haas et al. [25], with precision, recall, *F*_1_, and AUPRC macro-averaged over compartments. Localization error was computed as half the total absolute difference between compartment weights/probabilities,

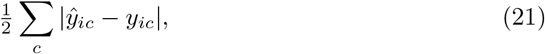

where *ŷ_ic_*and *y_ic_* are the predicted and true probabilities for protein *i* in compartment *c*.

Relocalization error was calculated with the same statistic applied to the change between conditions, 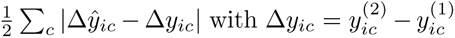. The predicted change was the difference between the outputs for the two conditions for BANDLE and competitive diffusion, and the published relocalization vector for C-COMPASS. DOMABC was not scored because the published table contains its output for condition 1 only. Unlike in the original study, no confidence cutoff was applied to the relocalization output of any method. This ensured a fair comparison across methods. We additionally scored each tool’s ability to predict relocalization, using the AUROC. For this we classified proteins as relocalized or static depending on whether their true compartment assignment was the same between conditions. We also computed the AUROC separately for each true relocalization magnitude, in each case against the same unchanged proteins, with the magnitude calculated as

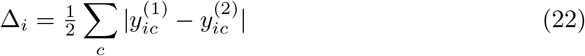

for each protein *i*. We then calculated the AUROC, on the 3094 proteins for which all statistics were available, for the following predicted relocalization statistics: BANDLE’s differential localization probability, C-COMPASS’s relocalization score and its negated *P* value, and the summed absolute change in propagated probability. Confidence intervals were calculated as the 2.5th–97.5th percentiles of 500 class-stratified bootstrap resamples over proteins.

#### 4.4.4 Benchmarking competitive diffusion and independent diffusion on real datasets

##### Datasets

Both annotation strategies were benchmarked on 12 published human maps from the *grassp* collection, chosen to span a variety of cell lines, enrichment protocols, and quantification strategies (Supplementary Table S1). For preprocessing steps, see Section 4.9. Each map was annotated with the labels reported by the source study and the set of marker proteins used in Hein et al. [3] (see Section 4.9.6). Every map was annotated using independent diffusion and competitive diffusion with the same parameters.

##### Annotation strategies

Three strategies were compared on each map. The *author annotation* is the label column reported by the source study, carrying no probabilities. *Competitive diffusion* (`grassp.tl.competitive_diffusion`) was seeded with the literature-curated marker set of Hein et al. [3] (16 compartments) and run as label spreading (Eq. 8, *α* = 0.8, with class balancing) with default parameters, except that markers were not clamped to their seed labels (`fix markers=False`), so that marker proteins were scored on their diffused probabilities. No probability cutoff was applied, so every protein received a label. *Independent diffusion* (`grassp.tl.independent_diffusion`) was run as described in Section 4.4.2 against the consolidated UniProt subcellular-location vocabulary (24 terms, see Section 4.9.5), with size-aware calibration, the hierarchy-aware likelihood resolver, default parameters, and a fixed random seed. No parameter was tuned for any dataset.

##### Label harmonization

Because the three compared annotations use different vocabularies, all labels were mapped into a consolidated vocabulary through explicit dictionaries. The Hein et al. classes were mapped one-to-one (e.g., “Cytosol” to “Cytoplasm” and “PM” to “Cell membrane”). Author labels were normalized (lower-cased, whitespace and punctuation collapsed) and looked up in a curated dictionary covering the vocabularies of all source studies. Labels without an unambiguous consolidated counterpart (e.g., “unclassified” or “protein complex”) were treated as unannotated, and multi-label author entries were reduced to their first term. The two cytosolic ribosomal-subunit terms of the consolidated vocabulary and the unsplit “Ribosome” marker class were collapsed into a single ribosome class for scoring. Probability matrices were mapped with the same dictionaries, summing columns that mapped to the same consolidated term. The complete mapping tables are part of the analysis code (`notebooks/_shared/author_vocab.py` and `notebooks/figure_2/_vocab.py` in the repository listed under Data and Code Availability).

##### Agreement with the published annotation

For Fig. 2f, the two strategies were run on the HEK293T OrgIP map of Hein et al. [3] as described above. Agreement is the fraction of proteins whose harmonized primary label equals the harmonized author label, among the proteins labeled by both (7584 for competitive diffusion and 7178 for independent diffusion).

##### Validation against external mitochondrial and membrane annotations

For Fig. 2g, external reference annotations were attached with `grassp.pp.add_external_validation_markers`, which matches UniProt accessions against a table distributed with *grassp* that was assembled from the transmembrane and intramembrane sequence features of reviewed human UniProt entries [37], from MitoCarta3.0 [45], and from MitoCoP [46]. For mitochondrial prediction, positives were proteins listed in both MitoCarta3.0 and MitoCoP (on average 591 per map), negatives were proteins listed in neither, and proteins on only one of the two lists were excluded so that the negative set is clean. The references were used only for scoring and never as input to any annotation strategy. A protein outside both mitochondrial lists that co-fractionates with mitochondrial proteins can therefore still be annotated as mitochondrial, but it counts as a false positive. Proteins were ranked by their harmonized probability for “Mitochondrion”. For membrane-domain prediction, positives were proteins carrying at least one annotated transmembrane or intramembrane segment, and proteins were ranked by their summed probability over the 12 membrane-bounded compartments of the consolidated vocabulary (cell membrane, nucleus membrane, ER, ERGIC, Golgi apparatus, endosome, early endosome, lysosome, mitochondrion, peroxisome, lipid droplet, and cilium). Because the author annotation carries no probabilities, its label was converted to a binary score by case-insensitive keyword rules for the mitochondrion or for the membrane-bounded compartments, which yields a valid single-operating-point AUPRC. All proteins of a map were scored, including the marker proteins that seeded competitive diffusion, mirroring the author annotation, whose own labels are part of the scored set as well. Performance was summarized as the AUPRC per map and reported as mean ± s.d. over the 12 maps. Absence of an annotated membrane segment is not a clean negative, since many membrane-associated proteins lack such a domain. The membrane-domain AUPRC therefore measures the enrichment of membrane features among predicted membrane proteins and should not be interpreted as an absolute measure.

##### Held-out validation against the HOU benchmark

For Fig. 2h,i, we used the HOU benchmark of Wefers et al. [47] as an independent, largely imaging-derived ground truth. The benchmark provides a test set of 3814 human proteins with multi-label compartment annotations validated across the three sources, together with a training set of 26,436 proteins. Both tables were obtained from the authors’ repository (https://github.com/CellProfiling/seq2loc_benchmark, commit b785552), and the finest annotation level (level 1) was used. Proteins were matched by UniProt accession, using the accession of each map entry and, where a map lists further accessions of a protein group, any of them with isoform suffixes removed. A protein matching both tables was assigned to the test set. Between 988 and 3017 matched test proteins per map carried at least one label with a consolidated counterpart and were scored (mean 1910).

To make the comparison genuinely held out, each strategy was allowed to draw on training proteins only. Competitive diffusion was seeded with the Hein et al. markers restricted to training proteins (271–939 markers per map), and independent diffusion was run on gene sets restricted to the gene symbols of training proteins, after additionally removing every gene symbol carried by a test protein so that a test protein could not seed itself through a paralog or isoform sharing its symbol. Both strategies were otherwise run exactly as above. The author annotation does not depend on the split and was scored as is. After harmonization, it covered on average 65% of the test proteins, whereas the two diffusion strategies label every protein.

Test labels were mapped into the consolidated vocabulary with a dictionary as above, retaining every term of a multi-label entry. HOU tokens without a single consolidated counterpart (“vesicles” and “endomembrane system”) were dropped. Three consolidated terms have no HOU counterpart of their own and were folded into their parent term in ground truth and predictions alike (ERGIC into ER, early endosome into endosome, and NPC into nucleus), summing the corresponding probability columns. Scoring was restricted to the compartments that all four vocabularies (HOU, Hein et al. markers, author labels, and consolidated) can express, which left 11 compartments: nucleus, cytoplasm, mitochondrion, cell membrane, ER, cytoskeleton, Golgi apparatus, endosome, centrosome, lysosome, and peroxisome. For each map and compartment we computed the one-vs-rest AUPRC over the map’s test proteins, counting a protein as positive if the compartment was among its test labels and ranking proteins by their harmonized probability for that compartment. The author annotation received a binary probability from its label. The macro AUPRC of Fig. 2h is the mean over the scored compartments of each map, so that rare compartments weigh as much as abundant ones. The per-compartment values are shown with each map as one point (Fig. 2i). The maps are downloaded and annotated, and all metrics computed, by the Fig. 2 scripts of the analysis repository (`notebooks/figure_2`).

#### 4.4.5 Runtime benchmark

To compare the runtime of competitive diffusion and independent diffusion with published classifiers (Supplementary Fig. S1), every method was run with its package defaults unless stated otherwise below, including its default parallelization, on the same input matrices and marker sets on one laptop (Apple M5 Max, 18 cores). Times are wall-clock times from the loaded data object to per-protein predictions. They include graph construction for the graph-based methods and hyperparameter selection where a method performs it.

Competitive diffusion (`grassp.tl.competitive_diffusion`) was timed together with the construction of the protein-neighborhood graph (*k* = 20) and was run as label spreading (*α* = 0.8) with clamped markers. Independent diffusion (`grassp.tl.independent_diffusion`, default settings, likelihood resolver) used the same graph and a vocabulary of gene sets instead of markers. On the experimental maps, the vocabulary was the UniProt subcellular-location gene sets bundled with *grassp*, matched by gene symbol, either the 24 consolidated terms or the full hierarchy of 235 terms, of which terms with fewer than 20 members were not called. The simulated data carry no gene identifiers, so there the marker set of each of the 8 compartments served as the vocabulary, which is not a meaningful use of the method but sufficient to time it. TAGM-MAP was run in *grassp* (`grassp.tl.tagm_map_train` and `grassp.tl.tagm_map_predict`) and in pRoloc (`tagmMapTrain` and `tagmMapPredict`). SVM classification was run in pRoloc with its default grid search (`svmOptimisation`, 100 repetitions of 5-fold cross-validation) followed by `svmClassification`. C-COMPASS was run through `grassp.tl.ccompass` on the biological replicates of each map, with one network per replicate as in its original publication [25]. For the HEK293T OrgIP map, the replicates were taken from the raw portal object (3 replicates of 84 pulldowns, 8448 of the 8450 proteins of the map).

The simulated maps were taken from Haas et al. [25]. The simulation of Supplementary Fig. S2 (4029 proteins, 16 fractions, 8 compartments, 854 markers) was subsampled to 500, 1000, 2000, and 3000 proteins, with markers and unlabeled proteins drawn in proportion and each smaller map a subset of the next. The simulations with 10 to 24 fractions are those of their Supplementary Table 3 (3916–4129 proteins). The mouse embryonic stem cell hyperLOPIT [9], U2-OS hyperLOPIT [48], and HeLa Dynamic Organellar Maps [11] datasets and their markers were obtained from pRolocdata, and the HEK293T OrgIP map [3] from the *grassp* portal with the markers of Hein et al. Competitive diffusion, independent diffusion, and the *grassp* implementation of TAGM-MAP were timed with *grassp* 0.8.0, C-COMPASS with version 2.0.1, and the pRoloc methods with pRoloc 1.52.0 under R 4.6.1.

### 4.5 Organelle immunoprecipitation and proteomic profiling of iNeurons

#### 4.5.1 iNeuron differentiation

KOLF2.1J human iPSCs harboring a stably integrated, doxycycline-inducible NGN2 cassette were differentiated into cortical excitatory neurons (iNeurons) using a protocol adapted from the Kampmann laboratory (UCSF) [87]. All differentiation steps were performed at 37 °C with 5% CO_2_. Three days prior to dissociation (Day −3), T-25 flasks were coated with Matrigel Growth Factor Reduced Basement Membrane Matrix (Corning #356231) diluted 1:100 in Knockout Dulbecco’s Modified Eagle Medium (DMEM, Thermo Fisher Scientific #10829018) and incubated at 37 °C for at least 1 h. iPSCs at 50–80% confluence were dissociated with StemPro Accutase (Thermo Fisher Scientific #A1110501) for 6 min, counted using a Countess cell counter, and seeded at 2.0–2.5 × 10^6^ cells per T-25 flask in N2 pre-differentiation medium consisting of Knockout DMEM/F-12 (Thermo Fisher Scientific #12660012), 1× MEM Non-Essential Amino Acids (NEAA, Thermo Fisher Scientific #11140050), 1× N-2 Supplement (100×, Thermo Fisher Scientific #17502048), 10 ng/mL BDNF (PeproTech #450-02), 10 ng/mL NT-3 (PeproTech #450-03), and 1 µg/mL mouse laminin (Thermo Fisher Scientific #23017015). Doxycycline (2 µg/mL, Sigma-Aldrich #D3447) was added to induce NGN2 expression, and the ROCK inhibitor Y-27632 (10 µM, Tocris #1254) was included to promote survival. Medium was replaced with fresh N2 pre-differentiation medium (without ROCK inhibitor, with doxycycline) on Days −2 and −1. On Day 0, cells were dissociated with Accutase (7 min at 37 °C), counted, and replated at 7.5 × 10^5^ cells per well onto poly-D-lysine (PDL)-coated plates in N2/B27 differentiation medium, prepared as a 1:1 mixture of DMEM/F-12 (Thermo Fisher Scientific #11320033) and Neurobasal-A (Thermo Fisher Scientific #10888022) supplemented with 1× NEAA, 1× GlutaMAX (100×, Gibco #35050061), 1× N-2 Supplement, 1× B-27 Supplement (minus vitamin A, Thermo Fisher Scientific #12587010), 10 ng/mL BDNF, 10 ng/mL NT-3, 1 µg/mL mouse laminin, and 2 µg/mL doxycycline. PDL coating was performed using a 1:1 ratio of PDL in sterile ultrapure water (2 mL per well for 6-well plates) for 2 h at room temperature (RT), followed by two washes with sterile ultrapure water. Half-medium changes with fresh N2/B27 medium (without doxycycline) were performed on Days 7 and 14. Cells were harvested for OrgIP or N/O/C fractionation on Day 21 of maturation.

#### 4.5.2 iNeuron OrgIP bait generation and validation

To generate the endogenously tagged cell lines for OrgIP, we performed cell engineering using homology-directed repair via RNP electroporation with the P3 Primary Cell 96-well-Nucleofector (Lonza, cat. #V4XP-3096) kit following the manufacturer’s protocol. For each reaction, a total of 20 µL master mix was prepared (16.4 µL P3 buffer + 3.6 µL P3 supplement). The RNP complexes were prepared in a separate tube by adding 4 µg HiFi Cas9 nuclease V3 (IDT) and 3.2 µg sgRNA in 10 µL of the P3/supplement master mix, followed by a brief incubation at RT. Meanwhile, 1.6 × 10^5^ iPSCs were pelleted and resuspended in the remaining 10 µL of P3/supplement mix. The cell suspension was added to the RNP mixture, followed by 1 µg of dsDNA donor for a final reaction volume of 20 µL. The sgRNA and donor sequences were as described in Hein et al. [3], except for TOMM70A, which was not tagged there (Supplementary Table S14). Sequential addition of donor DNA following cell-RNP homogenization proved to be an important consideration for ensuring the highest editing efficiency. The suspension was transferred to a 96-well Nucleocuvette plate. Electroporation was performed using the Amaxa 96-well Shuttle Nucleofector device (Lonza) with program CA-137. After electroporation, 80 µL prewarmed StemFlex medium (Gibco, cat. #A3349401) supplemented with 1× CloneR2 (STEMCELL Tech, cat. #100-0691) and Alt-R HDR Enhancer (IDT, cat. #1081072) was immediately added to the cells in the 96-well Nucleocuvette plate. Then, 50 µL of the cell suspension was transferred to a vitronectin-coated 96-well plate containing 150 µL of the same prewarmed recovery medium and cultured at 32 °C overnight. The next day, the medium was replaced with StemFlex supplemented with CloneR2, and the cells were incubated at 32 °C. On day 3, cells were moved to 37 °C in StemFlex containing CloneR2. Cells were then maintained in standard StemFlex medium for further culturing. Once the cells reached 80–90% confluence, fluorescence-activated cell sorting was used to collect the top 1% of the GFP-positive population for most OrgIP lines, except EEA1, PEX3, and GOLGA2, which were sorted for the top 0.5% of GFP-positive cells because of their weaker GFP expression. A Sony cell sorter (model LE-SH800SFP) was used to sort a polyclonal pool of GFP-positive, endogenously tagged iPSCs for each line. These OrgIP lines were then examined for correct tagging by imaging the cells at the iPSC stage on our in-house confocal microscope to verify the correct GFP localization signal. The OrgIP lines were also tested for their ability to differentiate into neurons to make sure gene editing did not affect differentiation. The morphology of iNeurons derived from endogenously tagged iPSCs was compared to that of iNeurons derived from untagged (control) iPSCs to make sure iNeuron differentiation was not affected by genome editing. To make sure organelle labeling remained intact after differentiation into iNeurons, all OrgIP lines were imaged again at the iNeuron stage by confocal microscopy to check for the correct GFP localization signal.

Each OrgIP pulldown experiment included 1–5 endogenously tagged OrgIP iPSC lines (baits). These iPSC lines were seeded on PDL-coated plates in triplicate along with untagged control iPSCs and differentiated into iNeurons using the protocol described above. On Day 21, the iNeurons were checked for correct morphology and cell number to make sure that all tagged and control iPSC lines had differentiated well into iNeurons and were present at comparable density before the cells were lysed for pulldowns. Each bait was profiled in one to three experiments (ATG101, ATP1B3, GOLGA2, RPL36, TOMM70A, and VPS35 in one; CLTA, G3BP1, NCLN, PEX3, RAB11A, and SEC61B in two; EEA1 and LAMP1 in three), giving 24 bait–experiment combinations. Every combination comprised three replicates except one LAMP1 experiment, in which one replicate failed quality control and was excluded, resulting in a total of 71 bait IP samples. Untagged controls were processed in triplicate in each experiment. Two further experiments, one profiling LAMP1 and one profiling LAMP1 and VPS35, lacked matched untagged controls and were excluded before analysis.

#### 4.5.3 N/O/C fractionation

Cells were fractionated into nuclear (N), organellar (O), and cytosolic (C) fractions (N/O/C) as described previously [3], with modifications. KOLF2.1J iPSCs harboring a stably integrated, doxycycline-inducible NGN2 cassette were seeded in 6-well plates and differentiated into iNeurons as described above. One well constituted a single replicate, and each condition was processed in triplicate. N/O/C fractionation was performed on Day 21 of differentiation. Unless stated otherwise, all steps were performed on ice or at 4 °C. Medium was aspirated from the 6-well plate and the wells were washed once with phosphate-buffered saline, followed by addition of 500 µL of homogenization buffer (hypotonic buffer — 25 mM Tris-HCl pH 7.5, 50 mM sucrose, 0.2 mM EGTA, 0.5 mM MgCl_2_ — supplemented with 1× Halt Protease/Phosphatase Inhibitor Cocktail, EDTA-free, 100×, Thermo Fisher Scientific, 78443) to the well. Cells were gently dislodged from the plate by pipetting with a P1000 tip. The resulting suspension was passed through a 25-gauge blunt needle six times to shear the cells. The homogenate was immediately mixed with 44.5 µL of concentrated sucrose buffer (2.5 M sucrose, 0.2 mM EGTA, 0.5 mM MgCl_2_), which restored isotonic conditions (∼250 mM sucrose final). Nuclei were pelleted at 1000 × *g* for 10 min. The supernatant was centrifuged at 20,000 × *g* for 30 min, giving an organelle pellet and a cytosolic supernatant. The nuclear and organelle pellets were resuspended in 48 µL and 32 µL, respectively, of 1× sodium dodecyl sulfate (SDS) buffer (5% SDS, 50 mM Tris pH 8, 2 mM MgCl_2_, 1× Halt protease and phosphatase inhibitors, Thermo Fisher Scientific, 78443). The cytosolic fraction was brought to 1× SDS buffer by adding 5× SDS buffer (125 µL per 500 µL). All fractions were treated with 25 U of benzonase (Millipore Sigma, 706643) for 15 min at RT on a Thermomixer set to 1500 rpm to digest nucleic acids, then clarified by centrifugation at 14,000 × *g* for 10 min at RT and stored at −80 °C. Fraction yields were quantified by BCA assay according to the manufacturer’s instructions (Pierce, Thermo Fisher Scientific, 23250), and fraction purity was confirmed by western blotting for histone H3 (nuclear), cathepsin D and COX IV (organelle), and *α*-tubulin and pyruvate kinase (cytosol).

Protein (10 µg per fraction) was processed using S-Trap micro spin columns according to the manufacturer’s instructions (ProtiFi, Fairport, NY, USA, K002-MICRO-0160KT) with the following modifications. Samples were reduced with dithiothreitol (DTT) at a final concentration of 1 mM and alkylated with iodoacetamide at a final concentration of 5 mM in the dark. Both steps were incubated at RT for 20 min on a Thermomixer set to 1500 rpm. Protein digestion was performed with 0.5 µg of trypsin and 1 mAU of LysC per sample. All samples were then dried down by centrifugal evaporation.

#### 4.5.4 OrgIP procedure

OrgIP was performed as previously described [3] with the following modifications for neuronal cultures. Neurons were grown in 6-well plates, with one well constituting a single replicate, and each condition was processed in triplicate. Because neurons detach readily from the plate, cells were harvested without scraping: following aspiration of the medium, wells were washed once with phosphate-buffered saline, and 500 µL of homogenization buffer (hypotonic buffer — 25 mM Tris-HCl pH 7.5, 50 mM sucrose, 0.2 mM EGTA, 0.5 mM MgCl_2_ — supplemented with 1× Halt Protease/Phosphatase Inhibitor Cocktail, EDTA-free, 100×, Thermo Fisher Scientific, 78443) was added directly to the well (rather than pelleting cells by centrifugation and resuspending in buffer, as for HEK293T cells). Cells were gently dislodged from the plate by pipetting with a P1000 tip. The resulting suspension was passed through a 25-gauge blunt needle six times (rather than the 23-gauge needle and four strokes used for HEK293T cells) to shear the cells. Homogenates were then mixed with sucrose buffer (2.5 M sucrose, 0.2 mM EGTA, 0.5 mM MgCl_2_) to restore the isotonic conditions and centrifuged at 1000 × *g* for 10 min at 4 °C to pellet nuclei, as previously described [3]. Supernatants were incubated with 10 µL (rather than 20 µL) of anti-HA magnetic beads (Thermo Scientific Pierce, PI88836) per replicate, and subsequent bead washes were performed as previously described [3]. On-bead digestion was performed as previously described [3]: beads were resuspended in 1× urea buffer (8 M urea, 50 mM Tris-HCl, 4 mM DTT, Pierce™ DTT NoWeigh™, Thermo Fisher Scientific, A39255) and alkylated with iodoacetamide (Pierce™ Iodoacetamide Single-Use, Thermo Fisher Scientific, A39271) at a final concentration of 5 mM for 30 min at 22 °C with shaking, protected from light. Proteins were then digested with 0.5 µg LysC (Wako Chemicals USA, Fisher Scientific, NC9223464) overnight at RT with shaking, followed by 1 µg trypsin (Pierce™ MS Grade Trypsin Protease, Thermo Scientific, PI90058) for 4 h at RT with shaking. Beads were removed on a magnet, and the peptide-containing supernatant was acidified with trifluoroacetic acid (TFA) to a final concentration of 1% and stored at −80 °C until MS analysis.

#### 4.5.5 Peptide desalting

Peptides were acidified to a final concentration of 1% TFA, followed by desalting on StageTips [88] packed with styrenedivinylbenzene reverse-phase sulfonate (SDB-RPS) material (CDS Analytical, Oxford, PA, USA, 70-2019-1004-9EA). StageTips were activated with 100% methanol, conditioned with 80% acetonitrile/0.1% TFA, and equilibrated with 0.2% TFA. Samples were then loaded and washed sequentially with 99.9% isopropanol/0.1% TFA, 0.2% TFA, and 0.1% formic acid. OrgIP peptides were eluted with 60% acetonitrile/0.5% ammonium hydroxide. To achieve proteomic depth, the N/O/C peptides were triple-fractionated [89] by eluting first with 40% acetonitrile/0.5% formic acid/100 mM ammonium formate, then with 60% acetonitrile/0.5% formic acid/150 mM ammonium formate, and finally with 80% acetonitrile/1% ammonium hydroxide. Desalted peptides were flash frozen, dried by centrifugal evaporation, and resuspended in 2% acetonitrile/0.1% TFA.

#### 4.5.6 Liquid chromatography and MS acquisition

##### N/O/C

Peptides were analyzed on a Fusion Lumos mass spectrometer (Thermo Fisher Scientific, San Jose, CA) equipped with a Vanquish Neo liquid chromatography system (Thermo Fisher Scientific, San Jose, CA) configured for direct injection with a nanoFlex electrospray ionization source. Peptides were separated by capillary reverse-phase chromatography on a 25 cm column (75 µm inner diameter, packed with 1.6 µm C18 resin, AUR2-25075C18A, IonOpticks, Victoria, Australia). The electrospray ionization voltage was set to 2000 V. Peptides were introduced into the mass spectrometer using a two-step linear gradient of 3–27% buffer B (0.1% (v/v) formic acid in 80% (v/v) acetonitrile) for 52.5 min followed by 27–40% buffer B for 14.5 min, at a flow rate of 300 nL/min. The column temperature was maintained at 50 °C throughout the procedure. Data were acquired in top-speed data-dependent acquisition (DDA) mode with a duty cycle time of 1 s. Full MS scans were acquired in the Orbitrap mass analyzer (FTMS) with a resolution of 120,000 (full width at half maximum, FWHM) and an m/z scan range of 375–1500. Selected precursor ions were subjected to fragmentation using higher-energy collisional dissociation (HCD) with a quadrupole isolation window of 0.7 m/z and a normalized collision energy of 31%. HCD fragments were analyzed in the ion trap mass analyzer (ITMS) set to Turbo scan rate. Fragmented ions were dynamically excluded from further selection for a period of 60 s. The automatic gain control (AGC) target was set to 1,000,000 and 10,000 for full FTMS and ITMS scans, respectively. The maximum injection time was set to Auto for both FTMS and ITMS scans.

##### OrgIP

Of the 10 µL of desalted peptides, 1 µL was analyzed on a timsTOF SCP mass spectrometer (Bruker Corporation) coupled to an EASY-nLC 1200 liquid chromatography system (Thermo Fisher Scientific). Peptides were separated by reverse-phase chromatography on a 25 cm column (75 µm inner diameter, packed with 1.6 µm C18 resin, AUR2-25075C18A-CSI, IonOpticks). Peptides were introduced into the mass spectrometer using a gradient of 3–27% buffer B (0.1% (v/v) formic acid in 80% acetonitrile) for 52 min followed by an increase to 40% buffer B over 8 min at a flow rate of 100 nL/min, and were ionized by captive spray ionization. Samples were analyzed by data-independent acquisition (DIA) using dia-PASEF with a duty cycle of 0.95 s comprising one MS1 survey scan for every 8 dia-PASEF scans, in a previously published high-speed window scheme [90]. The overall mass range was set to 400–1000 m/z. The trapped ion mobility ramp and accumulation times were set to 100 ms with a 100% duty cycle and a mobility range of 1*/K*_0_ = 0.7 to 1.3 V·s/cm^2^. Fragmentation was performed by collision-induced dissociation, with the collision energy linearly interpolated between 20 eV at 1*/K*_0_ = 0.6 V·s/cm^2^ and 59 eV at 1*/K*_0_ = 1.6 V·s/cm^2^.

#### 4.5.7 MS data processing

Raw data for N/O/C and OrgIP were both searched with FragPipe version 22. The N/O/C data were searched in DDA mode and the OrgIP data in DIA mode. Both were searched against canonical and isoform proteins from the reviewed (Swiss-Prot) and unreviewed (TrEMBL) human protein FASTA database (downloaded from UniProt on 8 January 2024) with reverse decoys and contaminants added by the built-in Frag-Pipe module, with the enzyme set to trypsin. Missed cleavages were set to a maximum of 2 and peptide length to between 7 and 50 amino acids. Cysteine carbamidomethylation was set as a fixed modification, and oxidation of methionine, acetylation of the protein N-terminus, and phosphorylation of serine, threonine, and tyrosine were added as variable modifications. Specifically, the N/O/C DDA raw data were searched with the default settings of the LFQ-phospho workflow. A target-decoy search strategy was applied using MSFragger version 4.1 [73]. Peptide-spectrum matches were rescored using MSBooster version 1.2.31 and Percolator version 3.6.5. Proteins were inferred and a 1% false discovery rate (FDR) applied using ProteinProphet in Philosopher version 5.1.1. Quantification was performed using IonQuant version 1.10.27 with match-between-runs and MaxLFQ activated. The OrgIP DIA raw data were searched with the default settings of the `DIA_SpecLib_Quant_Phospho_diaPASEF workflow`. First, pseudo-MS/MS spectra were generated using diaTracer version 1.1.5. Second, the pseudo-MS/MS spectra were searched against the FASTA database in a reversedecoy fashion using MSFragger version 4.1. Peptide-spectrum matches were rescored with MSBooster version 1.2.31 and Percolator version 3.6.5. Protein inference and a 1% FDR filter were performed using ProteinProphet and Philosopher version 5.1.1. A data-derived empirical library was generated using EasyPQP version 0.1.49, and quantification was performed on the original DIA data with the built-in DIA-NN version 1.8 module [74] with cross-run normalization activated.

### 4.6 Annotation and comparative analysis of the iNeuron map

#### 4.6.1 Data processing and map annotation

DIA-NN protein-group MaxLFQ intensities were imported into *grassp*, treating unquantified entries as missing. Sample protein counts, total intensity, dropout, and principal components were inspected. No samples were removed. Proteins required detection in at least three replicates of at least five of 33 groups (24 bait–experiment combinations and nine controls) and mean within-group coefficient of variation (CV) below 0.35, leaving 9196 proteins. Intensities were transformed as log(1 +*x*), and missing values were imputed per sample from a Gaussian distribution centered 2.2 s.d. below its observed log-intensity mean, with width 0.3 s.d. Within each experiment, enrichment was the difference between bait and untagged-control median log intensities. Two-sided, equal-variance two-sample *t*-test *P* values were retained without significance filtering.

Separately measured nuclear, organellar, and cytosolic fractions were imported from FragPipe. Proteins required detection in all three replicates of at least one fraction, protein probability *>* 0.95, at least two peptides, and mean CV *<* 0.3, retaining 7627 proteins. After the same transformation and imputation, fraction replicates were summarized by their median and each protein’s three-fraction profile was mean-centered. Combining 24 OrgIP contrasts with these fractions, standardizing each feature, and harmonizing identifiers yielded 5973 proteins with 27 features. Euclidean graphs used 20 nearest neighbors and random seed 0, and UMAP used minimum distance 0.2 and seed 42.

The iNeuron map was annotated with *grassp* using independent diffusion of UniProt subcellular-location annotations (Section 4.4.2). The protein-neighborhood graph was constructed from standardized OrgIP and fractionation profiles using 20 nearest neighbors, Euclidean distance, the UMAP connectivity kernel, and random seed 0. For the neuronal-localization comparison, we used the independent diffusion annotation of the portal map in the consolidated UniProt vocabulary (`ind_diff_uniprot_coarse`). We retained each protein’s existing compartment assignment from *grassp*. Where no assignment was available, we used the class with the highest probability. These assignments are recorded in Supplementary Table S3.

#### 4.6.2 Comparison of organelle enrichment

For Fig. 3a and Supplementary Fig. S4, both maps were taken from the portal (`Frank_2026_iNeuron` and `Hein_2025_HEK293_WT`). Of the 3750 proteins quantified under the same UniProt accession in both maps, 981 were part of the curated marker panel of Hein et al. [3] and served as reference proteins.

The mitochondrion enrichment comparison used one TOMM70A contrast in iNeurons and four TOMM20 contrasts in HEK293T cells, both baits being outer-membrane import receptors. The plasma membrane used one ATP1B3 contrast per map, and the ER used the SEC61B and NCLN contrasts, four in iNeurons and three in HEK293T cells. Each contrast is a replicate-aggregated bait–experiment enrichment, and the contrasts of a compartment were averaged per protein on the log_2_ scale. The same procedure was applied to the six further compartments with baits in both maps (Golgi apparatus, early endosome, lysosome, peroxisome, stress granule, and ribosome) for Supplementary Fig. S4a. For each compartment, the median enrichment in each map, the number of markers with lower enrichment in iNeurons, and the Spearman correlation between the maps over markers were reported.

Held-out resolution was scored by competitive diffusion as described in Section 4.8.3, on each map’s own portal graph. The 981 shared markers were split into five stratified folds, repeated five times (25 folds) with identical folds for both maps, and compartments with fewer than eight shared markers were not scored. Intervals are the mean ±1.96 standard errors over folds. To assess the contribution of differences in bait coverage between the two maps, we repeated the resolution comparison using 17 aligned features: 14 bait dimensions and the nuclear, organellar, and cytosolic fractions (Supplementary Fig. S4c). TOMM20 in HEK293T was paired with TOMM70A in iNeurons, and the other bait identities were shared. Within each map, enrichment contrasts for the same bait were averaged per protein, and each feature was standardized across proteins to zero mean and unit standard deviation. Each map retained its full protein set, and a new graph was constructed using Euclidean distance, 20 nearest neighbors, and random seed 0. Competitive diffusion and AUPRC scoring used the same 981 shared markers and the same 25 folds as the portal-graph comparison. Differences were calculated as iNeuron minus HEK293T within each fold, with confidence intervals computed as above.

All counts concern proteins, not biological replicates, and differences in bait, sample preparation, and acquisition limit attribution of the enrichment differences to cell type alone.

#### 4.6.3 Database-derived neuronal localization reference and comparison

We screened human proteins with experimentally supported Gene Ontology cellular-component annotations to neuronal structures. Of 377 proteins with at least one supporting publication, 190 lacked an annotation mapping to an eligible subcellular-compartment class and a further 88 were not detected in the iNeuron map, leaving 99 proteins. Requiring at least two distinct supporting publications per protein excluded 82 proteins and yielded a reference set of 17 proteins supported by 40 distinct publications (Supplementary Table S3). Annotations were retrieved from QuickGO (GO release 2026-09-27; retrieved 28 October 2026) for the terms synapse (GO:0045202), neuron projection (GO:0043005), neuronal cell body (GO:0043025), axon initial segment (GO:0043194), and node of Ranvier (GO:0033268), including all descendants connected through `is_a` and `part_of` relationships. Queries were restricted to protein gene products from *Homo sapiens* and experimental evidence represented by ECO:0000269 and its descendants. NOT-qualified annotations were discarded.

Expected compartment labels were initially derived from each protein’s experimental human GO cellular-component annotations supported by the same neuronal publications. Terms were mapped to compartment classes through `is_a` and `part_of` relations to fixed GO anchors, retaining the most specific matches. Synaptic vesicles mapped to plasma membrane and endosome; postsynaptic density, postsynaptic specialization, and presynaptic active zone to cytoskeleton and cell cortex, except for membrane terms; and dense-core vesicles to Golgi apparatus. For C9orf72, the expected-label set was supplemented with lysosome and an endosome proxy following review of the cited neuronal evidence. Expected ribonucleoprotein-granule labels admitted cytosolic ribosome classes as biologically acceptable alternatives, and generic cytoplasm annotations from at least two neuronal publications supported cytoplasm as an acceptable alternative. Expected-label sets remained fixed when comparing annotations, including classes absent from the annotation vocabulary (e.g., cell cortex). Annotations from other publications, records without a publication, and UniProt locations were excluded from the reference.

Each protein’s assignment was compared with its fixed expected-label set and acceptable alternatives. A match was recorded for an expected label, a documented subtype, or an acceptable alternative. Stress granule was recognized as a subtype of RNA granules. A coarse match was recorded for Cytoplasm against a neuronal or cytoplasmic RNA-granule expectation, or against Cytoskeleton or Cell cortex. These broader matches indicate compatibility without establishing localization to the specific structure. A Lysosome assignment against a combined Cell membrane and Endosome expectation was a nonmatch unless Lysosome was separately expected or acceptable. Exact agreement took precedence over coarse compatibility, and all remaining comparisons were classified as no match. Match categories were determined from label identity without a probability cutoff.

Each gene was represented by one protein group. Primary gene-symbol tokens were matched exactly, ignoring case, and multiple matches were resolved by the highest stored `total_intensity`. Unresolved genes were recorded as not detected. We note that human neuronal annotations were sparse, biased toward disease-associated proteins, receptor subunits, and sensory-tissue proteins, and did not distinguish cell lines from stem-cell-derived neurons or nervous tissue. Restricting labels to neuronal publications left most expected classes supported by a single publication.

### 4.7 Creation of a differential centrifugation HEK293T map

#### 4.7.1 Cell culture and differential centrifugation

Each replicate of HEK293T cells was grown in a 15 cm cell culture dish at 37 °C and 5% CO_2_ in DMEM (high glucose, Gibco 11965-092, Thermo Fisher Scientific) with 10% fetal bovine serum (Omega Scientific, FB-11). The differential centrifugation procedure for HEK293T cells was performed as previously described [11] with the following modifications. Samples were generated in a label-free experimental setup. Ultracentrifugation was performed with a 70 Ti fixed-angle rotor (Beckman Coulter). Each post-centrifugation protein lysate was subjected to probe sonication (Hielscher UP200St) with amplitude set to 40% and cycle time set to 50% for 1 min. Acetone precipitation was performed on 100 µg of protein lysate by adding 5 volumes of ice-cold acetone followed by incubation at −20 °C for 1 h and protein pellet centrifugation at 4 °C for 5 min at 10,000 × *g*. The recovered pellet was resuspended in 50 mM Tris pH 8, 8 M urea, and the following steps were performed on a Thermomixer set to 1000 rpm: reduction with 1 mM DTT (Pierce™ DTT NoWeigh™, Thermo Fisher Scientific, A39255) for 20 min at RT, alkylation with 5 mM iodoacetamide (Pierce™ Iodoacetamide Single-Use, Thermo Fisher Scientific, A39271) for 20 min at RT in the dark, and digestion with 2 mAU of LysC (Lysyl Endopeptidase, FUJIFILM Biosciences, Fisher Scientific, NC9242798) overnight at RT. The samples were then diluted 1 in 4 with 50 mM Tris pH 8 and further digested with 1 µg of trypsin (Pierce™ MS Grade Trypsin Protease, Thermo Scientific, PI90058) for 4 h at RT. Samples were then acidified to a final concentration of 1% TFA. Acidified peptide (20 µg per fraction) was desalted and triple-fractionated on SDB-RPS StageTips as described for the iNeuron N/O/C samples (Section 4.5.5).

#### 4.7.2 Liquid chromatography and MS acquisition

Dried peptides were resuspended in 10 µL of 2% acetonitrile/0.1% TFA. A 2 µL aliquot of each peptide fraction was analyzed on a Fusion Lumos mass spectrometer (Thermo Fisher Scientific, San Jose, CA) coupled to an EASY-nLC 1200 liquid chromatography system (Thermo Fisher Scientific, San Jose, CA). Peptides were separated by capillary reverse-phase chromatography on a 50 cm column with 75 µm inner diameter packed with 1.8 µm C18 particles (Dr Maisch GmbH, Germany) using a two-step linear gradient of 3–27% buffer B (0.1% (v/v) formic acid in 80% acetonitrile) for 105 min followed by 27–40% buffer B for 15 min at a flow rate of 300 nL/min. Data were acquired in DDA mode with a Top N = 20. Full MS scans were acquired in the Orbitrap mass analyzer with a resolution of 120,000 (FWHM) and an m/z scan range of 375–1500. Selected precursor ions were subjected to fragmentation using HCD with a quadrupole isolation window of 0.7 m/z and a normalized collision energy of 31%. HCD fragments were analyzed in the Orbitrap mass analyzer with a resolution of 15,000 (FWHM). Fragmented ions were dynamically excluded from further selection for a period of 60 s. The AGC target was set to 400,000 and 50,000 for full MS1 and MS2 scans, respectively. The maximum injection time was set to Auto in both cases.

#### 4.7.3 MS data processing

Raw data were searched against a Swiss-Prot reviewed-only human protein FASTA database (downloaded from UniProt on 20 April 2020) using MaxQuant version 1.6.6.0 [72] with default settings. Oxidation of methionine and acetylation of the protein Nterminus were added as variable modifications. Match between runs and the MaxLFQ normalization strategy were activated.

### 4.8 Integration of DC and OrgIP

To test whether a small panel of OrgIP pulldowns can complement a DC map (Fig. 3d– g), the HEK293T DC map described above was combined with the HEK293T OrgIP map of Hein et al. [3]. Both maps were processed from their protein intensities with *grassp*, so that every feature of the combined map was produced by the functions a user would apply.

#### 4.8.1 Enrichment profiles

The DC dataset comprised six replicates of seven fractions. Reverse hits, potential contaminants, and proteins identified only by site were removed, and proteins quantified in at least four replicates of at least two fractions were retained, leaving 8595 proteins in the final map. Intensities were transformed as log(1+*x*), and missing values were imputed per sample by drawing from a Gaussian distribution centered 1.8 s.d. below the mean of the observed log intensities of that sample, with a width of 0.3 s.d. (`grassp.pp.impute_gaussian`). Each fraction was expressed as the log fold change of a protein’s intensity over the median of the other fractions of the same replicate (`grassp.pp.calculate_enrichment_vs_all`), and the six replicates were summarized by their median, giving one seven-dimensional enrichment profile per protein. Each fraction was then standardized to zero mean and unit s.d. across proteins. The resulting object can be accessed on the *grassp* portal (`Frank_2026_HEK293`).

The OrgIP profiles were recomputed from the MaxQuant protein groups of Hein et al. [3] (14,551 protein groups, 84 pulldown and control samples in triplicate). Contaminants were removed as for the DC map, and the three crude nuclear, organellar, and cytosolic fractions were set aside. Proteins detected in at least three of the remaining samples were retained, intensities were log_2_-transformed and imputed per sample as above, and replicates were summarized by their median. Each pulldown was then expressed as the log_2_ fold change of a protein’s intensity over the median of all other samples of the same experimental batch, i.e., the other pulldowns and the untagged controls (`grassp.pp.calculate_enrichment_vs_all`), using the five batches defined by Hein et al. Comparing within batches rather than within experiments was necessary because two experiments carried no untagged control. The untagged controls were then discarded. The candidates for selection were the 37 pulldowns that Hein et al. selected to represent their baits, which excludes repeat experiments of a bait.

#### 4.8.2 Combined map

Both maps were collapsed to gene level by averaging the profiles of accessions sharing a curated gene symbol and were joined on the 7667 genes measured in both. Marker labels from the literature-curated panel of Hein et al. [3] were pooled over the two maps and assigned by gene symbol, labeling 1740 of these genes with one of 16 compartments. The combined feature matrix, the seven DC fractions plus the selected pulldowns, was standardized to zero mean and unit s.d. per feature, and a protein-neighborhood graph (*k* = 20, Euclidean distance) was built on it with `scanpy.pp.neighbors` as for the curated maps (Section 4.9.3). The UMAP of Fig. 3g was computed from this graph with a minimum distance of 0.1.

#### 4.8.3 Resolution on held-out markers

Resolution was scored as the recovery of held-out compartment markers by competitive diffusion. Markers were split into five stratified folds. The markers of four folds seeded label spreading over the map’s graph (Eq. 8, *α* = 0.5, with class balancing), and each compartment was scored on the held-out fold by the AUPRC of its diffused probability against membership of that compartment. Scores were averaged over folds and, for macro-AUPRC, over compartments. The split was repeated five times (25 folds) and intervals were computed as the mean ±1.96 standard errors over folds. For Fig. 3d, each map was scored at protein level on its own graph as distributed on the portal, with the markers matched to that map by UniProt accession (1613 proteins in the DC map, 1921 in the OrgIP map).

#### 4.8.4 Pulldown selection

Pulldowns were added to the DC map by forward greedy selection, starting from the DC-only map and adding the most informative pulldown. In each round, every remaining candidate was scored as an addition to the current panel, rebuilding the combined map with the candidate’s column appended and recomputing its graph. After all remaining candidates had been scored, the candidate with the highest held-out macroAUPRC was added, until all 37 had been added (703 total candidate evaluations). Candidates within a round were scored independently, so their evaluation order did not affect the selection. Because a pulldown’s contribution depends on the panel it joins, each candidate required rebuilding the combined map, and the selection could not be computed from scores of individual pulldowns. The selection used a single 5fold split. The resulting nested panels were then re-scored over 25 folds for Fig. 3f,g. As a baseline, the nested panels of 100 random orders of the 37 pulldowns were scored in the same way, and their mean and s.d. across orders are shown at each panel size. The full OrgIP map shown as a reference in Fig. 3f comprised all 68 recomputed pulldowns and the three crude fractions, each crude fraction expressed against the other two, without the DC fractions. It was built and scored on the same genes, markers, and folds. The combined map of Fig. 3g comprises the DC map and the first nine pulldowns of the greedy order.

### 4.9 Curation and analysis of published datasets

To create a comprehensive collection of processed public datasets, we curated MS-SP datasets from the literature with the following criteria:

- The dataset must be an MS-SP dataset described in a peer-reviewed or preprint publication (as of March 2026).
- Quantitative data must be available as supplementary information of the publication or in a public repository.
- The assayed organism must be one of the following: human (*Homo sapiens*), mouse (*Mus musculus*), or yeast (*Saccharomyces cerevisiae*).

The resulting collection comprised 103 maps: 101 maps from 30 publications appearing between 2010 and 2025, together with the two maps generated in this study. Of these, 88 were human, 11 mouse, and 4 yeast, and were classified into 14 subcellular enrichment strategies (Dynamic Organellar Maps, LOPIT-DC, hyperLOPIT, dLOPIT, DIA-LOP, DC, density-gradient centrifugation, differential detergent separation, protein correlation profiling, SubCellBarCode, OrgIP, BioID, COLA, and iCOLA) and five quantification strategies (data-dependent acquisition, data-independent acquisition, TMT, SILAC, and iTRAQ). The composition of the collection and the curated metadata of every map are given in Supplementary Table S6.

Each dataset was then manually downloaded and processed using *grassp*, including filtering and quality control, graph construction, dimensionality reduction, semi-supervised annotation, and visualization. Finally, ontology-restricted metadata about experimental technique, species, cell type, and publication were added, yielding a processed AnnData object. Each analysis was documented in a Jupyter notebook and stored in the curation repository (see Data and Code Availability). The guidelines for processing and curation are described in the following sections.

Curation was organized so that every map could be reproduced from its published source files. Each map was represented by one row of a manifest table, one directory, one download script that fetched the exact supplementary or repository files used, and one notebook that converted those files into AnnData. Because the source publications differed in what they deposited, the notebooks necessarily differed in their parsing and in which quality-control steps the source study justified. Everything downstream of the quantitative matrix followed the common rules described below, and per-map deviations are documented in the notebooks.

#### 4.9.1 Protein identifier harmonization

Comparing maps across studies requires a single identifier space, so all maps were keyed on UniProt [37] accessions. Where a study reported protein groups, the leading accession of the group was used. Where it reported only gene symbols, symbols were mapped to accessions offline against the same UniProtKB reference release used for validation, resolving one-to-many matches by preferring a primary gene-name match over a synonym match, then a reviewed (Swiss-Prot) over an unreviewed entry, then the higher UniProt annotation score, then the lexicographically smallest accession. Accessions that had been merged or demerged since publication were updated through the UniProt identifier-mapping service. Observations whose accession could not be resolved to a current entry, and duplicate accessions arising from identifier updates, were removed, because an unresolvable identifier cannot be compared across maps. This affected 1925 of 481,925 protein observations across the collection (0.40%, median 0.05% per map, maximum 8.9%), leaving 480,000 protein observations on a common accession space.

#### 4.9.2 Preprocessing and quality control

Quantitative matrices were read with *protdata* (see Section 4.1) where the deposited file was a search-engine output, and parsed directly otherwise. Reported quantities were placed in .X, secondary quantities that the study deposited alongside them (e.g., spectral or ratio counts) and a log transform of the reported intensities were kept as separate layers, and sample names were parsed into explicit .var columns for replicate, fraction or bait, and fraction order, so that the sample structure was machine-readable rather than encoded in column names.

Quality-control metrics were computed with `grassp.pp.calculate_qc_metrics`, yielding per-protein detection counts, mean and total intensity, and dropout fraction, and per-sample protein counts, total intensity, and dropout fraction. Filtering followed the source publication rather than a uniform threshold, since the studies differed in design and in how much filtering they had already applied. Imputation was applied only where the source study imputed, for 20 maps, by sampling from a downshifted Gaussian on log-transformed intensities.

Fractionation and pulldown designs were converted to subcellular enrichment profiles with `grassp.pp.calculate_enrichment_vs_all`. The function first summarized the replicate samples of each fraction by their median and then scored each fraction against all remaining fractions of the map. Where the original publication suggested it, enrichment was expressed as the proportion of a protein’s summed intensity across fractions, and proteins without a defined proportion were removed. The two maps generated in this study were instead expressed as natural-log fold changes, over the other fractions of the same replicate for the DC map and over the untagged controls of the same experiment for the iNeuron OrgIP map (Sections 4.8 and 4.6.1). The remaining 23 maps deposited ratio-type or already-normalized profiles, for which no enrichment was computed. Their replicates, where present, were summarized per fraction. For 34 maps, each fraction was additionally standardized to zero mean and unit variance across proteins before dimensionality reduction and graph construction. The input intensities were retained in a layer where applicable, and the background samples used for each comparison were recorded in .var.

#### 4.9.3 Graph construction

A protein-neighborhood graph was built for every map from the enrichment profiles with `scanpy.pp.neighbors` [77], using the UMAP connectivity kernel [32] and a standard neighborhood size of *k* = 20 for all 103 maps. Neighbors were computed on the profiles directly rather than on a principal-component representation, as is typically done in single-cell workflows. This graph was the substrate for every downstream step: embeddings, both annotation strategies, and the cross-study integration described below. Because one graph was built per map in the same way throughout, annotations and integration remained comparable across maps whose feature axes were not.

#### 4.9.4 Dimensionality reduction

Principal-component analysis was computed on the enrichment profiles, retaining up to 50 components and fewer where a map had fewer features. Two UMAP embeddings were then computed from the graph with a minimum distance of 0.2: a two-dimensional embedding and a three-dimensional embedding, both stored on the object, so that the portal’s flat and rotatable map views and the annotations shown on them were derived from the same graph.

#### 4.9.5 Compartment gene set generation for independent diffusion annotation

Independent diffusion annotates a map against a vocabulary of compartment gene sets rather than against a marker panel, so the vocabulary determines the granularity of the resulting labels. Four vocabularies of increasing granularity were built and are distributed with *grassp*:

1. a consolidated UniProt subcellular-location vocabulary (24 terms for human and mouse, 23 for yeast);
2. the full UniProt subcellular-location hierarchy (240 human, 241 mouse, and 107 yeast terms);
3. the knowledge channel of the COMPARTMENTS database [80] (874 terms, human-only);
4. a cleaned GO CC vocabulary [36] (1829 terms, human-only).

All four sources change between releases, so each vocabulary was built once, versioned, and distributed with *grassp* rather than fetched at annotation time. The UniProt sets were retrieved from the UniProt REST API [37] at release 2026 03 (2 September 2026). GO CC was built from the Gene Ontology release of 19 June 2026 (ontology version 2026-06-15). COMPARTMENTS publishes a single rolling file with no versioned archive and was obtained on 24 September 2026.

Both UniProt vocabularies were assembled by querying reviewed (Swiss-Prot) entries of each species for every subcellular-location term. These queries match down the location hierarchy, so a term carries the proteins annotated to it together with those annotated only to its descendants. Terms therefore contain their children rather than partitioning them. The full hierarchy keeps every term as its own gene set, after removing host-cell and viral locations, structures absent from the species, and container terms broad enough to absorb most of the proteome, such as “Membrane” and “Endomembrane system”. Terms with no reviewed protein in a species were also dropped, which removes organelles that species does not have, such as plastids and thylakoids in the mammals. The vacuole and the spindle pole body are genuine yeast structures and were kept for yeast, but were excluded by name for human and mouse. The consolidated vocabulary instead maps 52 fine-grained location terms onto 16 compartments that a fractionation or pulldown experiment can usually separate into distinct profiles. Five finer terms (ERGIC, early endosome, NPC, P-body, and stress granule) were kept as terms of their own in addition to the compartment that contains them. The two cytosolic ribosomal subunits and the proteasome were added from EBI Complex Portal cross-references (for the mouse proteasome, from the UniProt keyword “Proteasome”), because these complexes routinely resolve as their own clusters in enrichment profiles even though they are not subcellular locations. The yeast vocabulary additionally contains the vacuole and lacks the cilium and the lysosome.

GO CC terms were filtered because they were partially incompatible with detectable compartments: some terms exist only to organize the ontology, and its high-throughput evidence codes place most of the proteome in “extracellular exosome”. Annotations were therefore restricted to the `located_in`, `part_of`, and `is_active_in qualifiers`, high-throughput and no-data evidence codes were discarded, and the remainder were propagated over `is_a` and `part_of`. Terms that GO itself flags as not for direct annotation were removed. Additionally, seven structural root terms were removed for being too unspecific: membrane-enclosed lumen, organelle, protein-containing complex, membrane, `cellular_component`, cellular anatomical entity, and cytoplasm. Grouping terms that lie under none of 12 major compartments (nucleus, mitochondrion, ER, Golgi apparatus, endosome, lysosome, peroxisome, plasma membrane, cytosol, extracellular region, vacuole, and cytoskeleton) but whose descendants span three or more of them were removed as well, because they typically describe a type of structure rather than a distinct subcellular localization. Near-duplicate gene sets were finally collapsed at a Jaccard index of 0.9, keeping the most specific term of each cluster, since at this granularity synonymous terms otherwise split the evidence for one compartment between several labels.

The COMPARTMENTS vocabulary was downloaded from the manually curated knowledge channel, keeping assignments of at least three of five confidence stars and terms with at least five human genes. Generic and non-locational terms were removed by an explicit list of 43 manually curated terms and by 15 name patterns (e.g., “side of” and “component of membrane”), and terms with identical gene sets were merged. Near-duplicate gene sets were collapsed at a Jaccard index of 0.8, keeping the shortest and most canonical organelle name of each cluster.

#### 4.9.6 Compartment annotation of the curated maps

Every map carries three kinds of localization evidence, kept as separate columns rather than collapsed into one call: the labels of the source study, competitive diffusion of curated marker sets, and independent diffusion of compartment vocabularies. Both diffusion annotations were computed with *grassp* 0.8.0, over the protein-neighborhood graph described in Section 4.9.3.

The labels reported by the source study were stored as `author_annotation`, and where a study designated a subset of proteins as compartment markers, those were kept as a separate column (`author_markers`). The two maps generated in this study have `author_annotation` set to the annotations of `ind_diff_uniprot_coarse` (see below).

All marker sets distributed with *grassp* for the species of a map were matched onto each dataset by UniProt accession with `grassp.pp.add_markers`: eight for human maps, two for mouse maps, and one for yeast maps. The human markers consist of five sets distributed with pRoloc [15], the marker list and the full annotation of Hein et al. [3], and a set derived from the integrated pan-human map (Section 4.10). The last set holds the proteins that a separate independent diffusion run on the graph of the integrated map (*grassp* 0.7.0, current consolidated UniProt vocabulary) assigned to a compartment with a probability of at least 0.7 while no other compartment exceeded 0.3, ignoring compartments that are routinely co-annotated with the assigned one, for the 17 compartments with at least 10 such proteins. Each set seeded one competitive diffusion annotation (`grassp.tl.competitive_diffusion`, Section 4.4.1), named `comp_diff_followed` by the name of the marker set. Competitive diffusion was run with default parameters, except that markers were not clamped to their seed labels. Compartments with fewer than five markers in a map were left out of the seed, and proteins whose highest compartment probability was below 0.5 received no label.

Independent diffusion (`grassp.tl.independent_diffusion`, Section 4.4.2) was run with default parameters on all datasets. Every map was annotated with the consolidated and the full UniProt vocabulary of its species (`ind_diff_uniprot_coarse and ind_diff_uniprot_fine`), and the 88 human maps additionally with GO CC (`ind_diff_gocc`) and COMPARTMENTS (`ind_diff_compartments`). Terms with identical gene sets were merged on loading, which left 235 human, 236 mouse, and 99 yeast terms of the full UniProt hierarchy. For this vocabulary, labels were restricted to terms with at least 20 members in the map. The GO CC and COMPARTMENTS annotations of the portal maps were computed with earlier builds of the two vocabularies (1824 and 790 terms, built in August 2026), as documented in the curation repository (https://doi.org/10.5281/zenodo.24484245).

#### 4.9.7 Consistency of compartment labels across maps

To compare how consistently a protein is labeled across studies (Supplementary Fig. S5), four annotations were read from the portal objects of the 51 unperturbed human maps that entered the integrated pan-human map (Section 4.10), for the proteins of that map. The author annotation was used either as published, with every distinct label string counted as a label of its own, or after harmonization into the consolidated UniProt vocabulary with the dictionaries of Section 4.4.4. A compound author label (e.g., “ER, Peroxisome”) was kept as the set of compartments it names, and author labels without a consolidated counterpart were kept as labels of their own, so that both readings of the author annotation label the same proteins. The two diffusion annotations were competitive diffusion of the marker list of Hein et al. [3] (`comp_diff_hein2025_gt`) and independent diffusion of the consolidated UniProt vocabulary (`ind_diff_uniprot_coarse`), as stored in the portal objects (Section 4.9.6).

Because proteins are detected in different numbers of maps, the number of distinct compartments assigned to a protein was counted over a fixed number of maps. For each annotation and protein, 10 of the maps in which the protein is labeled were drawn without replacement, and the probability of obtaining 1, 2, . . . , 10 distinct labels was computed exactly over all possible draws rather than by sampling. As a second measure, the fraction of pairs of a protein’s labeled maps that assign different labels was computed over all of its labeled maps. The four annotations were compared on the 5665 proteins that each of them labels in at least 10 maps.

#### 4.9.8 Metadata curation

Each map was annotated by hand from its source publication with a fixed set of fields, each restricted to a controlled vocabulary or ontology so that maps could be selected and compared programmatically: organism (NCBITaxon [91]), tissue of origin (UBERON [92]), cell type (Cell Ontology [93]), cell line (Cellosaurus [94]), disease state (MONDO [95], or the PATO [96] term for normal), sex (PATO), development stage (HsapDv or MmusDv [97]), and sample class (cell line, tissue, primary cell culture, or organoid). Tissue of origin, cell type, and cell line were curated independently of one another, so that a cell-line map could carry the tissue its line was derived from without asserting a cell type. Two further fields described the experiment: the enrichment strategy and the quantification strategy, each drawn from a controlled list. A free-text field recorded the perturbation, and a Boolean field recorded whether the map was an unperturbed reference for its biological system or a perturbation arm. In total, 59 of the 103 maps were references, and the 51 human reference maps were exactly the subset used for the pan-human integration. Curated terms were checked against the publication, Cellosaurus, and the EBI Ontology Lookup Service [98].

Only the most specific tissue term was curated by hand. The broader levels served by the portal — general tissue, organ, and body system — were derived from it automatically as its first matching ancestor in the curated high-level term lists of CZ CELLxGENE Discover [67], so that the broad levels could never disagree with the specific one. Publication metadata (title, author list, journal, and date) were retrieved from CrossRef by DOI and stored on the object.

Every object was then checked with an adaptation of the CZ CELLxGENE Discover single-cell curation schema [67], extended with the proteomics-specific fields above. Validation confirmed that each ontology term existed, was not deprecated, and belonged to the branch its field required. On passing, the human-readable label of every ontology identifier was written alongside the identifier itself, so that the served objects carried both the machine-comparable term and the label a reader sees. Maps that failed validation were corrected and revalidated rather than served. All 103 maps in the collection passed.

#### 4.9.9 Curated AnnData object schema

Each curated map was exported as one AnnData object, in which proteins were observations and fractions, baits, or channels were variables (see Section 4.2). Two objects were produced per map: a pre-enrichment snapshot, which held the quantities that entered enrichment together with the formatted sample metadata, the author labels, and the marker sets, and the enriched object served by the portal, which additionally passed identifier harmonization, metadata curation, validation, and diffusion annotation. The enriched object had the following layout.

- .`X` held the subcellular enrichment profile, and .layers held, where applicable, the input intensities, their log transform, and the enrichment *P* values.
- .`obs` was indexed by UniProt accession and held protein identity (protein name, gene symbol), the author labels, the published marker sets, the per-protein quality-control metrics, and the competitive and independent diffusion annotation columns.
- .`var` was indexed by sample and held, where applicable, the sample structure (replicate, fraction or bait, fraction order, the background samples used for enrichment) and the per-sample quality-control metrics, as well as the curated metadata with both the ontology identifier and its resolved label.
- .`obsm` held the principal-component and two- and three-dimensional UMAP embeddings and the per-compartment probability matrices of the competitive diffusion annotations, and .obsp held the protein-neighborhood graph.
- .`uns` held the map title, the publication metadata, the schema version and reference, the parameters of every annotation, the compartment order of each competitive diffusion probability matrix, the fitted diffusion depth of every term of each independent diffusion vocabulary, the compartment color palettes used by the portal, and optional free-text notes that *grasspViz* displays above a specific embedding to explain a visible artifact.

Because quantitation, sample metadata, graph, embeddings, and every annotation layer live in one container, each curated map is distributed as a single h5ad file that can be downloaded from the portal and read back into *grassp*, scanpy, or — through *grasspio* — pRoloc without any further assembly.

### 4.10 Creation of the integrated pan-human map

#### 4.10.1 Dataset selection

The integrated pan-human map combines the protein-neighborhood graphs of every unperturbed human map in the collection. Of the 88 human maps, we kept the 51 that were curated as the reference state of their biological system and excluded experimental perturbations and disease states (drug treatment, genetic knockout, viral infection, growth-factor or inflammatory stimulation, stress, irradiation, dietary intervention, engineered protein relocalization, and disease tissue), so that the integrated map describes resting-state organization rather than a mixture of resting and perturbed states. The 51 maps come from 26 publications and cover 26 cell lines in addition to primary adipose progenitor cells [99], post-mortem brain tissue (angular gyrus) [100], and humanized mouse liver [25]. After the protein filtering described below, the datasets contained between 946 and 8825 proteins (median 4368). The 51 datasets used in this integration are the human maps flagged in the `serves_as_reference` column of Supplementary Table S6.

#### 4.10.2 Network construction and protein filtering

Datasets acquired with different enrichment strategies do not share a feature axis and have different sets of measured proteins. However, each protein-neighborhood graph provides information about local neighborhood structures that can be leveraged during integration. These graphs are the objects that are integrated. For every selected map *d* we took the symmetric affinity matrix **W**^(^*^d^*^)^ computed during curation (Section 4.9.3) and converted it to a list of undirected edges weighted by their affinity, without selfloops and without thresholding.

Nodes were then filtered jointly across all 51 networks, and an edge was kept only if both of its endpoints survived. We removed proteins with isoform identifiers or without a valid UniProt accession, keeping only reviewed (Swiss-Prot) human entries [37], retrieved from the UniProt REST API (20,431 entries). To prevent individual studies from dominating the signal for proteins with sparse observations, we kept only proteins quantified in at least 6 of the 51 maps. After filtering, the 51 networks contained *n* = 9530 proteins connected by 2,767,555 edges across datasets.

#### 4.10.3 Network integration with a graph convolutional network

The 51 networks were integrated with BIONIC v0.2.6 [54], an unsupervised graph-attention autoencoder that learns one *h*-dimensional embedding per protein from which every input network can be reconstructed. Each network is encoded by its own graph attention network (GAT). The network-specific representations are combined by learned per-network scaling into a single shared embedding **Z** ∈ ℝ*^n^*^×^*^h^*, and a single network is reconstructed from it by an inner product, **ZZ**^⊤^. Training minimizes the reconstruction error summed over networks,

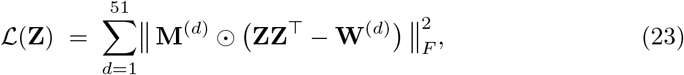

where ⊙ is the elementwise product and the mask **M**^(^*^d^*^)^ has *M_ij_*^(^*^d^*^)^ = 1 only if both proteins *i* and *j* were quantified in map *d*. BIONIC sets the diagonal of every **W**^(^*^d^*^)^ to 1 before training. The mask makes this objective usable on maps of very different depth: the loss is computed only for observed protein pairs, so the embedding is updated only where it has evidence. Note that this is an unsupervised objective. No labels or marker proteins were used to optimize the integration.

We used the architecture and optimizer settings BIONIC’s authors established for human protein–protein interaction networks: three GAT layers per network with 10 attention heads of dimension 68, a 512-dimensional embedding (*h* = 512), batches of 2048 proteins, two sampled neighbors per node, Kaiming initialization, and the Adam optimizer with a learning rate of 5.83 × 10^−4^. Two settings differed from that four-network configuration: in each epoch the 51 networks were randomly partitioned into six groups of 8 or 9, and each optimization pass encoded one group (BIONIC’s `sample_size` parameter set to 8, which by default encodes every network at every step), keeping the cost of a pass independent of the size of the corpus, and training ran for 5000 rather than 2123 epochs. The parameters of the epoch with the lowest total reconstruction loss (epoch 4992) were used to compute the final embedding. BIONIC v0.2.6 provides no option to fix the random seed and none was set, so retraining yields a numerically different embedding. The map is therefore distributed as a fixed object on the *grassp* portal. Training took 6.5 h on a single NVIDIA A100 GPU (PyTorch 1.9, PyTorch Geometric 1.7).

#### 4.10.4 Map construction

The integrated map is the 512-dimensional embedding **Z** of the 9530 proteins, stored as an AnnData object together with a protein-neighborhood graph computed on it with `scanpy.pp.neighbors` (*k* = 50, cosine distance, UMAP connectivity kernel). For the integrated map this graph plays exactly the role that **W** plays for a single map: it is the substrate for the embedding, for the annotation, and for every benchmark below.

### 4.11 Analysis of the integrated pan-human map

#### 4.11.1 Map annotation

The map was annotated by independent diffusion (Section 4.4.2) with *grassp* 0.4.0, under four label vocabularies of increasing granularity. The vocabularies were earlier builds of those described in Section 4.9.5: the consolidated UniProt subcellular-location vocabulary with 21 terms (it includes autophagosome and cell cortex and predates the ERGIC, early endosome, NPC, P-body, and stress granule terms), the full UniProt subcellular-location hierarchy (235 terms after merging identical gene sets), the COMPARTMENTS vocabulary (790 terms), and the GO CC vocabulary (1824 terms).

Each vocabulary was propagated over the map’s protein-neighborhood graph with default parameters and a fixed random seed, followed by the hierarchy-aware likelihood resolver. For the three fine-grained vocabularies, labels were restricted to terms having at least 20 members in the map. A UMAP layout (repulsion strength *γ* = 0.1, minimum distance 0.15) and Leiden [101] clusterings at eight resolutions between 0.1 and 20 were computed on the same graph, and each protein was additionally annotated with the curated marker panels, with MitoCarta3.0 [45] and MitoCoP [46] membership, and with UniProt membrane topology.

#### 4.11.2 Resolution benchmarking

##### Head-to-head design

Scoring the resolution of every map on a globally shared set of proteins would reduce the benchmark space below the number of proteins of the shallowest map, whereas scoring each map on its own proteins potentially biases the comparison in favor of maps with stricter filtering. We therefore scored every measure in a *pairwise* fashion, subsetting proteins to the intersection of each comparison pair. Write D for the maps in the analysis, *P_d_* for the proteins of map *d*, and **W**^(^*^d^*^)^ for its affinity matrix. For an unordered pair {*d, e*} we formed the shared pool

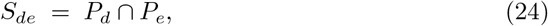

restricted both graphs to it, gave both maps the identical ground truth restricted to *S_de_*, and recorded the signed difference of their scores,

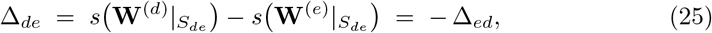

where *s*(·) is one of the two resolution scores defined below. The value reported for a map is the mean of its signed differences over the peers it could be scored against,

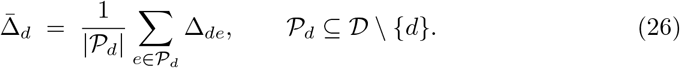

A map is thus never compared with another on proteins its dataset does not contain, and Δ̄ *_d_* reads directly as how much better, on average, a map resolves the ground truth than the maps it can be compared with. Because every comparison is paired, the ranking is also insensitive to which datasets are in the panel. The analysis comprises all 88 human maps of the *grassp* collection together with the integrated map (|*D*| = 89). No subset was selected. Both measures read a map’s protein-neighborhood graph as distributed with the portal object, and for the integrated map the graph computed on its BIONIC embedding.

##### Compartment-marker resolution

The first measure asks how well a map’s graph predicts the compartment of marker proteins it has not been told about. Ground truth was the literature-curated marker panel of Hein et al. [3], taken as the union over the corpus (2365 proteins in 16 compartments). A protein carries the same curated label wherever it appears, so the union is well defined and both members of a pair are labeled identically. On each shared pool we kept the compartments with at least five labeled proteins and scored the pair only if at least two such compartments remained.

The shared markers were split into five stratified folds, identical for both maps of the pair. In each fold, the training markers formed the seed matrix **Y**^(0)^, the labels were spread over each map’s own subgraph by label spreading (Eq. 8, `grassp.tl.competitive_diffusion`, *α* = 0.5, with class balancing, as in Section 4.8.3), and the propagated probabilities *Ŷ_ic_(d)* of the held-out markers were recorded. The out-of-fold predictions were pooled, so that every marker was predicted exactly once, from a graph that did not see its label. For each compartment *c*, the held-out probabilities *Ŷ_ic_*(*d*) were then scored against membership of *c* by the AUPRC of Eq. 3,

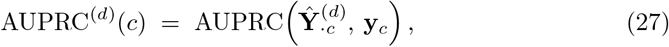

with **y***_c_* the indicator of the held-out markers of *c*. Unlike the *F*_1_ of the highest-probability label, this score does not depend on a decision threshold, and scoring by *F*_1_ instead ranks the maps the same way (Spearman *ρ* = 0.98). Because both maps of a pair receive identical folds and are scored on identical proteins, any fold-to-fold variability is shared by both sides and cancels in Eq. 25.

Differences were averaged over compartments within a pair before being averaged across peers,

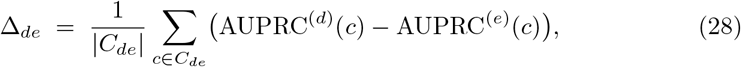

with *C_de_* the compartments scorable on *S_de_*, so that a map is not rewarded for sharing more proteins, and hence more scorable compartments, with some peers than with others. The quantity Δ̄*_d_* then follows from Eq. 26. Maps with fewer than two scorable peers were dropped. The bar strip of Fig. 4f shows Δ̄*_d_* and the heatmap below it the same average taken separately for each compartment,

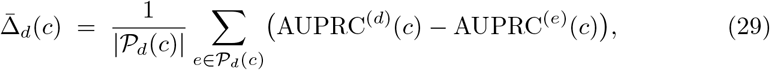

over the peers *P_d_*(*c*) for which compartment *c* was scorable. All 3916 pairs were scorable.

##### Protein-complex resolution

The second measure asks whether a map places the members of a protein complex next to one another. It is a finer question than compartment membership and uses no localization annotation at all. Ground truth was the human export of the EBI Complex Portal [66], release 2026-01-09, restricted to entries with manual physical-interaction evidence (ECO:0000353) and at least six members, with duplicate member sets collapsed. Because the curated complexes overlap, each protein was then assigned to a single complex, the last entry of the export that lists it, so that the complexes used for scoring are disjoint. This left 56 complexes covering 425 proteins. A map was scored only if it covered at least 30% of these complexes (≥ 17 of 56), where a complex counts as covered when at least four of its assigned members were quantified in the map. This held for 59 of the 89 maps. The excluded maps are the shallowest ones, and the filter tracks proteome depth rather than an identifier-mapping artifact: the fraction of accessions matching the Complex Portal lookup is nearly the same for retained and excluded maps (4.94% versus 5.04%), and complex coverage correlates with proteome size at Spearman *ρ* = 0.92. On each shared pool we kept the complexes with more than three members present (median 24 complexes over 3616 shared proteins).

Resolution was quantified as mAP in the retrieval framework of Kalinin et al. [55]. Each member *q* of a complex *K* acts in turn as a query: the remaining shared proteins are ranked by their edge weight *W_qj_*^(^*^d^*^)^ to it in that map’s graph, the co-complex members *K* \ {*q*} are the positives, and the average precision (AP) is computed over that ranking as in Eq. 3,

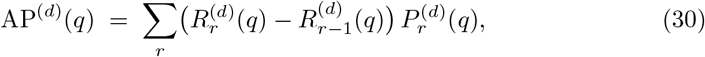

with *P_r_* and *R_r_* the precision and recall at rank *r*. Queries are averaged within a complex and complexes are then averaged,

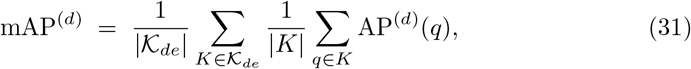

over the complexes *K_de_* retained on the shared pool, so that a large complex does not dominate the score through its many queries. Setting *s* = mAP in Eq. 25 and averaging over peers (Eq. 26) gives the mean difference plotted in Fig. 4e. All 1711 pairs of the 59 retained maps were scored.

##### Choice of the complex metric

Four alternative summaries — AP and AUROC, each pooled over all protein pairs and each macro-averaged over complexes — were computed on the same 1711 pairs. All five rank the integrated map first and have it win all 58 of its head-to-head comparisons, so the conclusion does not depend on the choice. We report mAP because it is the only well-conditioned variant. AP is sensitive to the number of positives and to the length of the ranked list: a per-query ranking orders ∼7600 candidates with 3–33 positives, whereas an all-pairs AP for a single complex rests on as few as six positives among ∼2.9 × 10^7^ pairs and is correspondingly quantized. Empirically this gives mAP about twice the signal-to-noise of the pooled AP (mean pairwise difference of the integrated map divided by its standard error, *t* = 50.5 against *t* = 25.7). The chance level of AP is the positive base rate, so AP is not comparable between complexes of different size. This does not affect the panel, which never compares complexes but plots a paired difference between two maps scored on an identical complex set, so the size-dependent term is common to both sides of Eq. 25 and cancels (measured Spearman *ρ* = −0.075 between complex size and the plotted difference).

#### 4.11.3 Comparison with simpler integration strategies

The integrated map was compared with three alternatives on its 9530 proteins (Fig. 4c and Supplementary Fig. S7). The single map was the input map with the highest marker macro-F1 (the wild-type map of Hein et al. [3]), represented by its own protein-neighborhood graph. The averaged graph was obtained by summing the affinity matrices of the 51 input maps, dividing each entry by the number of maps in which both proteins were quantified, keeping the 50 strongest edges of every protein, and symmetrizing by the elementwise maximum, so that its density matches that of the graph of the integrated map. For the annotation mean, no graph was built. Instead, every input map was annotated on its own graph and the resulting compartment probabilities of a protein were averaged over the maps in which it was quantified. Compartment resolution was scored on the ground-truth markers of Hein et al. [3] present in the integrated map, using the 16 compartments with at least eight markers and five stratified folds that were identical for all strategies. The training markers of each fold were propagated by competitive diffusion (label spreading, *α* = 0.5, markers not clamped), and the held-out markers were scored by the AUPRC of each compartment, averaged over compartments. Strategies were compared with the integrated map by two-sided Mann–Whitney *U* tests over the five folds. Protein-complex resolution was scored as the mAP of each complex (Section 4.11.2) on the same graphs and compared by twosided Wilcoxon signed-rank tests paired by complex. The annotation mean yields no graph and therefore has no complex score.

#### 4.11.4 Quantitative characterization of the Mito-cytosolic cluster

Leiden clustering was repeated on the stored integrated-map graph using resolution 12, random seed 0, and two iterations, with the undirected igraph implementation. The Mito-cytosolic group combined clusters 137 and 31 (53 and 57 proteins, respectively), yielding 110 proteins. These groups largely formed a single cluster at lower resolutions of 3–8. The mitochondrial island used as the comparison group comprised clusters 0, 1, 34, 59, 77, 110, and 134, containing 849 proteins. Group membership was held fixed across comparisons (Supplementary Tables S8 and S7).

Additional cytoskeleton and cytosol reference groups were defined at the cluster level. Cytoskeleton clusters required at least 50% of members to carry the label “Cytoskeleton” or “Cell cortex” in the map’s independent diffusion annotation (consolidated UniProt vocabulary) or the label “Actin Cytoskeleton” in the full protein annotation of Hein et al. [3]. Cytosol clusters required more than 50% of members labeled “Cytosol” in the same annotation of Hein et al. Both reference groups required at least ten members and fewer than 25% of members with mitochondrial evidence, defined as membership in MitoCarta3.0 or MitoCoP, the label “Mitochondrion” in the annotation of Hein et al., or the label “Mitochondrion” in the map’s independent diffusion annotation. These criteria yielded 230 cytoskeleton and 1453 cytosol proteins. Mitochondrial-matrix frequencies were calculated among MitoCarta3.0-listed members, counting only exact Matrix annotations as positive and retaining members with unknown subcompartments in the denominator. Denominators were 74, 793, 2, and 26 for the Mito-cytosolic, mitochondrial island, cytoskeleton, and cytosol groups, respectively. Transmembrane annotation was defined by the UniProt Transmembrane keyword. Mitochondrial–cytosolic annotation required both a mitochondrial term and the word cytoplasm or cytosol in the cached UniProt subcellular-location comment, after removing evidence tags and converting text to lower case. UniProt annotation frequencies used all group members as denominators, with missing records counted as negative. These text-derived flags do not distinguish isoforms or establish simultaneous localization.

OrgIP and fractionation measurements from wild-type HEK293T cells [3] were matched by UniProt accession. Per-protein mitochondrial enrichment was the median log_2_ enrichment across four TOMM20 pulldowns (09, 10, 12, and 15). The share of each protein in the cytosolic fraction of the nuclear, organellar, and cytosolic fractionation was used as supplied, as a proportion without transformation. Comparisons included 103 Mito-cytosolic, 742 mitochondrial island, 167 cytoskeleton, and 1226 cytosol proteins with measurements, summarized by group medians and interquartile ranges (Supplementary Fig. S8e). Because the Hein HEK293T wild-type dataset contributed to construction of the integrated map, these measurements characterize the experimental profiles underlying the cluster placement and do not constitute independent validation.

#### 4.11.5 Comparison with SurfaceBench

SurfaceBench ground-truth labels [61] (https://api.deliverome.org/surfaceome/v1/benchmark, accessed 19 November 2026) were matched to the integrated map by UniProt accession, which retained 80 of the 147 benchmark proteins (21 permanently accessible, 20 conditionally accessible, and 39 not accessible). The memberships for this comparison were computed with the benchmark proteins held out: the 80 proteins were removed from every gene set of the full UniProt subcellular-location vocabulary (235 terms after merging identical gene sets, Section 4.9.6), and independent diffusion (*grassp* 0.8.0, default settings) was run on the protein-neighborhood graph of the integrated map. For each protein, the top compartment was defined as the term with the highest membership probability, without applying the likelihood resolver. The plasma membrane was represented by the UniProt term “Cell membrane”. Plasma-membrane memberships of permanent and conditional proteins were compared with a two-sided Mann–Whitney *U* test. For conditional proteins, the baseline location was read manually from the free-text rationale that SurfaceBench provides for each protein and assigned to a compartment of the consolidated UniProt vocabulary or to “Secreted” (Supplementary Table S9). Agreement was counted when the top compartment, a term of the full vocabulary, fell within the baseline compartment, with the ERGIC accepted for a baseline location in either the ER or the Golgi apparatus. One protein (LY6K), whose rationale describes expression restricted to testis and cancer cells, was not scored. The 39 proteins that are not accessible are not shown in Supplementary Fig. S9. A protein that is not accessible is not necessarily absent from the plasma membrane: SurfaceBench classifies 7 of the 39 as anchored to the inner face of the plasma membrane, and the plasma membrane was the top compartment of 6 of these, as expected for a map that resolves membrane association but not the face of the membrane. SurfaceBench classifies the other 32 as intracellular or secreted proteins, and the top compartment of 30 of them was a compartment other than the plasma membrane (Supplementary Table S9).

#### 4.11.6 The integrated map as a prior for perturbation-response prediction

##### Perturbation-response prediction with PRESAGE

We used PRESAGE [65] (https://github.com/genentech/PRESAGE, commit 2c7b231) to test the integrated pan-human map as prior knowledge for predicting transcriptional responses to unseen genetic perturbations (Supplementary Fig. S10). In PRESAGE, each knowledge source is converted to a 128-dimensional gene embedding and passed through its own feed-forward network. The sources are combined by learned global weights and graph attention, and the model predicts the pseudobulk log-fold-change profile of each perturbation.

We used the K562 essential-gene CRISPR interference Perturb-seq screen [102] as distributed by scPerturb [103] (2058 targeted genes). Raw counts were normalized to 10,000 counts per cell and log-transformed (log(1 + *x*)), as described by Littman et al. [65]. We used the published PRESAGE 5-fold random perturbation splits, excluding split 0, which the authors used for hyperparameter tuning. Every model configuration was trained on each of splits 1–4, with three random seeds for the 40-source reference model and its ablations, and with one seed for models with a single knowledge source. All hyperparameters were taken from the released PRESAGE configuration and were identical across runs. Knowledge sources were loaded with the unmodified upstream PRESAGE code, so that the 40-source baseline reproduces the published model.

##### The integrated map as a knowledge source

The integrated map embedding **Z** (512 dimensions, 9530 proteins; Section 4.10) was provided to PRESAGE as an additional tabular source. Each protein’s embedding was *ℓ*_2_-normalized before input, because the embedding is designed to work with cosine distances. Perturbation targets were matched to map proteins through their Ensembl gene identifiers (UniProt ID mapping to Swiss-Prot accessions), with exact gene symbols as a fallback, and proteins sharing an identifier were averaged. This covered 1874 of the 2058 perturbation targets (91.1%). All other knowledge sources used the symbolkeyed embeddings distributed with PRESAGE. The map was appended after the 40 default PRESAGE knowledge sources.

##### Evaluation

Models were evaluated on held-out test perturbations with the PRESAGE evaluation suite, using four tasks. Effect size was scored as the AUROC for distinguishing perturbations with a significant global transcriptional effect, ranked by the norm of the predicted response. Response direction was scored as the cosine similarity between predicted and observed responses, both centered on the mean training response, computed on the union of the top 20 differentially expressed genes of all perturbations, for perturbations with a significant effect. Phenocopy was scored as the recall of each training perturbation’s ten nearest test-set neighbors in observed response space among its ten nearest neighbors in predicted response space. Gene-set phenotypes were scored as the AUROC for identifying perturbations that strongly up- or down-regulate curated gene programs (mean of up and down).

To summarize these metrics into one value per knowledge source, we computed for each task and split the normalized gain (*s* − *s*_0_)*/*(*s*_ref_ − *s*_0_), where *s* is the score of the model, *s*_0_ the chance level, and *s*_ref_ the score of the reference. The chance level is an AUROC of 0.5, a cosine similarity of 0, and a recall of 10 divided by the number of test perturbations. The reference is the 40-source PRESAGE model on the same split, averaged over its three training seeds. The composite prior score is the mean of the four normalized scores, averaged over the four splits. It can be interpreted as the fraction of the 40-source model’s above-chance signal that a source achieves on its own.

The effect of adding a knowledge source to the 40-source model was assessed with the composite prior score of each of the 12 runs (4 splits × 3 seeds), paired with the 40-source model of the same split and seed. We used a two-sided Wilcoxon signed-rank test and a bootstrap 95% confidence interval of the mean difference (4000 resamples).

##### Experiments

For the single-source benchmark, one PRESAGE model was trained per knowledge source (each of the 40 default sources and the integrated map). To test the map in addition to the default sources, two configurations were compared with the 40source reference: adding the integrated map, and adding a gene-shuffled version of the map in which rows were permuted among covered genes, preserving coverage and the distribution of values. To compare the integrated map with its inputs, each of the 51 input maps was used as the only PRESAGE knowledge source. Its profile table entered PRESAGE with the default ingestion (*z*-scoring and principal component analysis to at most 128 components), and perturbation targets were matched to map proteins as for the integrated map.

##### Software

PRESAGE was run in its released environment (Python 3.11, PyTorch 2.2.0, PyTorch Geometric 2.5.2), with anndata 0.12.19 to read current AnnData files. The evaluation results of all models, and the code that derives the scores and statistics from them, are available in the analysis repository (see Data and Code Availability).

### 4.12 AI-assisted literature review and evidence curation

We combined AI-assisted literature searching and evidence curation with scripted retrieval to assemble localization evidence for proteins lacking database annotations. The search strategy and eligibility criteria are described below. Experimental context and source limitations were retained when interpreting the reported locations. This targeted review was limited by source coverage and access to full text.

#### 4.12.1 Localization evidence for proteins lacking database annotations

We reviewed all 400 integrated-map proteins lacking both a UniProt subcellular-location comment and a COMPARTMENTS knowledge-channel annotation rated at least four stars, through the Paperclip MCP interface with source=papers and all time=true, without a publication-date restriction. The literature review was conducted in two passes: a first pass over all 400 proteins, and a second that increased the hit limit for capped exact-phrase searches of proteins left without retained direct localization evidence for the human protein or a vertebrate ortholog. Claude (Anthropic) curator agents worked in 20 batches of 20 proteins in the first pass and 25 batches of 8 in the second. In the first pass each protein received an exact search on its quoted gene symbol (394 proteins) or, where the symbol was ambiguous or absent, on an alias or full protein name (6 proteins), and a query combining symbol and protein name with “subcellular localization” that included the abstract-only PubMed corpus. Adaptive follow-ups used aliases, former names, and hints from retrieved sources (1529 search calls, median 4 per protein). All queries are available in Supplementary Table S15. Hits present only as abstracts were retrieved as open-access full text or supplementary files through Europe PMC, PubMed Central, or publisher pages and reassessed by a follow-up agent curator.

Every first-pass search returned at most 10 hits, and 1021 of its 1343 completed searches (76%) reached that cap. Every protein was affected, with a median of two capped searches per protein. Of the 211 proteins without retained direct localization evidence for the human protein or a vertebrate ortholog after the first pass, 189 had at least one capped exact-phrase search and were included in the second pass. The remaining 22 had no capped exact-phrase search. The 241 eligible queries were re-issued with limit=200, exact=true, and all time=true, 6 of them with include abstracts=true. These queries returned 33,402 hits in total, with a maximum of 198 hits per query, so the 200-hit limit did not truncate any of the re-run result lists.

Evidence was screened at the experiment and construct level to retain direct localization measurements of the target protein: imaging of the endogenous or full-length tagged protein, fractionation or protease protection with specific detection, or proximity-ligation assays. Reviews, database records, predictions, and conference abstracts were excluded, as were expression-only, functional-only, interaction-only, and substrate- or partner-localization evidence, isolated targeting-peptide reporters, tissue staining without subcellular resolution, and activity-only fractions. Published primary abstracts were eligible when they named the target, a system of determinable species, the location, and a method or an explicit colocalization with a named marker, and were distinguished from full-text assessments. Separate verifier agents reopened every included source and checked metadata, protein identity, species and system, construct, method, compartment attribution, and excerpt. Corrections replaced curator fields, and rejected items were excluded (Supplementary Table S11).

The comparison used the map’s COMPARTMENTS-vocabulary assignment rather than its UniProt or Gene Ontology labels, the only one of the three that resolves the cytoplasm into distinct compartments (the UniProt vocabulary assigns 222 of the 400 proteins the single term Cytoplasm). The comparison reported in the Results was restricted to the 131 proteins with a maximum COMPARTMENTS-vocabulary class probability strictly greater than 0.8. Agreement was calculated among those with qualifying human-system evidence and comparable map assignments. Both sides were then projected onto a common comparison vocabulary by frozen tables: the map’s hierarchical labels by a fixed table, and the literature terms by a table fixed before the comparison, with curator free-text terms mapped by a further table written by the supervising agent after curation (Supplementary Table S16). All accepted compartments of a protein formed an unranked set. An assignment was consistent when its comparison compartment matched a literature compartment directly or through a predefined compatible pair. When no such match was present, a cytoplasmic map assignment was consistent at coarse resolution if the literature set included broad cytoplasm, even when specific cytoplasmic compartments were also reported. Direct or compatible matches took precedence over coarse matches. An assignment was inconsistent when a resolved, comparable map compartment met neither agreement criterion, and not comparable when the map label was unresolved or unharmonized or the literature had no map counterpart. The primary comparison used evidence for the human protein in human cells or tissues. A secondary comparison added human constructs in nonhuman cells and vertebrate orthologs. Both comparisons excluded proteomic-dataset evidence and publications flagged as inputs to the integrated map (Supplementary Table S10).

#### 4.12.2 Neuronal localization reference

The neuronal localization reference proteins and initial expected compartment labels were not derived from AI-assisted literature review, but instead from experimentally supported Gene Ontology annotations of human proteins to neuronal structures, each of which cites its publication. The expected labels for C9orf72 were subsequently amended after review of the cited neuronal evidence, as described in Section 4.6.3.

### 4.13 The *grassp* portal

#### 4.13.1 Implementation and dataset catalog

The *grassp* portal (https://grassp.apps.biohub.org/) is a web application written in TypeScript with Next.js and React, with an interface built on our in-house design system, the Science Design System (https://github.com/chanzuckerberg/sci-components). The curated .h5ad objects are stored in cloud object storage and can be downloaded directly from the portal. When an object is added or updated, its map-, sample-, and protein-level metadata are imported into a MongoDB database, and the three-dimensional UMAP embedding stored in the object is converted for in-browser visualization.

The catalog filters (Fig. 5a) draw on the curated metadata fields (Section 4.9.8): organism, tissue (the curated specific term), sample class (cell line, tissue, or primary cell culture), cell line, disease state, and enrichment strategy. These are complemented by the publication year from the CrossRef metadata and by two counts computed from each object: the number of quantified proteins and the number of samples (fractions or baits). Keyword search performs a case-insensitive match of the query against the dataset identifier, publication title and authors, organism, and enrichment strategy.

The protein-centric report (Fig. 5c) is assembled at query time from two sources. Protein identity, curated subcellular-location terms, and cross-references come from reviewed UniProtKB/Swiss-Prot entries retrieved through the UniProt REST API. The Ensembl gene identifier links to external resources such as the Human Protein Atlas and OpenCell. Localization evidence is aggregated across every map in which the protein was quantified: the *grassp* independent diffusion annotations under the consolidated or the full UniProt vocabulary and the author labels (Section 4.9.6) are counted per term to produce the two word clouds, and the protein’s compartment scores in the integrated pan-human map (Section 4.10) populate the “Integrated Pan-Human Map” view. The same catalog and protein queries are available programmatically through a REST API.

The portal runs as containerized services on Kubernetes on Amazon Web Services.

#### 4.13.2 Natural-language querying

The chatbot (Fig. 5d) is the Biohub Chatbot (https://chatbot.apps.czbiohub.org), configured with a *grassp*-specific agent and backed by Anthropic’s Claude Opus 4.8. The agent accesses portal data through a Model Context Protocol (MCP) server hosted by the portal, which lets the agent inspect the database schema and query map, sample, and protein records. All queries connect to the database with read-only credentials. A second MCP server lets the agent search the full text of the publications behind the collection’s maps, so it can answer methodological questions not captured in the curated metadata. The agent’s instructions also describe the *grasspViz* URL scheme, so answers can include stateful links that open a map with the relevant proteins highlighted.

For analyses that go beyond database queries, the agent writes Python code that is executed in an isolated cloud sandbox. The sandbox provides the standard Python data science stack (NumPy, pandas, SciPy, scikit-learn, matplotlib, and seaborn), together with *anndata*, *scanpy*, and the *grassp* library itself, so the agent can compute statistics, generate plots, and apply *grassp* functions to portal data. To stay within the model’s limited context window, large query results are not passed to the model. Instead, the portal writes them directly into the sandbox as a file, and the model receives only the number of records and the file size. The agent then analyzes the full result set with code, so large exports, such as protein-level records across many maps, never enter the model’s context. Tables, figures, and result files produced in the sandbox are returned to the user for download. Querying and narrative answers are available without an account. Code execution and downloads require a free account registered with a university or research-organization email address.

### 4.14 grasspViz

*grasspViz* is a Streamlit application written in Python that enables interactive exploration of a subcellular map in AnnData format stored as an .h5ad file. Files can either be uploaded by the user or fetched from the *grassp* data portal. The application can be installed locally using conda or Docker, or accessed via the public web interface at https://grasspviz.apps.biohub.org/.

Scatter plots, violin plots, heatmaps, and bar plots are drawn with Plotly. The protein-embedding and volcano plots support point, box, and lasso selection. The protein-neighborhood graph is drawn with Cytoscape.js [104] and supports click and box selection.

The state of the application is encoded in the URL query string, so any prepared view is shareable as a link.

*grasspViz* supports gene-set enrichment calculation on a selected group of proteins. Enrichment is computed with a one-tailed hypergeometric test, using the proteins of the loaded object as the background reference proteome. *P* values are adjusted within each library by the Benjamini–Hochberg procedure. Gene-set libraries are retrieved from Enrichr [105]. Human, mouse, yeast, fly, fish, and worm libraries are available.

The source code for *grasspViz* is available at https://doi.org/10.5281/zenodo.24484249.

## 5 Supplementary information

The supplementary figures are provided in a separate Supplementary Information document. The supplementary tables listed below are provided as a single Excel workbook (Supplementary Tables.xlsx) with one sheet per table, named by its table number, and a contents sheet.

## List of Supplementary Tables

**Supplementary Table S1. – Benchmark datasets.** The 12 published human sub-cellular maps used for the annotation benchmarks of Fig. 2g–i, with portal identifiers, publication details, and curated experimental metadata.

**Supplementary Table S2 – iNeuron map annotations.** Localization annotations of the 5973 proteins of the iNeuron OrgIP map generated in this study (portal dataset `Frank_2026_iNeuron`).

**Supplementary Table S3. – Neuronal localization reference.** GO-derived neuronal localization reference for the 17 proteins detected in the iNeuron map shown in Fig. 3c and Supplementary Table S4, with the expected compartments, their publication support, and the iNeuron map assignments.

**Supplementary Table S4. – iNeuron localization comparison.** Comparison of *grassp* compartment assignments with GO-derived expected labels for the 12 human reference proteins detected in the iNeuron map with neuronal-structure support from exactly two publications, including at least one supporting a mapped compartment. The table provides compartment labels, match, coarse-match, and no-match categories, and supporting references. The five reference proteins with three or more publications are shown in Fig. 3c.

**Supplementary Table S5 – HEK293T DC map annotations.** Localization annotations of the 8595 proteins of the HEK293T differential centrifugation map generated in this study (portal dataset `Frank_2026_HEK293`).

**Supplementary Table S6. – Curated dataset collection.** The 103 subcellular maps curated for the *grassp* portal, with publication details and curated experimental metadata given as ontology identifiers and labels.

**Supplementary Table S7. – Mitochondrial island.** The 849 proteins of the mitochondrial island of the integrated pan-human map, with *grassp* localization calls, MitoCarta3.0, MitoCoP, and UniProt annotations, HEK293T OrgIP and fractionation measurements, and Leiden clusters.

**Supplementary Table S8. – Mito-cytosolic cluster.** The 110 proteins of the Mitocytosolic cluster of the integrated pan-human map, with the fields of Supplementary Table S7 and cluster tier assignments.

**Supplementary Table S9. – SurfaceBench proteins in the integrated panhuman map.** The 147 proteins of the SurfaceBench benchmark of cell-surface accessibility with their benchmark labels and, for the 80 proteins in the integrated pan-human map, the independent diffusion memberships and top compartments in the full UniProt subcellular-location hierarchy, computed with the benchmark proteins held out of the gene sets, and the canonical locations of the conditional proteins read from the benchmark rationales.

**Supplementary Table S10. – Proteins lacking database localization annotations.** Integrated-map assignments and literature-review outcomes for the 400 proteins lacking both a UniProt subcellular-location annotation and a high-confidence COMPARTMENTS assignment.

**Supplementary Table S11. – Publications assessed for proteins lacking database annotations.** The 1666 protein–publication records assessed in the literature review of Supplementary Table S10, with evidence classification, verbatim excerpts, and curator and verifier decisions.

**Supplementary Table S12. – Knowledge sources for perturbation-response prediction.** Composite prior scores, normalized gains in the four prediction tasks, and evaluation metrics of PRESAGE models trained with a single knowledge source: each of the 40 default sources of PRESAGE and the integrated pan-human map (Supplementary Fig. S10a), and each of the 51 input maps of the integrated map (Supplementary Fig. S10c).

**Supplementary Table S13. – Figure dataset names.** Correspondence between the short dataset names used in the figures and the dataset identifiers of the *grassp* portal (113 figure names resolving to 88 portal datasets and the integrated pan-human map).

**Supplementary Table S14. – iNeuron OrgIP bait cell lines.** The 14 endogenously tagged iPSC lines used as OrgIP baits in iNeurons, with the targeted compartment, the tagged terminus, and the sgRNA and donor template sequences.

**Supplementary Table S15. – Literature-search records.** The 4239 search queries and source-retrieval records of the AI-assisted literature review of the proteins lacking database annotations.

**Supplementary Table S16. – Comparison vocabulary.** Projections of map labels and literature-reported localizations onto the common comparison vocabulary, with the compatible compartment pairs and broad-cytoplasm compatibility rules used to score agreement.

## Data and Code Availability

Code for the *grassp* framework is available at: https://github.com/czbiohub-sf/grassp.

Code for the *grasspio* R package is available as a subrepository of the *grassp* repository at: https://github.com/czbiohub-sf/grassp/tree/main/r/grasspio.

Documentation and tutorials for *grassp* are available at: https://czbiohub-sf.github.io/grassp/tutorials/index.html.

Code to reproduce analyses and figures from this publication is available at: https://doi.org/10.5281/zenodo.24484247.

Code for *protdata*, the underlying mass spectrometry data-handling library, is available at: https://github.com/czbiohub-sf/protdata.

Code for the curation of the *grassp* collection is available at: https://doi.org/10.5281/zenodo.24484245.

Code for the *grasspViz* interactive web-explorer is available at: https://doi.org/10.5281/zenodo.24484249.

Raw data for the iNeuron OrgIP map and the HEK293T DC map will be made available upon publication.

## Acknowledgements

We thank the Biohub and its donors, Priscilla Chan and Mark Zuckerberg, for funding this work. We thank Carlos Gonzales and Sarah Lin for help with processing of raw mass spectrometry data. We thank Aline Tschanz for help with illustrations in this manuscript.

## Declarations

### Author contributions

Conceptualization: M.F., M.D.L., and D.P.; methodology: M.F. and D.P.; software: M.F. (*grassp* and *grasspViz*), K.A. (*grassp* portal), and L.L. (*grasspViz*); portal design: K.A., M.F., and D.P.; data curation: M.F.; formal analysis: M.F., M.C., and D.P.; investigation: S.V., M.B., F.M., J.T., B.C.D., C.J., R.B.-N., Z.Y., M.V., L.N., L.S., and D.N.I.; visualization: M.F. and D.P.; writing: M.F., M.D.L., and D.P.; supervision: J.E.E., M.D.L., and D.P.

### Use of generative AI and AI-assisted technologies

During the preparation of this work, the authors used ChatGPT/Codex (OpenAI) and Claude/Claude Code (Anthropic) to assist with writing code, to perform literature searches, and to edit the text. After using these tools, the authors reviewed and edited the content as needed and take full responsibility for the content of the published article.

