## Supplemental Figures for "grassp: A graph-based framework unifies subcellular proteomics across assays and studies"

### Supplementary Figures

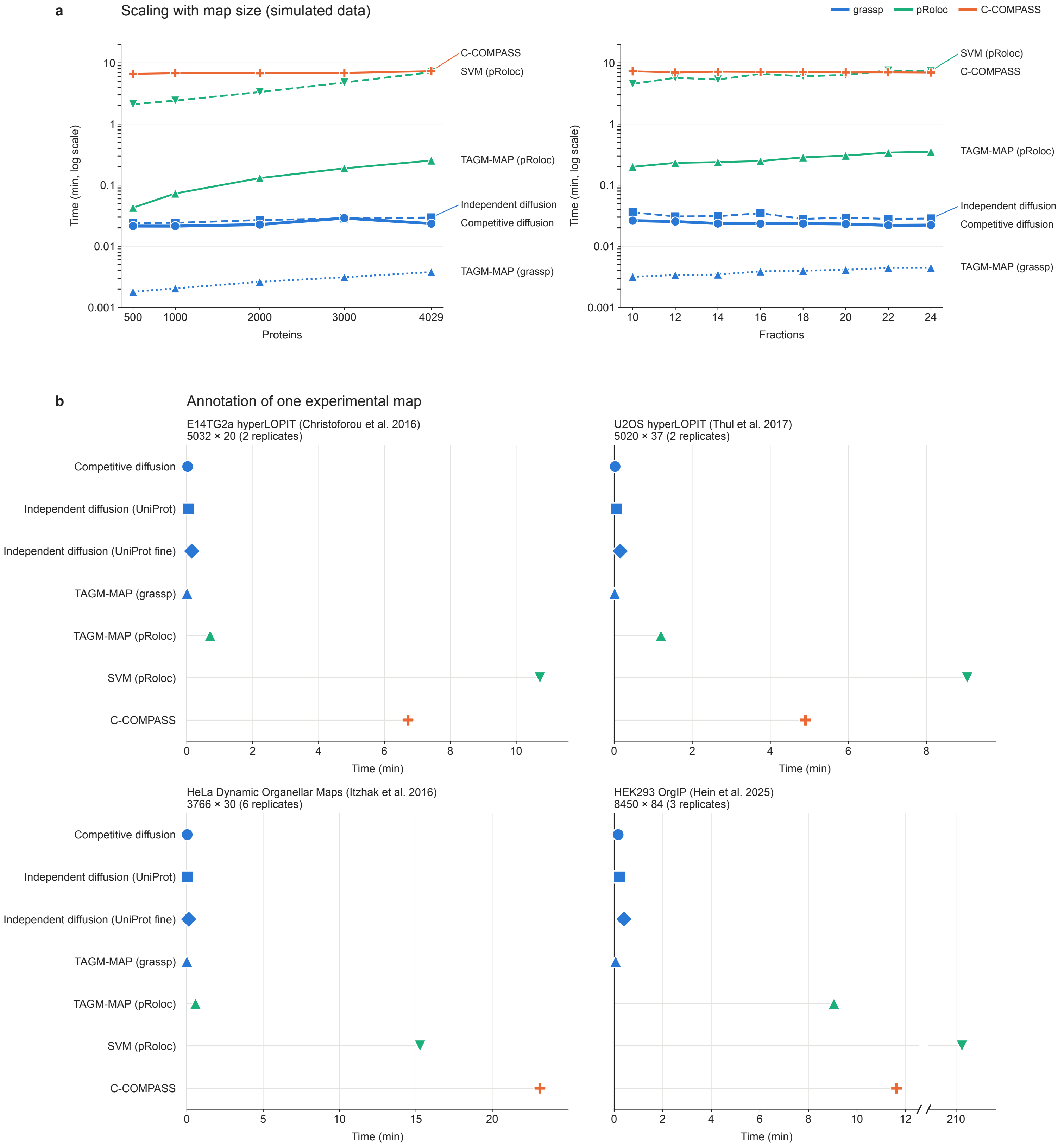
a
Scaling with map size (simulated data)
grassp
pRoloc
C-COMPASS
Time (min, log scale)
10
1
0.1
0.01
0.001
500
1000
2000
3000
4029
Proteins
C-COMPASS
SVM (pRoloc)
TAGM-MAP (pRoloc)
Independent diffusion
Competitive diffusion
TAGM-MAP (grassp)
Time (min, log scale)
10
12
14
16
18
20
22
24
Fractions
SVM (pRoloc)
C-COMPASS
TAGM-MAP (pRoloc)
Independent diffusion
Competitive diffusion
TAGM-MAP (grassp)
b
Annotation of one experimental map
E14TG2a hyperLOPIT (Christoforou et al. 2016)
5032 × 20 (2 replicates)
U2OS hyperLOPIT (Thul et al. 2017)
5020 × 37 (2 replicates)
Competitive diffusion
Independent diffusion (UniProt)
Independent diffusion (UniProt fine)
TAGM-MAP (grassp)
TAGM-MAP (pRoloc)
SVM (pRoloc)
C-COMPASS
0
2
4
6
8
10
Time (min)
HeLa Dynamic Organellar Maps (Itzhak et al. 2016)
3766 × 30 (6 replicates)
HEK293 OrgIP (Hein et al. 2025)
8450 × 84 (3 replicates)
0
5
10
15
20
12
210
Time (min)

**Supplementary Fig. S1** Competitive diffusion annotates a subcellular map in seconds, faster than every published implementation tested. On the experimental maps, independent diffusion used either the 24 consolidated UniProt subcellular-location terms (UniProt) or the full hierarchy (UniProt fine). Color gives the implementation (*grassp*, blue; pRoloc, green; C-COMPASS, orange), and within one implementation, line style and marker distinguish the methods. **(a)** Time (log scale) against map size on the simulated data of Haas et al. [1]. Left: the simulation of Supplementary Fig. S2 (8 compartments, 16 fractions) subsampled to 500, 1000, 2000, 3000, and all 4029 proteins. Right: the authors' simulations with 8 compartments and 10 to 24 fractions (3916–4129 proteins). Competitive diffusion took 1.3–1.7 s at every size and was at least twice as fast as every published implementation. **(b)** Annotation of four experimental maps: mouse embryonic stem cell hyperLOPIT [2], 5032 proteins, 20 fractions, 926 markers; U2-OS hyperLOPIT [3], 5020 proteins, 37 fractions, 638 markers; HeLa Dynamic Organellar Maps [4], 3766 proteins, 30 fractions, 1018 markers; and HEK293T organelle immunoprecipitation (OrgIP) [5], 8450 proteins, 84 pulldowns, 1921 markers. Competitive diffusion took 1.4–9.7 s per map and was 23–1310 times faster than the published implementations (TAGM-MAP in pRoloc 34 s–9.0 min, C-COMPASS 4.9–23.1 min, SVM 9.0 min–3.5 h). Its first run is dominated by one-time just-in-time compilation, and repeated runs took 0.04–0.73 s. The TAGM-MAP implementation of *grassp* took 0.42–3.7 s. Independent diffusion took 2.7–12.8 s with the consolidated UniProt vocabulary and 7.1–24.1 s with the full hierarchy. Time axes are linear and scaled to each map, and the HEK293T OrgIP axis is broken so that SVM (pRoloc), at 3.5 h, does not compress the other methods.

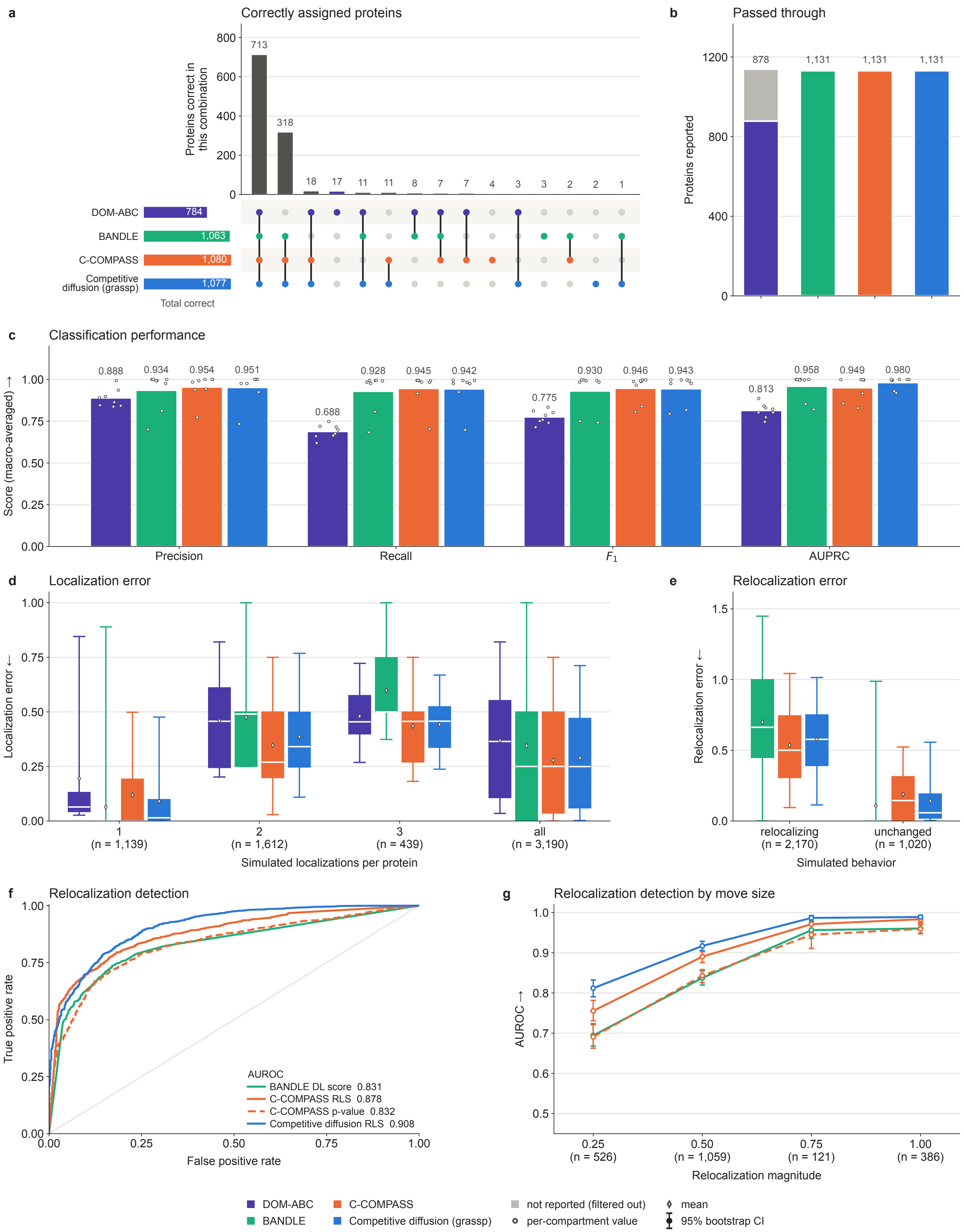
a
Correctly assigned proteins
Proteins correct in this combination
800
600
400
200
0
713
318
18
17
11
11
8
7
7
4
3
3
2
2
1
DOM-ABC 784
BANDLE 1,063
C-COMPASS 1,080
Competitive diffusion (grassp) 1,077
Total correct
b
Passed through
Proteins reported
1200
800
400
0
878
1,131
1,131
1,131
c
Classification performance
Score (macro-averaged) →
1.00
0.75
0.50
0.25
0.00
0.888
0.934
0.954
0.951
0.688
0.928
0.945
0.942
0.775
0.930
0.946
0.943
0.813
0.958
0.949
0.980
Precision
Recall
$F_1$
AUPRC
d
Localization error
Localization error ←
1.00
0.75
0.50
0.25
0.00
1 (n = 1,139)
2 (n = 1,612)
3 (n = 439)
all (n = 3,190)
Simulated localizations per protein
e
Relocalization error
Relocalization error ←
1.5
1.0
0.5
0.0
relocalizing (n = 2,170)
unchanged (n = 1,020)
Simulated behavior
f
Relocalization detection
True positive rate
1.00
0.75
0.50
0.25
0.00
False positive rate
0.00
0.25
0.50
0.75
1.00
AUROC
BANDLE DL score 0.831
C-COMPASS RLS 0.878
C-COMPASS p-value 0.832
Competitive diffusion RLS 0.908
g
Relocalization detection by move size
AUROC →
1.0
0.9
0.8
0.7
0.6
0.5
0.25 (n = 526)
0.50 (n = 1,059)
0.75 (n = 121)
1.00 (n = 386)
Relocalization magnitude
DOM-ABC
BANDLE
C-COMPASS
Competitive diffusion (grassp)
not reported (filtered out)
per-compartment value
mean
95% bootstrap CI

**Supplementary Fig. S2** Competitive diffusion matches or outperforms published spatial proteomics classifiers on a published simulated benchmark dataset. Data and annotation results for existing tools and benchmark metrics were taken from Haas et al. [1], and only competitive diffusion (*grassp*) was computed here. Arrows on axis labels point in the direction of better performance. **(a)** UpSet plot of the 1139 non-marker proteins the simulation assigns to a single compartment. Vertical bars give the size of each combination, sorted by size. The matrix below identifies the methods making up each combination, and the horizontal bars on the left the total correct per method. All methods largely agree on labels for proteins with a single localization. **(b)** Number of proteins for which each method reports a compartment. Gray marks proteins filtered out by the method. **(c)** Precision, recall, $F_1$ and area under the precision–recall curve (AUPRC), over the eight simulated compartments. Numbers above the bars are the macro-averaged scores and points represent the score per compartment. Note that the scores for DOM-ABC are artificially lowered because of the filtering. **(d)** Localization error, defined as in Haas et al. [1] as half the summed absolute difference, between a method's predicted compartment vector and the simulated ground truth, shown separately by the number of compartments the simulation assigned to each protein (1139, 1612, and 439 proteins with one, two, and three compartments) and pooled over all 3190 proteins. Boxes represent quartiles, white lines medians, diamonds means and whiskers the 5th and 95th percentiles. **(e)** The same statistic applied to the change in compartment vector between conditions 1 and 2 against the simulated change, split by whether the simulation moved the protein (2170 moved and 1020 unchanged proteins). BANDLE scores all of them, whereas C-COMPASS and competitive diffusion report relocalization for 2092 moved and 1002 unchanged proteins. DOM-ABC is absent as in the original publication. **(f)** Receiver operating characteristic curves for detecting whether the simulation moved a protein, using each tool's relocalization statistic, on the 3094 proteins for which all four statistics are available (2092 moved, 1002 unchanged). Statistics used were: BANDLE's differential localization probability (bandle.differential.localisation); C-COMPASS's relocalization score (RLS), the summed absolute change in class contributions; the negative C-COMPASS relocalization $P$ value; and the same RLS statistic computed from the compartment probabilities of competitive diffusion. **(g)** The same area under the receiver operating characteristic curve (AUROC) computed separately for each size of simulated move (526, 1059, 121, and 386 moved proteins per bin), against the same 1002 unchanged proteins. Error bars are 95% bootstrap intervals over proteins with 500 resamples.

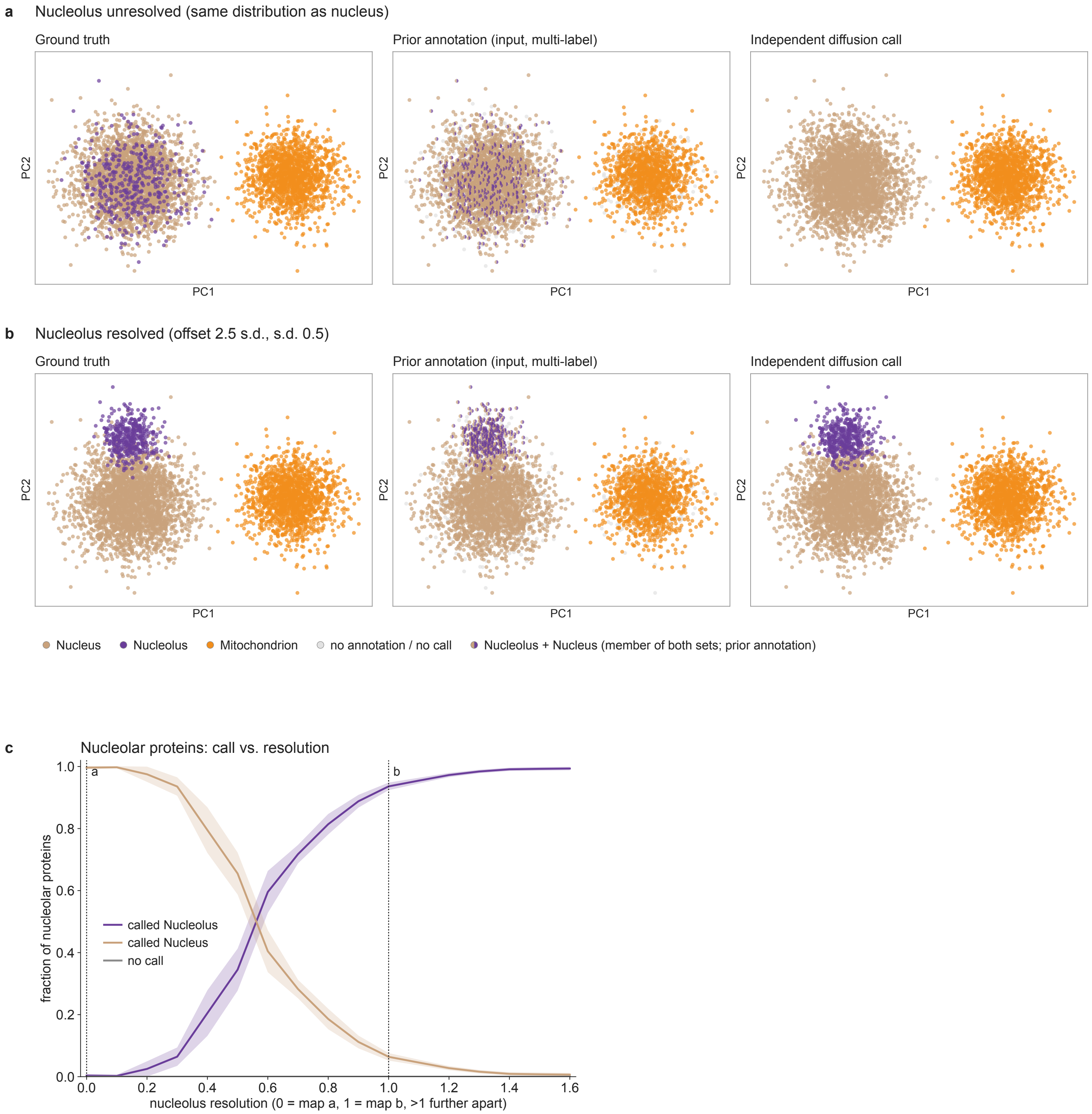

a
Nucleolus unresolved (same distribution as nucleus)
Ground truth
Prior annotation (input, multi-label)
Independent diffusion call
PC1
PC2
b
Nucleolus resolved (offset 2.5 s.d., s.d. 0.5)
Ground truth
Prior annotation (input, multi-label)
Independent diffusion call
Nucleus
Nucleolus
Mitochondrion
no annotation / no call
Nucleolus + Nucleus (member of both sets; prior annotation)
c
Nucleolar proteins: call vs. resolution
fraction of nucleolar proteins
called Nucleolus
called Nucleus
no call
nucleolus resolution (0 = map a, 1 = map b, >1 further apart)
0.0
0.2
0.4
0.6
0.8
1.0
1.2
1.4
1.6

**Supplementary Fig. S3** Independent diffusion annotates a subcompartment only where the map resolves it (simulation). Three isotropic Gaussian compartments were simulated in 6 dimensions (2800 nuclear, 400 nucleolar, and 1600 mitochondrial proteins), and 80% of proteins were given a correct prior annotation with 20% unannotated. The vocabulary is hierarchical, as in UniProt or GO: an annotated nucleolar protein is a member of the Nucleolus and of the Nucleus set. A protein-neighborhood graph ($k = 20$) was built on the six fractions and annotated with *grassp* (`grassp.tl.independent_diffusion`, default settings), which diffuses each term one-vs-rest, calibrates the result to a per-term membership probability, and labels each protein with the most specific term that explains its membership profile. Panels **a** and **b** each show one simulated map in the same principal component projection, as ground truth (left), prior annotation (middle), and independent diffusion annotation (right). **(a)** Map with unresolved nucleolus: nucleolar proteins are drawn from the same distribution as the rest of the nucleus. In the prior annotation vocabulary (middle), an annotated nucleolar protein is a member of the Nucleolus set *and* of the Nucleus set, and is drawn as a circle split half tan, half purple. Gray proteins are unannotated. In the independent diffusion annotation (right), all ground-truth nucleolar proteins are called Nucleus (100% Nucleus, 0% Nucleolus), because no neighborhood is enriched for Nucleolus members specifically. The well-separated mitochondrion is called correctly (100%). **(b)** The resolved map: the same simulated proteins, with the nucleolar ones forming a tighter cloud. The prior annotation (middle) is identical to that in **a**. Now 95% of nucleolar proteins are called Nucleolus and 5% Nucleus, while other nuclear proteins are essentially never called Nucleolus (2%). One protein in each map received no call (gray, right). **(c)** The transition between the two regimes. The nucleolar cloud was moved in steps of 0.1 from the unresolved geometry of **a** (resolution 0) to the resolved geometry of **b** (resolution 1), its offset growing and its spread shrinking linearly, and on to resolution 1.6, where it keeps the resolved spread and sits 4 s.d. from the nucleus center (5 independent simulations per step; line, mean; band, $\pm$ s.d.). Shown is the fraction of nucleolar proteins called Nucleolus, Nucleus, or unannotated. The Nucleolus call is absent while the nucleolus is buried in the nucleus and only appears with increasing separation. We refer to this as automatic resolution detection, where more specific terms only win over their parent terms when there is enough evidence for them in the data.

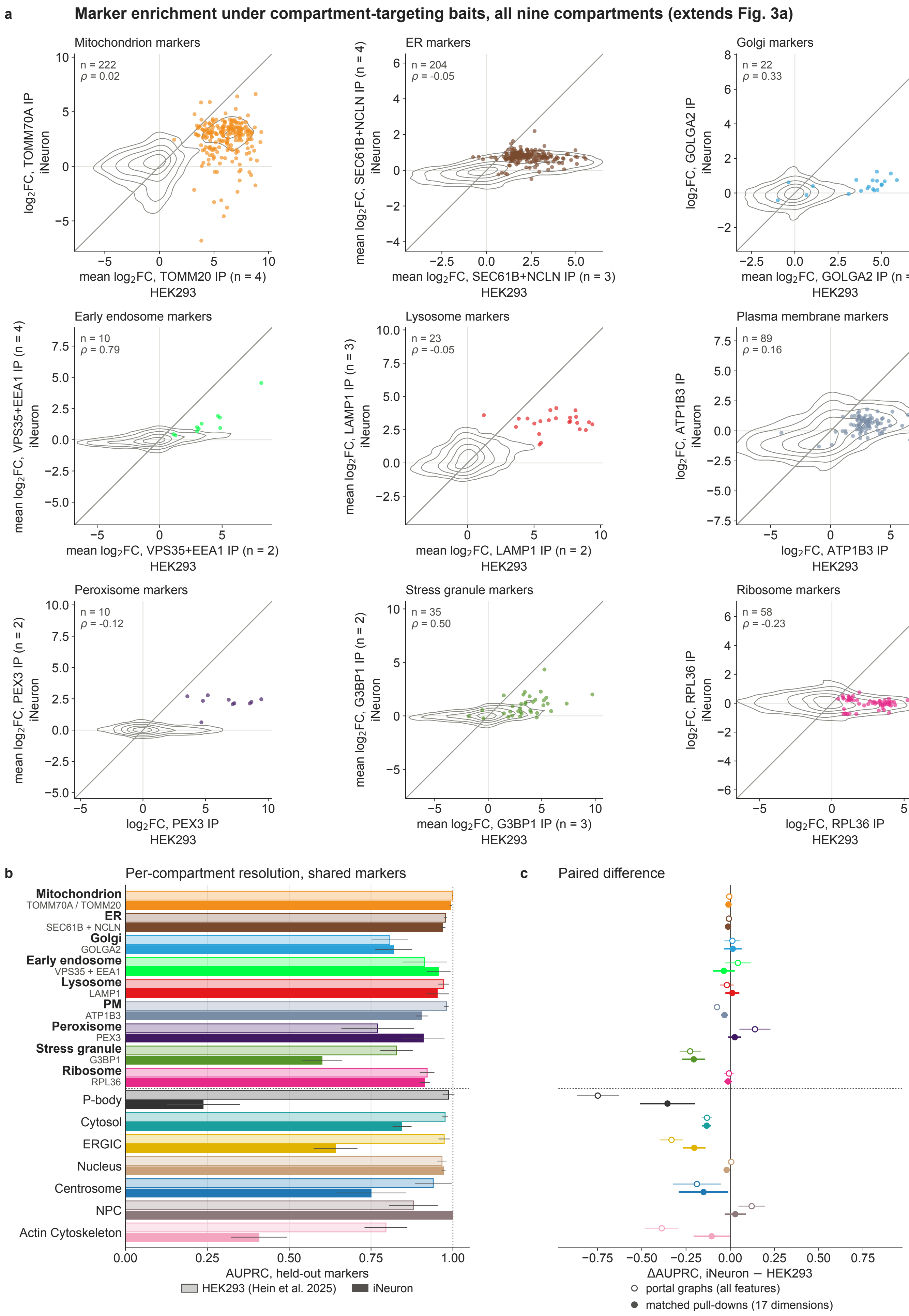
a
Marker enrichment under compartment-targeting baits, all nine compartments (extends Fig. 3a)
Mitochondrion markers
n = 222
ρ = 0.02
log2FC, TOMM70A IP
iNeuron
mean log2FC, TOMM20 IP (n = 4)
HEK293
ER markers
n = 204
ρ = -0.05
mean log2FC, SEC61B+NCLN IP (n = 4)
iNeuron
mean log2FC, SEC61B+NCLN IP (n = 3)
HEK293
Golgi markers
n = 22
ρ = 0.33
log2FC, GOLGA2 IP
iNeuron
mean log2FC, GOLGA2 IP (n = 4)
HEK293
Early endosome markers
n = 10
ρ = 0.79
mean log2FC, VPS35+EEA1 IP (n = 4)
iNeuron
mean log2FC, VPS35+EEA1 IP (n = 2)
HEK293
Lysosome markers
n = 23
ρ = -0.05
mean log2FC, LAMP1 IP (n = 3)
iNeuron
mean log2FC, LAMP1 IP (n = 2)
HEK293
Plasma membrane markers
n = 89
ρ = 0.16
log2FC, ATP1B3 IP
iNeuron
log2FC, ATP1B3 IP
HEK293
Peroxisome markers
n = 10
ρ = -0.12
mean log2FC, PEX3 IP (n = 2)
iNeuron
log2FC, PEX3 IP
HEK293
Stress granule markers
n = 35
ρ = 0.50
mean log2FC, G3BP1 IP (n = 2)
iNeuron
mean log2FC, G3BP1 IP (n = 3)
HEK293
Ribosome markers
n = 58
ρ = -0.23
log2FC, RPL36 IP
iNeuron
log2FC, RPL36 IP
HEK293
b
Per-compartment resolution, shared markers
Mitochondrion
TOMM70A / TOMM20
ER
SEC61B + NCLN
Golgi
GOLGA2
Early endosome
VPS35 + EEA1
Lysosome
LAMP1
PM
ATP1B3
Peroxisome
PEX3
Stress granule
G3BP1
Ribosome
RPL36
P-body
Cytosol
ERGIC
Nucleus
Centrosome
NPC
Actin Cytoskeleton
0.00
0.25
0.50
0.75
1.00
AUPRC, held-out markers
HEK293 (Hein et al. 2025)
iNeuron
c
Paired difference
−0.75
−0.50
−0.25
0.00
0.25
0.50
0.75
ΔAUPRC, iNeuron − HEK293
portal graphs (all features)
matched pull-downs (17 dimensions)

**Supplementary Fig. S4** The iNeuron organelle immunoprecipitation (OrgIP) map against the HEK293T OrgIP reference. Both maps can be obtained on the *grassp* data portal: `Frank_2026_iNeuron` (5973 proteins) and `Hein_2025_HEK293_WT` (8450 proteins). In total, 3750 proteins are quantified under the same UniProt accession in both maps, and 981 of these carry curated compartment marker labels from Hein et al. [5]. **(a)** Comparison plot of $\log_2$ OrgIP enrichment of shared marker proteins in pulldowns targeting a compartment, for nine compartments with baits in both maps. This extends Fig. **??**a, which shows three of them. Marker proteins for the compartment are shown as points. When a compartment had several pulldowns, the enrichment values were averaged. The diagonal line marks equal enrichment. Gray contours are the kernel density of the non-marker shared proteins under the same baits (all 3750 shared proteins except the plotted markers). Enrichment was systematically lower in iNeurons for 649 of 673 markers (below the diagonal). Median $\log_2$ fold changes, iNeuron versus HEK293T, were 3.0 versus 6.2 (Mitochondrion), 0.7 versus 2.1 (endoplasmic reticulum), 0.4 versus 4.6 (Golgi), 1.0 versus 3.2 (Early endosome), 3.1 versus 6.6 (Lysosome), 0.6 versus 2.9 (Plasma membrane), 2.3 versus 7.0 (Peroxisome), 0.9 versus 3.5 (Stress granule), and 0.0 versus 2.9 (Ribosome). **(b)** Per-compartment resolution of the two maps, measured by held-out marker area under the precision–recall curve (AUPRC). The 981 shared markers were split into stratified 5-fold cross-validation, repeated five times (identical folds for both maps). AUPRC was measured on held-out markers after *grassp* competitive diffusion ($\alpha = 0.5$) over each map's connectivity graph. Bars represent mean AUPRC over 25 folds; whiskers, 95% confidence interval. The bold compartments correspond to panel **a**. Macro-averaged AUPRC was 0.918 for HEK293T and 0.805 for iNeurons. Over the nine compartments with baits in both maps (bold), it was 0.908 for HEK293T and 0.891 for iNeurons, a difference of 0.016. **(c)** The paired difference of the per-compartment AUPRC, iNeuron minus HEK293T, over the same 25 folds (point, mean; line, 95% confidence interval). Open markers use each map's published graph as in **b**. The iNeuron map scored lower in 11 of 16 compartments. Filled markers repeat the comparison at matched pulldowns, where both maps were reduced to the 17 shared dimensions. Restricting the HEK293T map to the shared pulldowns changes the macro-averaged AUPRC difference from 0.113 to 0.075 (HEK293T 0.870, iNeuron 0.796).

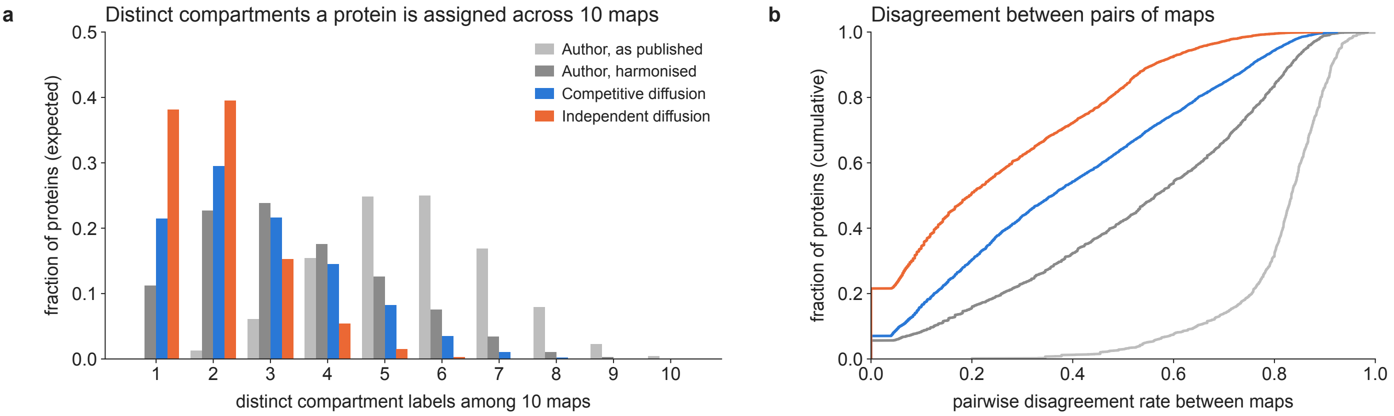
a
Distinct compartments a protein is assigned across 10 maps
fraction of proteins (expected)
distinct compartment labels among 10 maps
Author, as published
Author, harmonised
Competitive diffusion
Independent diffusion
0.0
0.1
0.2
0.3
0.4
0.5
1
2
3
4
5
6
7
8
9
10
b
Disagreement between pairs of maps
fraction of proteins (cumulative)
pairwise disagreement rate between maps
0.0
0.2
0.4
0.6
0.8
1.0

**Supplementary Fig. S5** Consistency of compartment labels across maps. Labels from four annotation strategies were compared across the 51 unperturbed human maps, for the 5665 proteins that each strategy labels in at least 10 maps (Methods): the authors' annotation as published, the same annotation after harmonization, competitive diffusion of the marker list of Hein et al. [5], and independent diffusion of the UniProt subcellular-location terms. **(a)** Number of distinct compartment labels a protein receives across 10 maps. Bars show the probability of each number over all draws of 10 of a protein's labeled maps, averaged over proteins. The expected number of distinct labels per protein averages 5.59 (author, as published), 3.36 (author, harmonized), 2.20 (competitive diffusion), and 2.06 (independent diffusion). Compared protein by protein, independent diffusion yields fewer distinct compartments than the harmonized author annotation for 88% of proteins, and competitive diffusion for 82%. **(b)** Cumulative distribution, over the same proteins, of the fraction of pairs of a protein's labeled maps that assign different labels (mean 0.81, 0.52, 0.28, and 0.27, same order as in **a**).

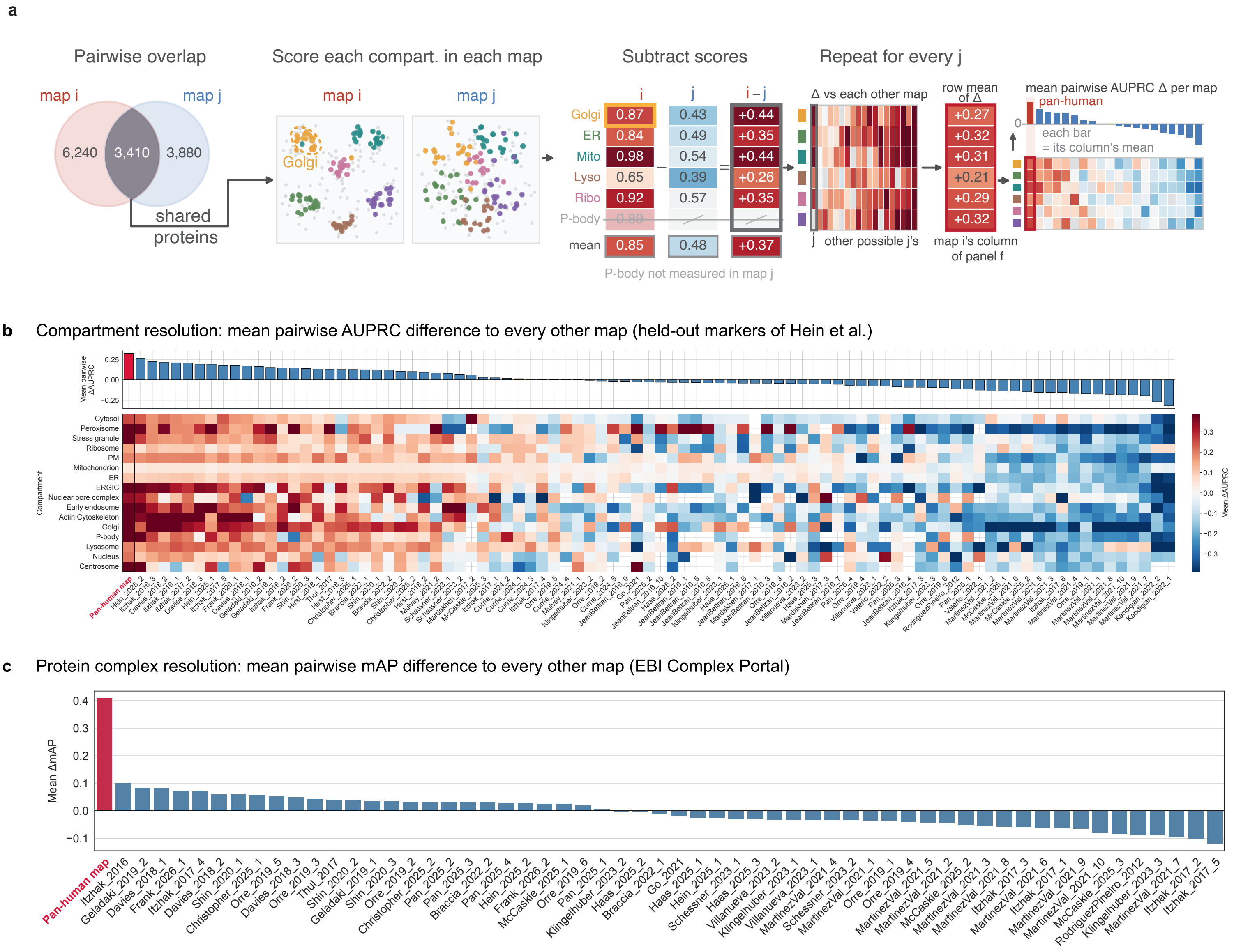
a
Pairwise overlap
map i
map j
6,240
3,410
3,880
shared
proteins
Score each compart. in each map
map i
map j
Golgi
Subtract scores
i
j
i – j
Golgi 0.87 0.43 +0.44
ER 0.84 0.49 +0.35
Mito 0.98 0.54 +0.44
Lyso 0.65 0.39 +0.26
Ribo 0.92 0.57 +0.35
P-body 0.80
mean 0.85 0.48 +0.37
P-body not measured in map j
Repeat for every j
Δ vs each other map
j other possible j's
row mean of Δ
+0.27
+0.32
+0.31
+0.21
+0.29
+0.32
map i's column of panel f
mean pairwise AUPRC Δ per map
pan-human
0
each bar = its column's mean
b
Compartment resolution: mean pairwise AUPRC difference to every other map (held-out markers of Hein et al.)
Mean pairwise ΔAUPRC
Compartment
Cytosol
Peroxisome
Stress granule
Ribosome
PM
Mitochondrion
ER
ERGIC
Nuclear pore complex
Early endosome
Actin Cytoskeleton
Golgi
P-body
Lysosome
Nucleus
Centrosome
Mean ΔAUPRC
Pan-human map
c
Protein complex resolution: mean pairwise mAP difference to every other map (EBI Complex Portal)
Mean ΔmAP
Pan-human map

**Supplementary Fig. S6** Marker-protein prediction performance of the integrated pan-human map compared with the individual input datasets, supplementing Fig. **??**. **(a)** Pairwise procedure used to benchmark compartment resolution. For each pair of maps $i$ and $j$, scoring is restricted to their shared proteins. The curated markers on the shared pool are split into five stratified folds, identical for both maps, and each fold's training markers seed competitive diffusion over each map's own graph. A held-out area under the precision–recall curve (AUPRC) is calculated for each measured compartment in each map from the pooled out-of-fold predictions, and the difference $\Delta\text{AUPRC} = \text{AUPRC}_i - \text{AUPRC}_j$ is recorded. Compartments with fewer than five shared markers are omitted. Repeating the comparison against every other map $j$ gives a mean pairwise difference for each compartment in map $i$. These compartment-specific means form one column of the heatmaps in Fig. **??**f and in panel **b**. The bar above that column averages the same differences first over the compartments of a pair and then over the comparison maps (Methods). The analogous pairwise-difference procedure is used with complex mean average precision in Fig. **??**e and panel **c**. **(b)** Compartment resolution of the integrated map and of every individual dataset, scored by held-out marker AUPRC (marker set shown: the curated markers of Hein et al. [5], 16 compartments). Top: mean pairwise AUPRC difference of each map against all other maps, sorted, with the integrated map (crimson) first (rank 1/89). Positive values indicate that the indicated map resolves markers better, on average, than the maps it is compared with. Bottom: the same difference resolved per compartment (red, indicated map better; blue, worse). The dataset names on the axes of **b** and **c** are resolved to portal identifiers in Supplementary Table **??**. **(c)** Protein-complex resolution of the integrated map and of every individual dataset with sufficient complex coverage, scored as the pairwise mean average precision (mAP) with which each map's graph ranks the members of a complex next to each member in turn (protein complexes were obtained from the EBI Complex Portal). Mean pairwise mAP difference per dataset, sorted. Positive values indicate that a map ranks complex members closer together than the maps it is compared with. The integrated map (crimson) ranks 1/59, colocalizing complex members more tightly than any single input dataset.

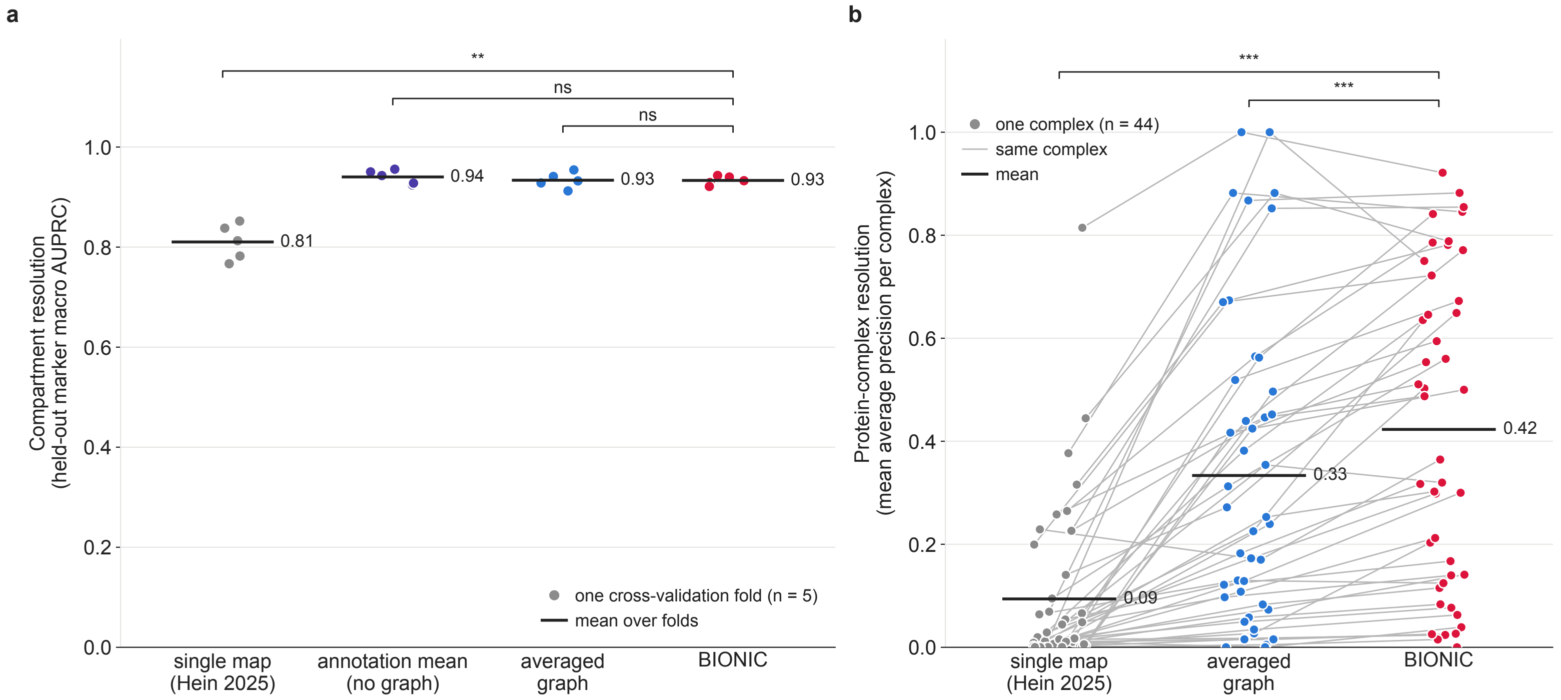
a
**
ns
ns
1.0
0.8
0.6
0.4
0.2
0.0
Compartment resolution
(held-out marker macro AUPRC)
0.81
0.94
0.93
0.93
one cross-validation fold (n = 5)
mean over folds
single map
(Hein 2025)
annotation mean
(no graph)
averaged
graph
BIONIC
b
***
***
one complex (n = 44)
same complex
mean
Protein-complex resolution
(mean average precision per complex)
0.09
0.33
0.42
single map
(Hein 2025)
averaged
graph
BIONIC

**Supplementary Fig. S7** Deep-learning integration against simple baselines on the 51 input maps of the integrated pan-human map. Four ways of combining the 51 unperturbed human maps are compared on the 9530 proteins of the integrated map: a high-resolution single input map (the HEK293T wild-type map of Hein et al. [5]), the annotation mean (which annotates every map separately by competitive diffusion and averages the resulting compartment probabilities over the maps that measured the protein), the averaged graph, which averages each edge weight of the 51 protein-neighborhood graphs over the maps that measured both proteins and keeps the 50 strongest edges of each protein, as in the graph of the integrated map, and BIONIC [6], the graph-convolutional integration used throughout this work. **(a)** Compartment resolution, measured as the macro-averaged area under the precision–recall curve (AUPRC) of competitive diffusion on held-out curated markers (16 compartments). Each point is one of five stratified cross-validation folds, identical for every strategy, and the black bar is the mean over folds. Brackets give two-sided Mann–Whitney $U$ tests of each strategy against BIONIC over the five folds (single map $P = 0.008$, annotation mean $P = 0.55$, averaged graph $P = 1.00$; ns, not significant; *, $P < 0.05$; **, $P < 0.01$; ***, $P < 0.001$). At the coarse resolution of 16 compartments with marker proteins, BIONIC does not outperform the baseline integration strategies. **(b)** Protein-complex resolution over 44 EBI Complex Portal [7] complexes. Each point represents the mean average precision with which the graph ranks the members of a complex next to each member in turn [8]. This score uses no labels: each complex member serves as a query, all other proteins are ranked by their edge weight to it in the graph, and the average precision of its co-members is recorded. It therefore requires a graph, and the annotation mean, which yields compartment probabilities but no graph, is not shown. Gray lines connect the same complex across strategies. Brackets indicate two-sided Wilcoxon signed-rank tests of each strategy against BIONIC, paired by complex (averaged graph $P = 1.2 \times 10^{-5}$, single map $P = 1.1 \times 10^{-8}$; stars as in **a**).

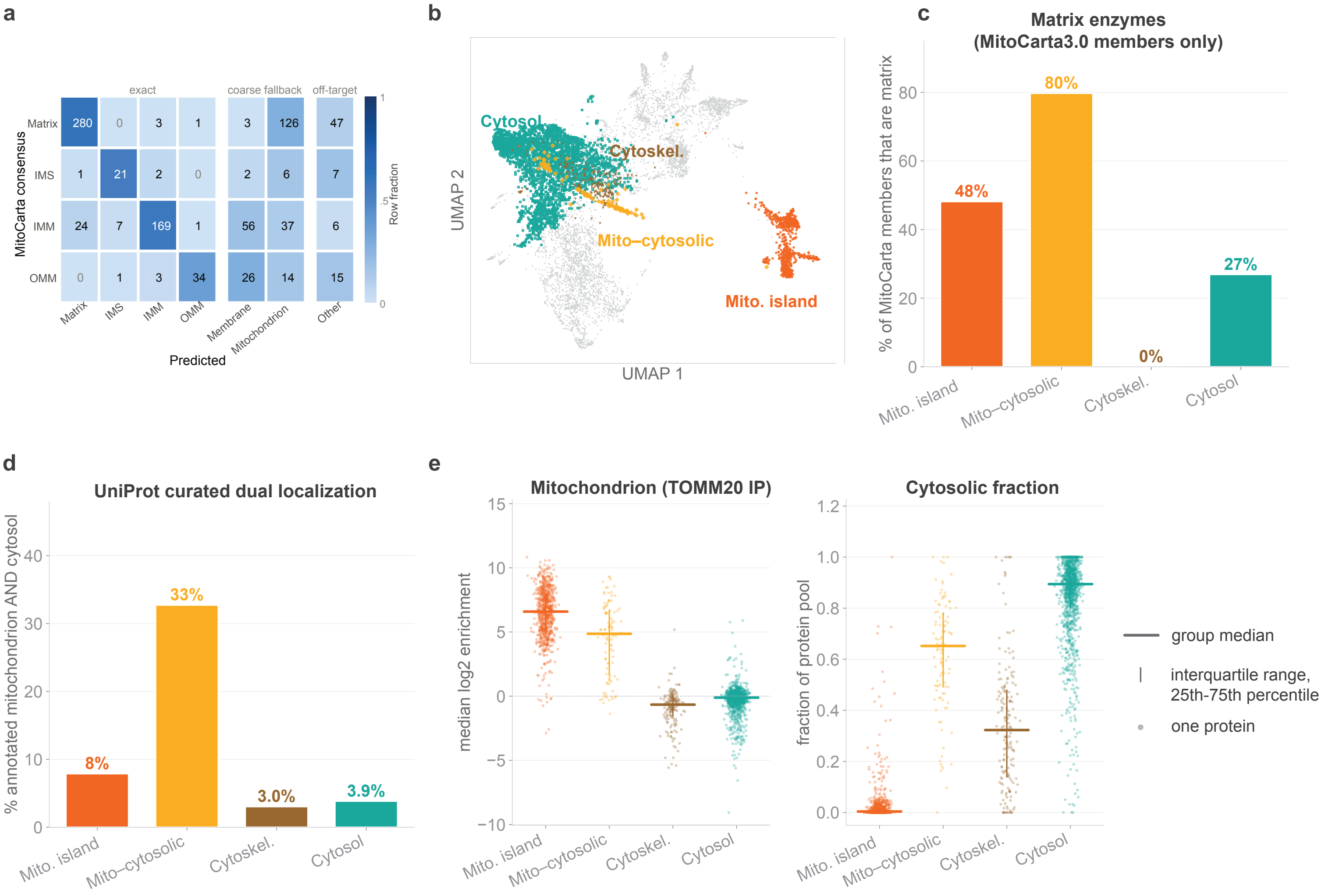
a
exact
coarse fallback
off-target
MitoCarta consensus
Matrix 280 0 3 1 3 126 47
IMS 1 21 2 0 2 6 7
IMM 24 7 169 1 56 37 6
OMM 0 1 3 34 26 14 15
Matrix
IMS
IMM
OMM
Membrane
Mitochondrion
Other
Predicted
Row fraction
1
.5
0
b
Cytosol
Cytoskel.
Mito–cytosolic
Mito. island
UMAP 2
UMAP 1
c
Matrix enzymes
(MitoCarta3.0 members only)
% of MitoCarta members that are matrix
48%
80%
0%
27%
Mito. island
Mito–cytosolic
Cytoskel.
Cytosol
d
UniProt curated dual localization
% annotated mitochondrion AND cytosol
8%
33%
3.0%
3.9%
Mito. island
Mito–cytosolic
Cytoskel.
Cytosol
e
Mitochondrion (TOMM20 IP)
median log2 enrichment
Cytosolic fraction
fraction of protein pool
Mito. island
Mito–cytosolic
Cytoskel.
Cytosol
group median
interquartile range, 25th-75th percentile
one protein

**Supplementary Fig. S8** Characterization of the Mito-cytosolic cluster placed off the mitochondrial island and near the cytosol. **(a)** Row-normalized confusion matrix of the map's fine Gene Ontology cellular-component calls for mitochondrial proteins against the MitoCarta/MitoCoP consensus four-way subcompartment ground truth (matrix, intermembrane space [IMS], inner membrane [IMM], outer membrane [OMM]). Numbers are protein counts. The predicted axis additionally allows correctly coarser fall-back calls (mitochondrial membrane, mitochondrion) and off-mitochondrion. Among proteins that received one of the four subcompartment labels, accuracy was 92.1% (macro-F1 0.89). Most of the other proteins received a correct coarser label. **(b)** The Mito-cytosolic cluster (amber, $n$ = 110) sits away from the mitochondrial island (orange, $n$ = 849) in the UMAP representation of the integrated pan-human map (Fig. **??**d), near the cytoskeleton (brown, $n$ = 230) and the cytosol (teal, $n$ = 2983). Teal marks the 1453-protein cytosol reference group analyzed in **c**–**e** together with 1530 other proteins that the map assigns to the cytoplasm. The remaining 5358 proteins are shown in gray. **(c)** Percentage of each group's MitoCarta3.0 members assigned to the mitochondrial matrix (382 of 793, 59 of 74, 0 of 2, and 7 of 26). **(d)** Percentage of each group whose UniProt subcellular-location annotation matches both a mitochondrial and a cytosolic term (regular-expression match, denominators 849, 110, 230, and 1453). **(e)** Mitochondrial pulldown enrichment (TOMM20 immunoprecipitation, left) and share of each protein found in the cytosolic fraction (right) of a nuclear/organellar/cytosolic fractionation in wild-type HEK293T cells (Hein et al. [5]), for group members quantified there (742, 103, 167, and 1226). Each dot is one protein. The horizontal bar is the group median, and the vertical line is the interquartile range (25th to 75th percentile).

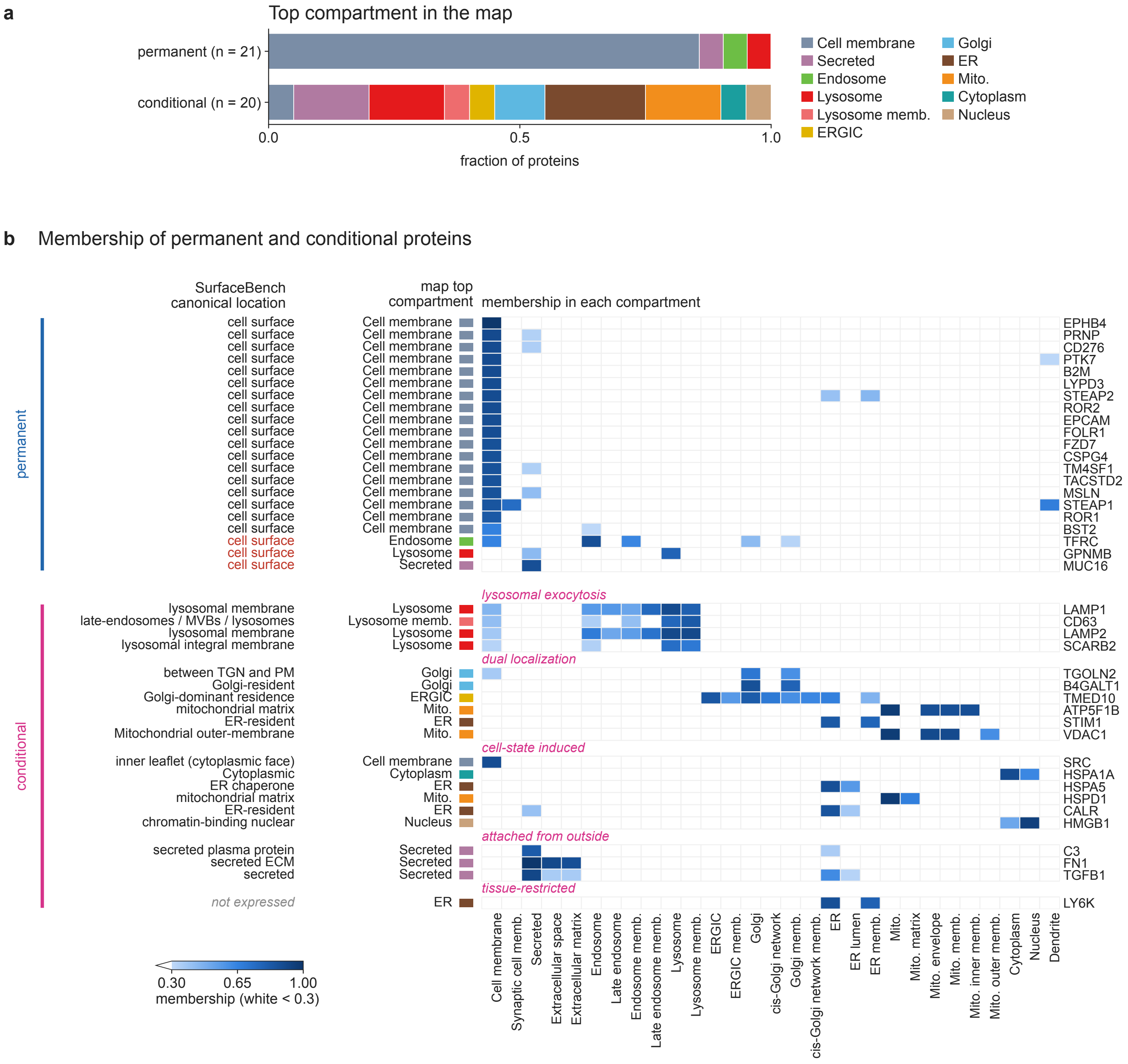

a
Top compartment in the map
permanent (n = 21)
conditional (n = 20)
0.0
0.5
1.0
fraction of proteins
Cell membrane
Secreted
Endosome
Lysosome
Lysosome memb.
ERGIC
Golgi
ER
Mito.
Cytoplasm
Nucleus
b
Membership of permanent and conditional proteins
SurfaceBench canonical location
map top compartment
membership in each compartment
permanent
conditional
cell surface
lysosomal membrane
late-endosomes / MVBs / lysosomes
lysosomal integral membrane
between TGN and PM
Golgi-resident
Golgi-dominant residence
mitochondrial matrix
ER-resident
Mitochondrial outer-membrane
inner leaflet (cytoplasmic face)
Cytoplasmic
ER chaperone
chromatin-binding nuclear
secreted plasma protein
secreted ECM
secreted
not expressed
lysosomal exocytosis
dual localization
cell-state induced
attached from outside
tissue-restricted
EPHB4
PRNP
CD276
PTK7
B2M
LYPD3
STEAP2
ROR2
EPCAM
FOLR1
FZD7
CSPG4
TM4SF1
TACSTD2
MSLN
STEAP1
ROR1
BST2
TFRC
GPNMB
MUC16
LAMP1
CD63
LAMP2
SCARB2
TGOLN2
B4GALT1
TMED10
ATP5F1B
STIM1
VDAC1
SRC
HSPA1A
HSPA5
HSPD1
CALR
HMGB1
C3
FN1
TGFB1
LY6K
Cell membrane
Synaptic cell memb.
Secreted
Extracellular space
Extracellular matrix
Endosome
Late endosome
Endosome memb.
Late endosome memb.
Lysosome
Lysosome memb.
ERGIC
ERGIC memb.
Golgi
cis-Golgi network
Golgi memb.
cis-Golgi network memb.
ER
ER lumen
ER memb.
Mito.
Mito. matrix
Mito. envelope
Mito. memb.
Mito. inner memb.
Mito. outer memb.
Cytoplasm
Nucleus
Dendrite
0.30
0.65
1.00
membership (white < 0.3)

**Supplementary Fig. S9** Placement of SurfaceBench cell-surface proteins in the integrated pan-human map. SurfaceBench [9] labels human proteins by whether the protein is reachable from outside the cell: permanently, conditionally, or not at all. Of its 147 proteins, 80 are in the integrated map (21 permanent, 20 conditional, and 39 not accessible). The figure shows the permanent and conditional proteins, and Supplementary Table **??** lists all three classes. Memberships were computed by independent diffusion of the full UniProt subcellular-location hierarchy over the map's protein-neighborhood graph, after removing all SurfaceBench proteins from the gene sets. **(a)** Top compartment of each permanent and conditional protein. The plasma membrane is the top compartment of 18 of 21 permanent proteins and of 1 of 20 conditional proteins. **(b)** One row per permanent or conditional protein, with conditional proteins grouped by the reason SurfaceBench gives for the conditional label. Left: the canonical location of each conditional protein in the words of its SurfaceBench rationale (hand-coded, Supplementary Table **??**), and the cell surface for every permanent protein, by SurfaceBench's definition. The text is dark where the top compartment of the map (next column) is the same compartment and red where it is not. The two agree for 18 of 21 permanent proteins and for all 19 conditional proteins with a stated baseline location. The rationale of LY6K describes it as normally not expressed (gray). Center: membership in every compartment that reaches 0.5 in at least one row, with memberships below 0.3 shown in white. Right: gene symbol. The median plasma-membrane membership is 0.87 for permanent and 0.12 for conditional proteins (two-sided Mann–Whitney $U$ test, $P = 8.7 \times 10^{-6}$). The 4 lysosomal-exocytosis proteins have a plasma-membrane membership of 0.33–0.45 in addition to their lysosomal membership. Abbreviations used in the figure: ER, endoplasmic reticulum; ERGIC, ER–Golgi intermediate compartment; Mito., mitochondrion; memb., membrane; MVBs, multivesicular bodies; TGN, trans-Golgi network; PM, plasma membrane; ECM, extracellular matrix.

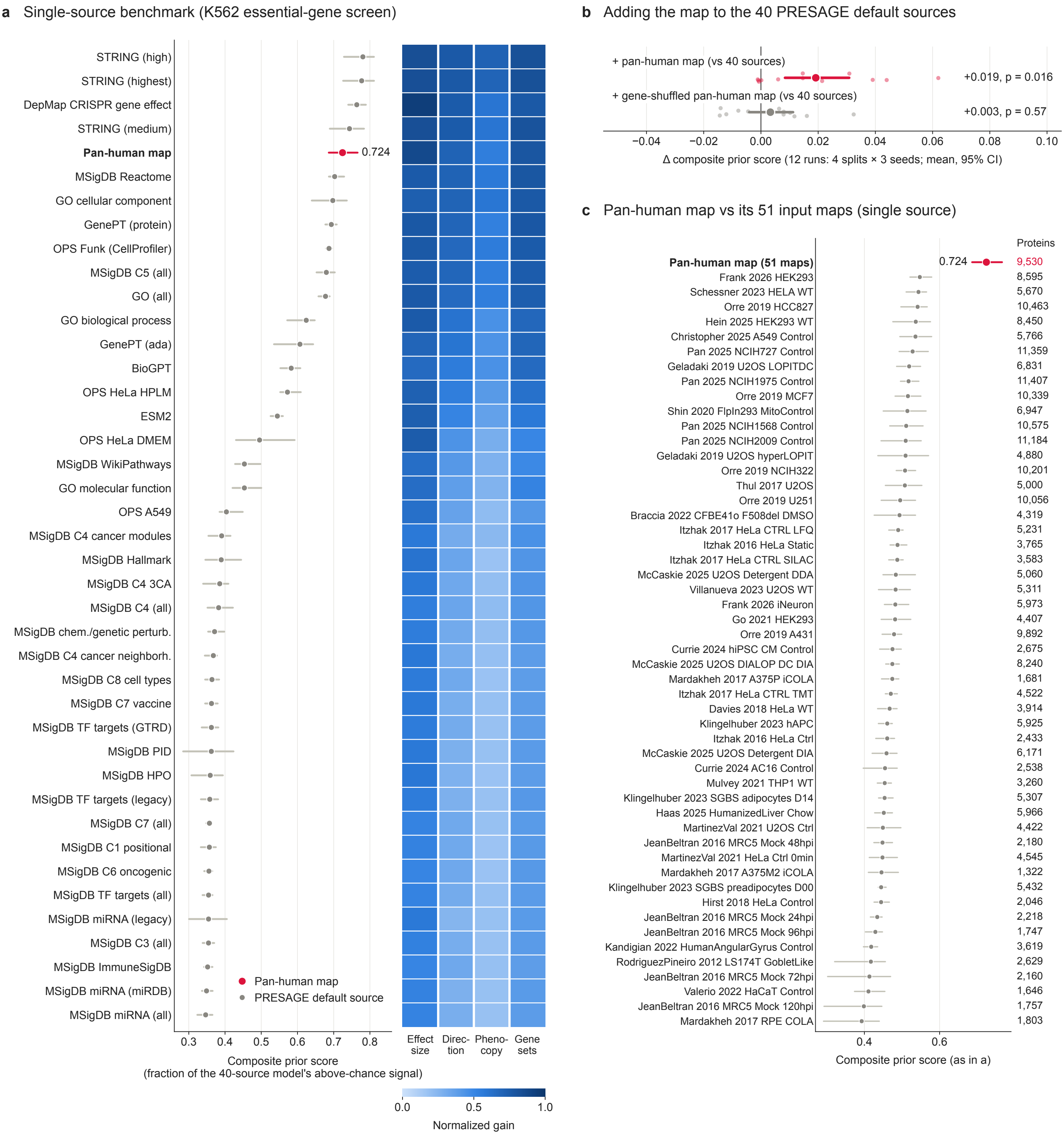
a Single-source benchmark (K562 essential-gene screen)
STRING (high)
STRING (highest)
DepMap CRISPR gene effect
STRING (medium)
Pan-human map
0.724
MSigDB Reactome
GO cellular component
GenePT (protein)
OPS Funk (CellProfiler)
MSigDB C5 (all)
GO (all)
GO biological process
GenePT (ada)
BioGPT
OPS HeLa HPLM
ESM2
OPS HeLa DMEM
MSigDB WikiPathways
GO molecular function
OPS A549
MSigDB C4 cancer modules
MSigDB Hallmark
MSigDB C4 3CA
MSigDB C4 (all)
MSigDB chem./genetic perturb.
MSigDB C4 cancer neighborh.
MSigDB C8 cell types
MSigDB C7 vaccine
MSigDB TF targets (GTRD)
MSigDB PID
MSigDB HPO
MSigDB TF targets (legacy)
MSigDB C7 (all)
MSigDB C1 positional
MSigDB C6 oncogenic
MSigDB TF targets (all)
MSigDB miRNA (legacy)
MSigDB C3 (all)
MSigDB ImmuneSigDB
MSigDB miRNA (miRDB)
MSigDB miRNA (all)
Pan-human map
PRESAGE default source
0.3 0.4 0.5 0.6 0.7 0.8
Composite prior score
(fraction of the 40-source model's above-chance signal)
Effect size
Direc-tion
Pheno-copy
Gene sets
0.0 0.5 1.0
Normalized gain
b Adding the map to the 40 PRESAGE default sources
+ pan-human map (vs 40 sources)
+0.019, p = 0.016
+ gene-shuffled pan-human map (vs 40 sources)
+0.003, p = 0.57
−0.04 −0.02 0.00 0.02 0.04 0.06 0.08 0.10
Δ composite prior score (12 runs: 4 splits × 3 seeds; mean, 95% CI)
c Pan-human map vs its 51 input maps (single source)
Proteins
Pan-human map (51 maps) 0.724 9,530
Frank 2026 HEK293 8,595
Schessner 2023 HELA WT 5,670
Orre 2019 HCC827 10,463
Hein 2025 HEK293 WT 8,450
Christopher 2025 A549 Control 5,766
Pan 2025 NCIH727 Control 11,359
Geladaki 2019 U2OS LOPITDC 6,831
Pan 2025 NCIH1975 Control 11,407
Orre 2019 MCF7 10,339
Shin 2020 FlpIn293 MitoControl 6,947
Pan 2025 NCIH1568 Control 10,575
Pan 2025 NCIH2009 Control 11,184
Geladaki 2019 U2OS hyperLOPIT 4,880
Orre 2019 NCIH322 10,201
Thul 2017 U2OS 5,000
Orre 2019 U251 10,056
Braccia 2022 CFBE41o F508del DMSO 4,319
Itzhak 2017 HeLa CTRL LFQ 5,231
Itzhak 2016 HeLa Static 3,765
Itzhak 2017 HeLa CTRL SILAC 3,583
McCaskie 2025 U2OS Detergent DDA 5,060
Villanueva 2023 U2OS WT 5,311
Frank 2026 iNeuron 5,973
Go 2021 HEK293 4,407
Orre 2019 A431 9,892
Currie 2024 hiPSC CM Control 2,675
McCaskie 2025 U2OS DIALOP DC DIA 8,240
Mardakheh 2017 A375P iCOLA 1,681
Itzhak 2017 HeLa CTRL TMT 4,522
Davies 2018 HeLa WT 3,914
Klingelhuber 2023 hAPC 5,925
Itzhak 2016 HeLa Ctrl 2,433
McCaskie 2025 U2OS Detergent DIA 6,171
Currie 2024 AC16 Control 2,538
Mulvey 2021 THP1 WT 3,260
Klingelhuber 2023 SGBS adipocytes D14 5,307
Haas 2025 HumanizedLiver Chow 5,966
MartinezVal 2021 U2OS Ctrl 4,422
JeanBeltran 2016 MRC5 Mock 48hpi 2,180
MartinezVal 2021 HeLa Ctrl 0min 4,545
Mardakheh 2017 A375M2 iCOLA 1,322
Klingelhuber 2023 SGBS preadipocytes D00 5,432
Hirst 2018 HeLa Control 2,046
JeanBeltran 2016 MRC5 Mock 24hpi 2,218
JeanBeltran 2016 MRC5 Mock 96hpi 1,747
Kandigian 2022 HumanAngularGyrus Control 3,619
RodriguezPineiro 2012 LS174T GobletLike 2,629
JeanBeltran 2016 MRC5 Mock 72hpi 2,160
Valerio 2022 HaCaT Control 1,646
JeanBeltran 2016 MRC5 Mock 120hpi 1,757
Mardakheh 2017 RPE COLA 1,803
0.4 0.6
Composite prior score (as in a)

**Supplementary Fig. S10** The integrated pan-human map is a strong prior for perturbation-response prediction. PRESAGE [10] was trained to predict the transcriptional responses to held-out genetic perturbations of the K562 essential-gene Perturb-seq screen [11] using gene embeddings of prior knowledge. **(a)** Single-source benchmark. PRESAGE trained with one knowledge source at a time: each of its 40 default sources, and the integrated map (crimson). Point, mean over 4 cross-validation splits. Line, range over the splits. The composite prior score is the mean, over four tasks (effect size, response direction, phenocopy, and gene-set phenotypes)(see Methods). The heatmap shows the normalized gain for each task. The scores of all knowledge sources and input maps are given in Supplementary Table **??**. **(b)** Adding the integrated map, or a gene-shuffled version with identical coverage, to the 40 default sources (upstream PRESAGE loader). Change in composite prior score relative to the 40-source model, paired by split and training seed. Small dots, individual runs (4 splits $\times$ 3 seeds); large dot and line, mean and bootstrap 95% confidence interval (CI); $P$, two-sided Wilcoxon signed-rank test over the 12 paired runs. The map raises the composite prior score by 0.019 (95% CI 0.008 to 0.031, 9 of 12 runs improved, $P = 0.016$) and the gene-shuffled map by 0.003 ($P = 0.57$). **(c)** Each of the 51 input subcellular maps used as the only PRESAGE knowledge source, ranked by composite prior score as in **a** (point, mean over 4 cross-validation splits; line, range). Crimson, the integrated map of all 51 maps. Right, number of proteins per map. The best single map scores 0.547 and the median map 0.475, against 0.724 for the integrated map. Abbreviations used in the figure: GO, Gene Ontology; OPS, optical pooled screen; TF, transcription factor.
